# FAST: a fast and scalable factor analysis for spatially aware dimension reduction of multi-section spatial transcriptomics data

**DOI:** 10.1101/2023.07.11.548486

**Authors:** Wei Liu, Xiao Zhang, Xiaoran Chai, Zhenqian Fan, Huazhen Lin, Jinmiao Chen, Lei Sun, Tianwei Yu, Joe Yeong, Jin Liu

## Abstract

Biological techniques for spatially resolved transcriptomics (SRT) have advanced rapidly in both throughput and spatial resolution for a single spatial location. This progress necessitates the development of efficient and scalable spatial dimension reduction methods that can handle large-scale SRT data from multiple sections. Here, we developed FAST as a fast and efficient generalized probabilistic factor analysis for spatially aware dimension reduction, which simultaneously accounts for the count nature of SRT data and extracts a low-dimensional representation of SRT data across multiple sections, while preserving biological effects with consideration of spatial smoothness among nearby locations. Compared with existing methods, FAST uniquely models the count data across multiple sections while using a local spatial dependence with scalable computational complexity. Using both simulated and real datasets, we demonstrated the improved correlation between FAST estimated embeddings and annotated cell/domain types. Furthermore, FAST exhibits remarkable speed, with only FAST being applicable to analyze a mouse embryo Stereo-seq dataset with >2.3 million locations in only 2 hours. More importantly, FAST identified the differential activities of immune-related transcription factors between tumor and non-tumor clusters and also predicted a carcinogenesis factor *CCNH* as the upstream regulator of differentially expressed genes in a breast cancer Xenium dataset.

## Introduction

Spatially resolved transcriptomics (SRT) encompasses a set of breakthrough technologies that enable gene expression profiling with spatial information on tissues. Spatial location information is of paramount significance in comprehending the mechanisms underlying processes such as cell biology [1], tumor biology [2], and developmental biology [3]. Among the many factors that influence the choice of these technologies [4, 5], throughput in profiling and spatial resolution are two of the most important. Technologies based on in situ hybridization (e.g., MERFISH [6], seqFISH [7], and seqFISH+ [8]) and in situ sequencing (ISS) (e.g., FISSEQ [9], and Xenium) provide single-molecule/single-cell resolution, but are for targeted genes that require prior knowledge. While next-generation sequencing (NGS)-based technologies, such as Visium, Slide-seq [10, 11], and Stereo-seq [12], are unbiased and involve high-throughput expression measurements, most do not provide single-cell resolution. The diverse range of SRT technologies has enabled the exploration of intricate transcriptional structure across heterogeneous tissues and will revolutionize the worlds of cell biology and molecular biology and advance our understanding in many areas of biology [13, 14].

For “high-dimensional”, often noisy, expression measurements obtained using SRT technologies, dimensionality reduction is a key step in generating a low-dimensional data representation that enriches biological signals by aggregating gene expression relevant to biological effects [15]. Moreover, transformation in dimension reduction circumvents the curse of dimensionality usually present in “high-dimensional” expression profiles constructed in genomic studies, including those based on SRT [16, 17]. A plethora of dimension reduction methods have been developed, including the most popular method, principal component analysis (PCA) [18], which is routinely used in many software pipelines, such as Seurat [19] and Cell Ranger [20] for single-cell RNA sequencing (scRNA-seq) analysis and BayesSpace [21], SpaGCN [22], and SC-MEB [23] for SRT data analysis. However, PCA does not consider the spatial nature of SRT data in the process of estimating low-dimensional embeddings, omitting the influence of the microenvironment in neighboring locations.

In SRT data analysis, expression patterns among neighboring locations exhibit the “similarity” induced by the shared microenvironment. Recently, SpatialPCA [17] and non-negative spatial factorization (NSF) [24] were proposed for spatially aware dimension reduction using Gaussian-type kernels over spatial locations. To reduce the computational burden, SpatialPCA applies a low-rank approximation, while NSF implements a sparse Gaussian process but is not applicable to spatial locations from multiple sections [24, 25]. With improved spatial resolution, the number of spatial locations profiled increases substantially, while multiple sections are needed to either generate a spatial map or recover the spatiotemporal transcriptomics atlas of a whole organ [12, 26, 27]. More recently, we proposed PRECAST as a method to unify the tasks of dimension reduction, cluster allocation, and embedding alignment for SRT data from multiple sections [28]. However, PRECAST did not model the count nature of SRT data, while it was designed for a unified task of dimension reduction and clustering. When facing downstream tasks other than clustering (e.g., trajectory inference and cell-cell interaction; CCI), PRECAST may not be optimal. Although the computational order for PRECAST is linear to location number *n*, it still takes a few days to complete the analysis if the number of spatial locations goes into the millions due to its unified framework. Ideally, an efficient method that allows for spatial dimension reduction across multiple sections and is capable of capturing information for both biological effects and spatial correlation structure is also required. This method should also account for the count nature of SRT data, be applicable to multiple downstream tasks, and be scalable to millions of spatial locations.

To address the limitations of existing methods and facilitate scalability, we proposed FAST as a generalized probabilistic factor analysis model for spatial transcriptomics that efficiently estimates embeddings across multiple sections intrinsic to biological effects, taking into account local expression similarities induced by the shared microenvironment. Uniquely, FAST explicitly allows simultaneous spatially aware dimensionality reduction across multiple sections while modeling the count nature of many existing SRT datasets. Moreover, scalability is promoted by modeling of local spatial dependence using a conditional autoregressive component with an improved computational complexity linear to *n*, facilitating its applicability to the analysis of multi-section high-resolution SRT data. FAST is versatile in analyzing various multi-section SRT datasets obtained from distinct spatial transcriptomics technologies and tissue structures.

## Results

### Spatial dimension reduction using FAST

Similar to scRNA-seq analysis, dimension reduction is an essential step for many downstream analyses (Fig. 1a, left panel). We describe FAST in the “Methods” section and provide its technical details in the Supplementary Notes. Briefly, FAST is a generalized probabilistic factor analysis for spatially aware dimension reduction across multi-section spatial transcriptomics data with millions of spatial locations (Fig. 1b). Taking the normalized/count gene expression matrices from multiple sections as input data, FAST factorizes the expression matrices into factor matrices with a shared loading matrix, while assuming a conditional autoregressive (CAR) component for factors from each section (Fig. 1a, right panel). Instead of applying a global kernel for all spatial locations, FAST models local spatial dependence induced by neighboring microenvironments using CAR components. We showed that this consideration in FAST not only reduces the computational complexity in linear form to *n*, scalable to millions of spatial locations, but also estimates meaningful embeddings that correlate more closely with biological effects. Subsequently, by performing an integrative analysis to remove batch effects via iSC-MEB, the aligned embeddings can be paired with many existing software/tools developed in scRNA-seq studies to enhance the effectiveness of downstream analyses in SRT studies (Fig. 1c). FAST is implemented as an R package available at https://github.com/feiyoung/FAST.

**Figure 1:**
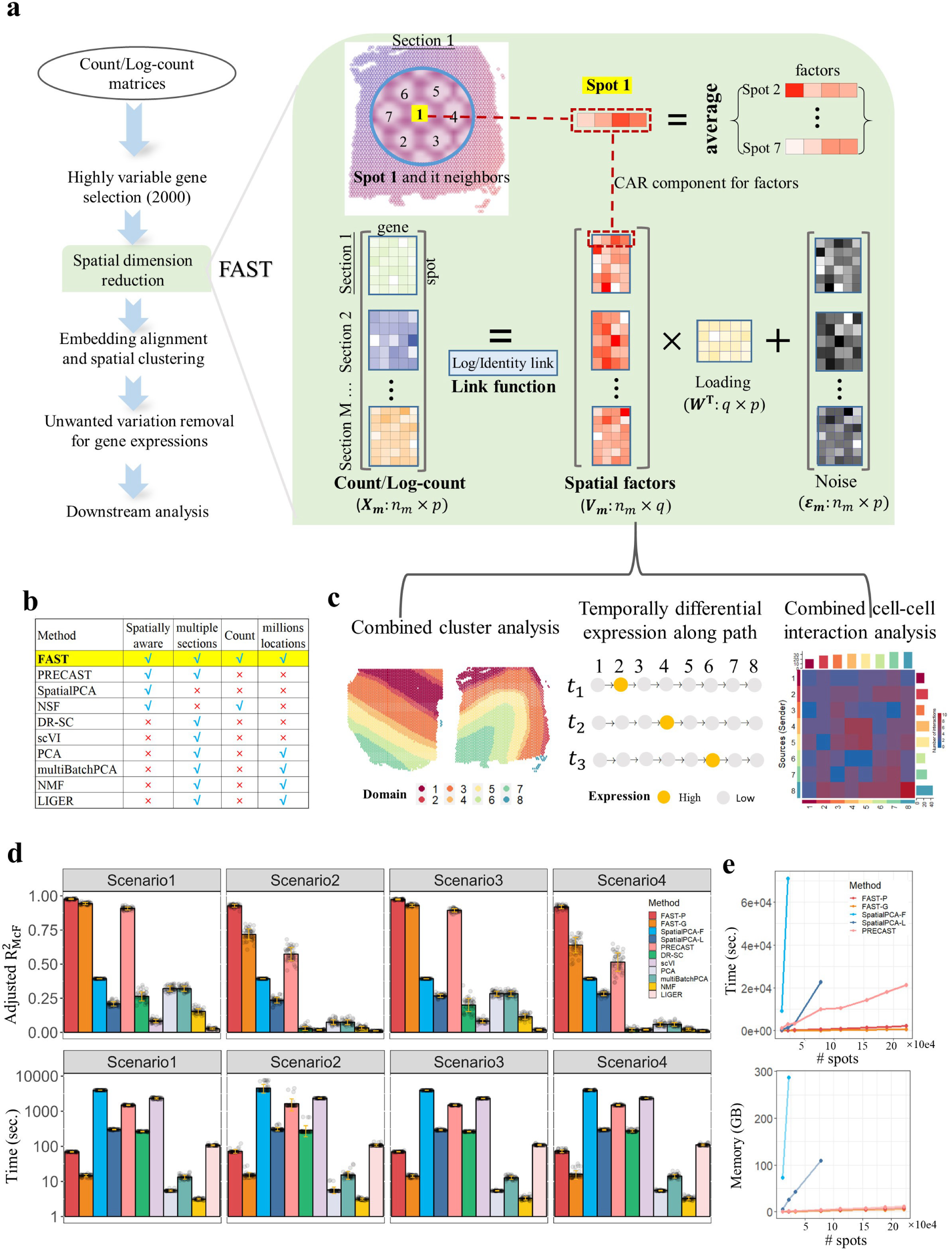
Schematic overview of FAST and simulation results. (a) Left panel: Workflow of multi-section SRT data analysis comprising the following steps: count or log-count as input, highly variable gene selection, spatial dimension reduction based on FAST, embedding alignment and spatial clustering, unwanted variation removal for combing gene expressions from multi-sections, and a series of downstream analyses. Right panel: FAST is a powerful generalized probabilistic factor model that efficiently estimates embeddings while incorporating spatial smoothness across multiple expression matrices. It has the flexibility to utilize either count or log-normalized gene expression matrices as input via log or identity link, enabling it to project all spots onto a shared low-dimensional space. This capability greatly enhances downstream analyses by effectively utilizing information from all sections. (b) A comparative analysis of four aspects: spatial awareness, multi-section applicability, count matrix utilization, and scalability to millions of locations in FAST and other dimension reduction methods. (c) Representative FAST downstream analyses: combined clustering analysis, temporally differential expression analysis along differentiation path, and combined CCI analysis. (d) In the simulations, we conducted tests in four different scenarios to assess the effectiveness of FAST by varying the parameters of batch effects and biological effects between low and high values. The four scenarios tested were as follows: Scenario 1 (batch effect=low, biological effect=high), Scenario 2 (batch effect=low, biological effect=low), Scenario 3 (batch effect=high, biological effect=high), and Scenario 4 (batch effect=high, biological effect=low). Top panel: Bar plots of the adjusted 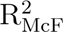 values for FAST and other methods. Bottom panel: Bar plots of the running time (seconds) for FAST and other methods. (e) Comparison of computational cost in terms of running time and memory usage for FAST, SpatialPCA and PRECAST with regard to the number of spots. Note that the lines of both versions of SpatialPCA were truncated due to their inability to handle data of such scale.

### Validation using simulated data

We conducted extensive simulations to assess the performance of FAST and compare it to several other methods (Fig. 1d; Supplementary Fig. S1). The methods compared included a range of spatially aware and non-spatially aware dimension reduction methods such as SpatialPCA [17], PRECAST [28], DR-SC [29], scVI [30], PCA, multiBatchPCA [31], NMF and LIGER [32]. Among these methods, SpatialPCA provides a full-rank, SpatialPCA-F, and a low-rank approximation version, SpatialPCA-L. We performed simulations using three sections from the human dorsolateral prefrontal cortex (DLPFC) Visium dataset with eight predefined spatial domains [33], comprising layers 1–6, whiter matter (WM), and unknown cells (Unknown). Supplementary Fig. S1a provides a visual representation of the spatial distribution of the eight domains for each section. The simulation details are provided in the “Methods” section. Briefly, we considered four simulation scenarios in combinations of high and low values for both biological effects and batch effects, reflecting both shared and section-specific effects.

To quantify the performance of the embedding estimation, we calculated the adjusted McFadden’s pseudo R^2^ (adjusted R^2^) between the estimated embeddings and the true labels for each section. As shown in Fig. 1d (top panel), FAST outperformed other spatially aware methods such as PRECAST and SpatialPCA, and performed much better than the non-spatially aware methods such as PCA and scVI, across all scenarios. In general, spatially aware methods performed better than the non-spatially aware methods. With biological effects between domains varying from strong to weak, the performance of all methods decreased, but the Poisson version of FAST, FAST-P, was the least sensitive compared with the other methods. However, all methods suffered slightly from the increased batch effects in each section. Despite its inferiority to FAST-P, the Gaussian version of FAST, FAST-G, was more computationally efficient. In all scenarios, both versions of FAST were more computationally efficient than other spatially aware methods such as SpatialPCA and PRECAST (Fig. 1d, bottom panel). By applying iSC-MEB, we aligned embeddings estimated using different methods to detect spatial domains. Not surprisingly, FAST-P achieved the highest adjusted Rand index (ARI) and normalized mutual information (NMI) across all scenarios (Supplementary Fig. S1b).

To evaluate the scalability of FAST, we compared its computational efficiency by varying the number of spatial locations analyzed while fixing the number of genes to 2000 (Fig. 1e). Clearly, both FAST-P and FAST-G demonstrated superior computational efficiency. The computational complexity orders for both FAST and PRECAST were linear to the number of spatial locations while they both used less memory. In our study, SpatialPCA-F required approximately 19 hours and 287 GB of memory to analyze a dataset with around 20,000 locations, but experienced breakdowns when reaching 40,000 locations. Meanwhile, SpatialPCA-L took approximately 6 hours and 108 GB of memory to analyze a dataset with 78,000 locations, but experienced breakdowns at 80,000 locations. However, FAST-P and FAST-G exhibited impressive scalability. FAST-P was able to analyze 200,000 locations in just 30 minutes with 7 GB of memory usage, while FAST-G achieved the same task in only 10 minutes with 5 GB of memory usage.

### Application to the human dorsolateral prefrontal cortex Visium dataset

We applied FAST and the other methods to the analysis of four published datasets obtained via either Visium, Xenium, or Stereo-seq technologies (see “Methods”). The four datasets included a DLPFC dataset [33] and a hepatocellular carcinoma dataset [34] generated using 10*×* Visium, a breast cancer dataset [35] generated using 10*×* Xenium, and a mouse embryo dataset [12] generated using Stereo-seq. First, we examined the dimension reduction performance of FAST in comparison with SpatialPCA, PRECAST, DR-SC, scVI, PCA, multiBatchPCA, NMF, and LIGER, followed by the application of iSC-MEB to perform clustering analysis and align embeddings from multiple sections. For some or all four datasets, downstream analyses were also performed, including DE analysis, CCI analysis, cell-type deconvolution, and somatic mutation.

To quantify the ability of FAST to outperform existing methods for dimension reduction, we first analyzed the LIBD human DLPFC dataset generated using 10*×* Visium [33], which contained a total of 47,681 spatial locations across 12 tissue sections from three donors. Taking the manual annotations for the tissue layers based on the cytoarchitecture provided by the original study as ground truth, we were able to evaluate the performance of the dimension reduction. For this purpose, we used the adjusted McFadden’s pseudo R^2^ (adjusted 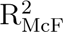) between the estimated embeddings and manual annotations for each section. As shown in Fig. 2a, FAST achieved the highest adjusted 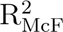 while it was 50 times faster and required only 3% memory usage of SpatialPCA with a low-rank approximation, SpatialPCA-L. In detail, FAST-P required 661 seconds with 2 GB memory usage to complete the analysis compared with the 37,172 seconds and 80 GB memory usage required by SpatialPCA-L. After obtaining estimated embeddings from a variety of dimension reduction methods, we performed integrative clustering analysis to detect spatial domains by applying iSC-MEB. For each method, we summarized the aligned embeddings using three components extracted from UMAP. We then visualized the resulting UMAP components using red/green/blue (RGB) colors in the RGB plot (Fig. 2b, upper right panel; Supplementary Fig. S2), accompanied by the corresponding spatial heatmap of cluster assignment (Fig. 2b lower right panel; Supplementary Fig. S3) and the tSNE plot (Supplementary Fig. S4). The results using FAST embeddings exhibited stronger laminar patterns while presenting a harmonious blending of locations from various sections. These findings illustrated the utility of FAST for estimating embeddings of high-dimensional expression profiles among spatial locations. To evaluate the clustering accuracy of the methods, we used both ARI and NMI. As shown in Fig. 2c, FAST achieved the highest ARI and NMI, with median ARIs for FAST-P, FAST-G, SpatialPCA-L, for PRECAST, DR-SC, scVI, PCA, multiBatchPCA and NMF of 0.56, 0.52, 0.20, 0.45, 0.39, 0.45, 0.42, 0.42 and 0.41, respectively.

**Figure 2:**
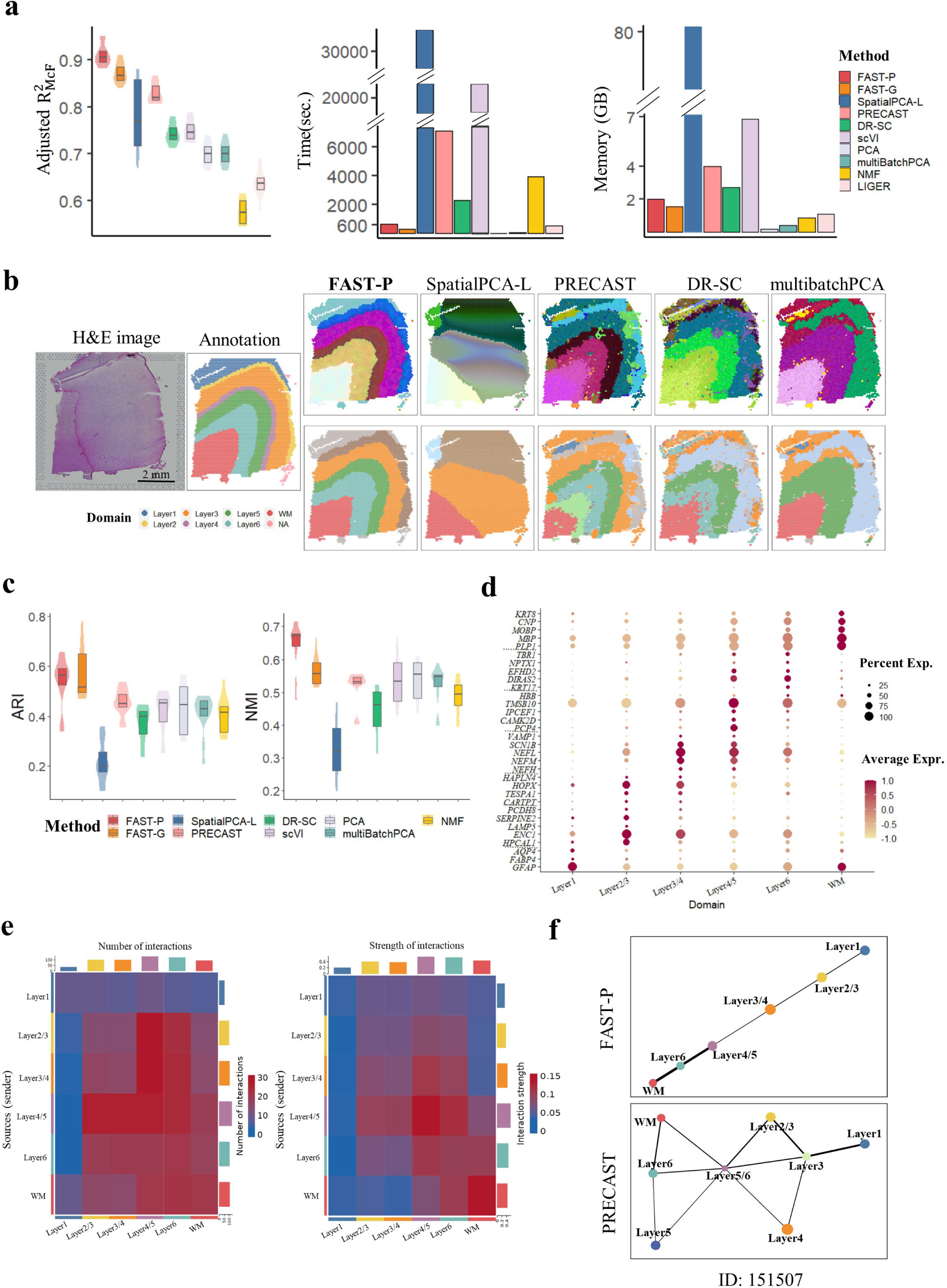
Analysis of human DLPFC data (*n* = 47, 680 locations over 12 tissue sections). (a): Box/violin plot of adjusted 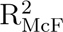 values for FAST and other methods, and bar plots of running time (seconds) and memory usage (GB) for FAST and other methods. In the boxplot, the center line and box lines denote the median, upper, and lower quartiles, respectively. (b): Left panel: H&E image and manual annotation of sample ID 151674. Top panel: UMAP RGB plots of sample ID 151674 for FAST, SpatialPCA-L, DR-SC, mutibatchPCA, and NMF. Bottom panel: Clustering assignment heatmaps for these five methods. (c): Box/violin plots of ARI/NMI values for FAST and other methods. In the boxplot, the center line and box lines denote the median, upper, and lower quartiles, respectively. (d): Dot plot of the normalized expression levels aligned across 12 sections for the marker genes of the layers detected by FAST-P. (e) Combined cell-cell interaction analysis using the normalized expression levels aligned across 12 sections and layer labels obtained by FAST-P. Heatmap of the number/strength of interactions among the different layers. (f) PAGA trajectory for sample ID 151507 using the embeddings and layer labels obtained by FAST-P and PRECAST, respectively.

A key feature of FAST is its ability to estimate low-dimensional embeddings for spatial locations across multiple sections, facilitating many downstream analyses requiring cross-section embedding alignment such as DE analysis and CCI analysis. First, we performed DE analysis for all 12 sections by removing unwanted variations between expression profiles in multiple sections (see “Methods”). In total, we detected 1069 differentially expressed genes (DEGs) with adjusted *p*-values *<* 0.001 among the eight domains identified by FAST-P, with 163 genes specific to Domain 1, which corresponds to layer 1 (Supplementary Data 1). A dot plot of normalized expression aligned across 12 sections showed good separation of the DEGs across the detected spatial domains, many of which are human layer-specific markers such as *PCP4* [36], *DIRAS2* [37], *MBP* [33], and *MOBP* [33] (Fig. 2d).

Next, we performed CCI analysis for all spatial locations across multiple sections using CellChat [38] (see “Supplementary Notes”). We observed strong spatial patterns in both the number and the strength of interactions (Fig. 2e), with WM showing substantial interactions with other layers, consistent with its crucial role in transmitting messages between different regions of the brain [39]. We further examined the signaling pathways enriched in each layer during cell-cell communications and found that WM had the highest score in both incoming and outgoing signals, sending electrical signals across different layers via 11 signaling pathways (Supplementary Fig. S5). These pathways included somatostatin, a known presynaptic modulator of glutamatergic signaling in the central nervous system, and the neuregulins signaling pathway, which is crucial in various aspects of the nervous system, including the development, maintenance, and repair processes [40]. Moreover, an estimated PAGA graph [41] generated using FAST-P embeddings demonstrated an almost linear development trajectory from WM to layer 1 in many of the 12 DLPFC sections (Fig. 2f, top panel; Supplementary Fig. S6). But, when employing PRECAST embeddings for the PAGA graph, the resulting topology showed a messy structure (Fig. 2f, bottom panel; Supplementary Fig. S7).

### Application to the breast cancer Xenium dataset

We further applied FAST and the other methods to analyze two breast cancer sections generated using 10*×* Xenium [35] and containing a total of 72,651 spatial locations with expression profiling for 313 genes and their corresponding H&E images (Fig. 3a). In detail, we estimated 15- dimensional embeddings using FAST and the other methods followed by the application of iSC-MEB for clustering analysis and alignment of the embeddings for the two breast cancer sections. For each method, we summarized the aligned embeddings using three components extracted from UMAP. We then visualized the resulting UMAP components using RGB colors in the RGB plot (Fig. 3b and Supplementary Fig. S8). As shown in Fig. 3c and Supplementary Fig. S9, we detected a total of 17 domains in two sections using FAST-P, with the analysis completed in 133 seconds using 0.33 GB memory usage while SpatialPCA-L required 105,417 seconds and 11.31 GB memory usage (Fig. 3d and Supplementary Fig. S10). The estimated cluster proportions matched well in two adjacent sections (Fig. 3e).

**Figure 3:**
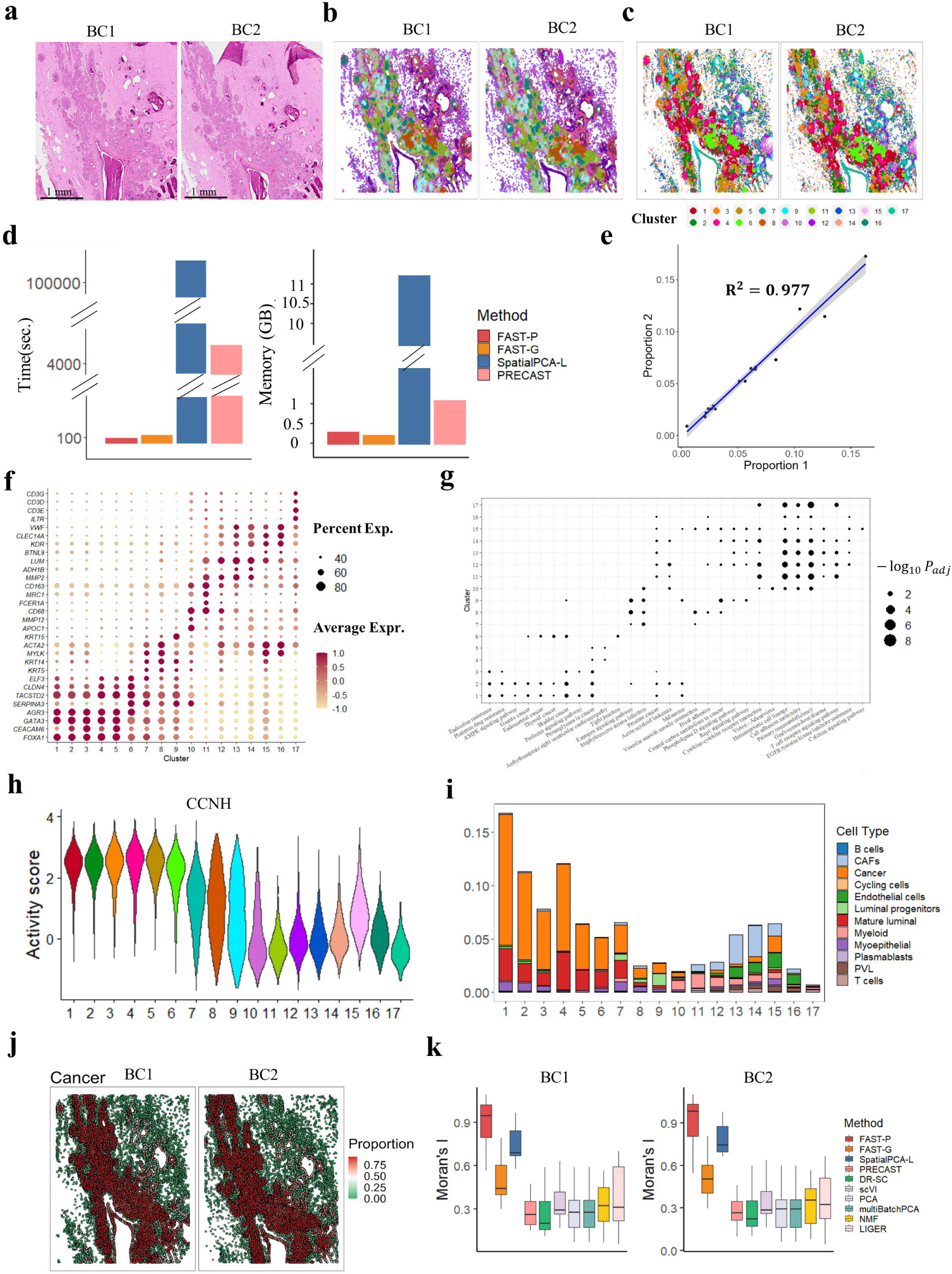
Analysis of human breast cancer Xenium data (*n* = 72, 651 locations over two tissue sections). (a): H&E images of the two sections. (b): UMAP RGB plots of these two sections for FAST-P. (c): Clustering assignment heatmaps of these two sections for the 17 clusters detected by FAST-P. (d): Bar plots of running time (seconds) and memory usage (GB) for FAST, SpatialPCA-L and PRECAST. (e): Scatter plot of the proportions of the 17 clusters in sections BC1 and BC2, with the fitted smoothing line and confidence band determined by the linear regression. (f) Dot plot of the normalized expression levels aligned across two sections for the marker genes of the 17 clusters detected by FAST-P. (g) Dot plot of the significant KEGG pathways of the marker genes for the 17 clusters identified by FAST-P. (h) Violin plot of the activity score for the regulatory protein *CCNH* in the 17 clusters. (i) Percentage of different cell types in each domain detected by FAST-P with scaling to the summation of all cell types across all domains equal to 100%. CAFs: cancer-associated fibroblasts; PVL: perivascular-like cells. (j) Spatial heatmap of deconvoluted cell proportions in cancer cells. (k) Boxplots of the Moran’s I for the 15-dimensional embeddings obtained by FAST and other methods. In the boxplot, the center line, box lines and whiskers represent the median, upper, and lower quartiles, and the 1.5-times interquartile range, respectively.

After removing unwanted variations in expression profiles, we performed DE analysis for the two sections and detected a total of 1416 DEGs with adjusted *p*-values *<* 0.001 in all 17 of the spatial domains detected by FAST-P (Supplementary Data 2), including marker genes for breast cancer such as *TACSTD2* and *FOXA1* [35]. The dot plot of average expression levels across all 17 of the detected domains showed that many of the highly expressed genes specific to Domains 1–6 were marker genes for breast cancer (Fig. 3f). Further detailed examination revealed many marker genes for other cell types present in breast cancer, including *MYLK* [42] (Cluster 7, myoepithelial cells), *KRT5, KRT14* [43] (Cluster 8, basal cells), *KRT15* [44] (Cluster 9, luminal progenitor cells), *APOC1* [45] (Cluster 10, macrophages), *FCER1A, MRC1* [46] (Clusters 11 and 12, dendritic cells), *ADH1B* [47], *MMP2* [48] (Clusters 13–16, fibroblasts), and *IL7R, CD3D* [49] (Cluster 17, T cells). Further KEGG enrichment analysis revealed that genes specific to Clusters 1–6 were significantly enriched in many cancer-related pathways (Fig. 3g and Supplementary Fig. S11). For example, Clusters 1–3 were enriched in endocrine resistance, platinum drug resistance, and some cancer-related pathways. These findings provide valuable insights into the molecular mechanisms underlying cancer development and progression, and have important implications for the development of targeted therapies.

To investigate the transcription factors that regulate gene expression, we performed aberrant protein activity analysis for the two sections. As shown in Fig. 3h and Supplementary Fig. S12, the activities of immune-related transcription factors, such as *IL16*, *CD86*, *TNFRSF4*, and *POU2AF1*, in non-tumor regions (Clusters 10–17) were significantly higher than those in tumor regions (Clusters 1–6). Among these transcription factors, the activity levels of *POU2AF1* and *TNFRSF4* were much higher in the T cell cluster (Cluster 17). This observation is consistent with the active transcription of *POU2AF1* [50] and the function of *TNFRSF4* as a tumor necrosis factor [51] in activated T cells. Although *CCNH* was not measured among the 313 gene profiles, we still predicted a higher activity level for *CCNH* in tumor regions (Clusters 1–6). Further deconvolution analysis revealed that Clusters 1–6 were enriched for cancer cells, which was consistent with the H&E images (Fig. 3i&j and Supplementary Fig. S13&14). Moran’s I values of the top 15 components from FAST-P were on average 155% higher than those with no spatial consideration in estimating embeddings and 17% higher than those from SpatialPCA-L (Fig. 3k).

### Application to the hepatocellular carcinoma Visium dataset

To study mutational patterns in tumor and tumor-adjacent tissues, we also analyzed four sections of the HCC dataset [34] generated using the 10*×* Visium platform, with two sections from tumors (HCC1 and HCC2) and two from tumor-adjacent tissues (HCC3 and HCC4) collected from a patient with HCC, containing a total of 9813 spatial locations with a median number of 3635 genes per location. Fig. 4a shows a histology image (top panel), accompanied by manual annotations made by a pathologist for tumor/normal epithelium (TNE) and stroma regions, and the spatial heatmap of nine spatial domains detected by FAST-P (bottom panel). By performing integrative clustering via iSC-MEB, we aligned embeddings across multiple sections and allocated domain labels for each location. The aligned embeddings were visualized using two components of tSNE for each method (Fig. 4b; Supplementary Fig. S15). Both FAST and PRECAST achieved better data integration performance, while the computational speed for FAST was much faster than that for PRECAST and SpatialPCA (Fig. 4c; Supplementary Fig. S16). For each method, we summarized the aligned embeddings using three components extracted from UMAP. We then visualized the resulting UMAP components using RGB colors in the RGB plot (Supplementary Fig. S17), with the one for FAST showing clear segregation of the TNE and stroma.

**Figure 4:**
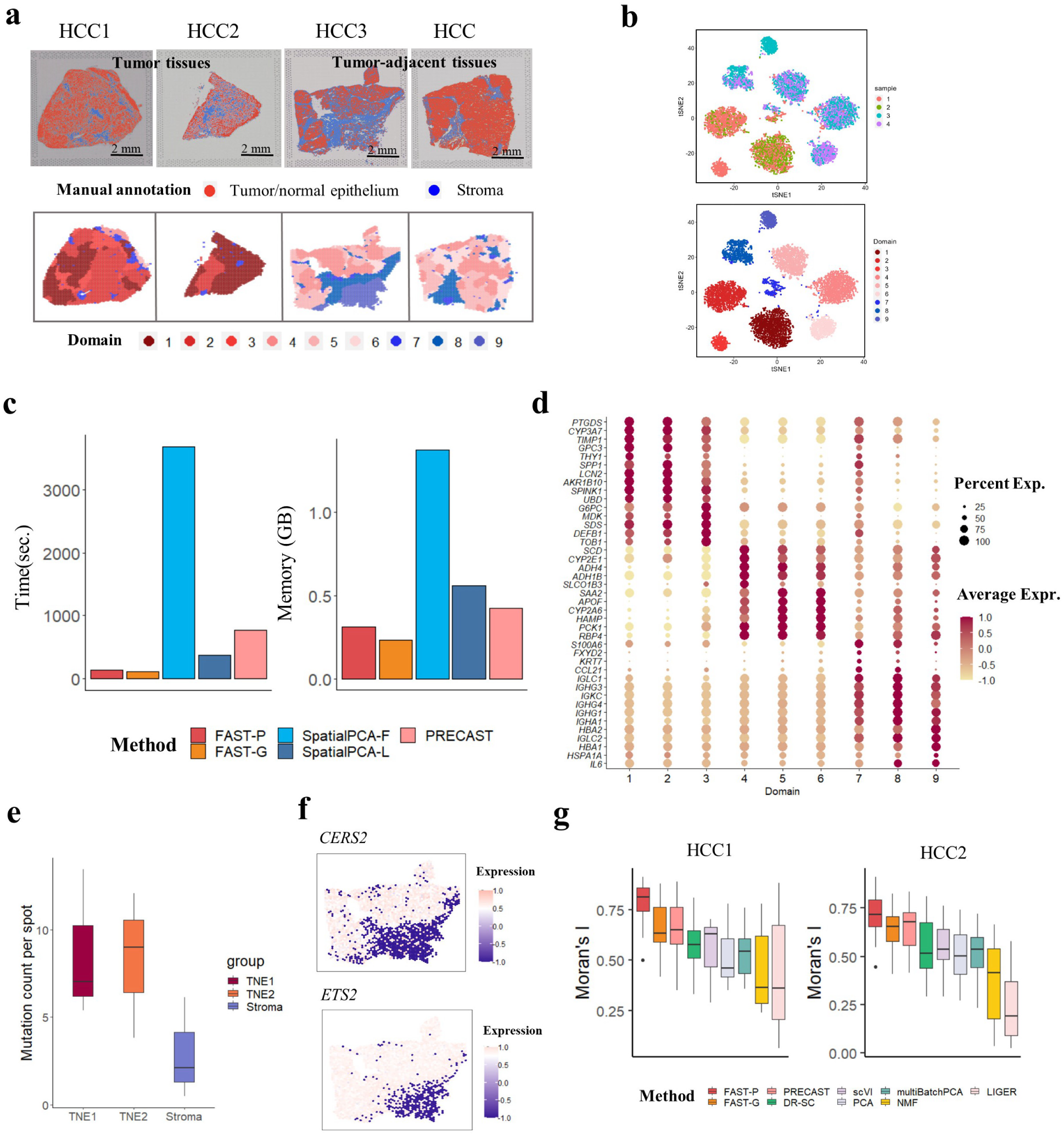
Analysis of human hepatocellular carcinoma Visium data (*n* = 9, 813 locations over four tissue sections). (a): Top panel: H&E images from four tissue sections; Bottom panel: Clustering assignment heatmaps for four tissue sections by FAST-P. (b): tSNE plots of the aligned embeddings of FAST-P. (c): Bar plots of running time (seconds) and Memory usage (GB) for FAST, SpatialPCA and PRECAST. (d): Dot plot of the normalized expression levels aligned across four sections for the marker genes of the nine clusters detected by FAST-P. (e): Box plot of mutation count per spot for TNE1 (Domains 1-3), TNE2 (Domains 4-6) and stroma (Doamins 7-9). (f): Spatial heatmap of the expression levels of *CERS2* and *ETS2* genes in HCC3, corresponding to the top two SNPs with somatic mutations. (g): Boxplots of Moran’s I for the 15-dimensional embeddings obtained by FAST and other methods. We did not plot Moran’s I for SpatialPCA because SpatialPCA produced almost identical embedding for each spot and had a Moran’s I value of 1. In the boxplots, the center line, box lines and whiskers represent the median, upper, and lower quartiles, and 1.5-times interquartile range, respectively.

After removing unwanted variations in the expression profiles, we performed DE analysis and detected a total of 2745 DEGs with adjusted *p*-values *<* 0.001 in all nine spatial domains detected by FAST-P (Supplementary Data 3). These DEGs included HCC marker genes such as *GPC3* and *CYP2A6* [52, 53]. The dot plot of the average expression levels across all nine detected domains revealed many genes specific to tumor regions in tumor/tumor-adjacent tissues and immune regions (Fig. 4d). In detail, Domains 1–3 comprised tumor regions present only in tumor tissues, among which *CYP2A6* was downregulated, while *THY1* [54], *GPC3*, and *CYP3A7* [55] were upregulated. Domains 4–6 comprised tumor regions present primarily in tumor-adjacent tissues, among which *SCD* [56] and *PCK1* [57] were upregulated. Domains 7-9 were enriched in many immune-related marker genes, including *IGLC1*, *IGHG3*, and *IGKC* [58].

To study the somatic mutation landscapes, we performed location-level mutation detection in the spatial transcriptomics data for all four sections (see “Supplementary Notes”). As shown in Fig. 4e, we detected more somatic mutations in TNE (Domains 1–6) than in stroma (Domains 7–9). In detail, we identified four genes containing the top four SNPs with somatic mutations and visualized their spatial expression across four HCC sections (Supplementary Fig. S18–20), in which *CESR2* and *ETS2* showed higher expression in TNE than in stroma (Fig. 4f). The Moran’s I values of the top 15 components of FAST-P were, on average, 19% higher than those of the other methods (Fig. 4g; Supplementary Fig. S21). In total, FAST-P completed the analysis in 110 seconds and used 0.31 GB memory, while SpatialPCA-F required 3680 seconds and 1.37 GB of memory usage (Fig. 4c).

### Application to the mouse embryo Stereo-seq dataset

The Stereo-seq technology was recently developed for high-resolution spatial transcriptomics using 0.22-*µ*m-diameter DNA nanoball (DNB)-patterned arrays [12]. To demonstrate its scalability, we compared the application of FAST with other methods to learn a low-dimensional representation of all spatial locations from a large-scale spatial transcriptomics study on C57BL/6 mouse embryos. In the analysis, we first binned the data as 50bin, with manual annotations based on the expression of marker genes [12]. In total, we analyzed Stereo-seq data from 26 sagittal sections of mouse embryos collected at one-day intervals from E12.5 to E16.5, containing an average of 27,295 genes over a total of 2,323,044 spatial locations. Among all the methods capable of being used for spatial dimension reduction, only FAST was applicable to the analysis of this dataset, while SpatialPCA, PRECAST, and DR-SC were unable to analyze data at this scale. For the analysis of these data, we exclusively utilized FAST-G due to its high computational efficiency compared with the other methods. FAST-G completed the analysis in approximately 2 hours, requiring 94 GB of memory.

First, we showed that FAST achieved the highest adjusted 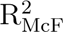 (Fig. 5a), with a median of 0.79 for FAST-G, 0.71 for PCA and 0.67 for LIGER. To assess the accuracy of clustering, we conducted integrative clustering analysis using iSC-MEB to detect spatial clusters across sections. As shown in Fig. 5b, FAST-G achieved the highest ARI and NMI. FAST-G also provided a more accurate representation of the annotated spatial domains compared to alternative methods (Fig. 5c, left panel). Moreover, our summary of the aligned embeddings using three components from UMAP and their visualization as RGB colors in the RGB plot (Fig. 5c, right panel) further highlighted the superior performance of FAST.

**Figure 5:**
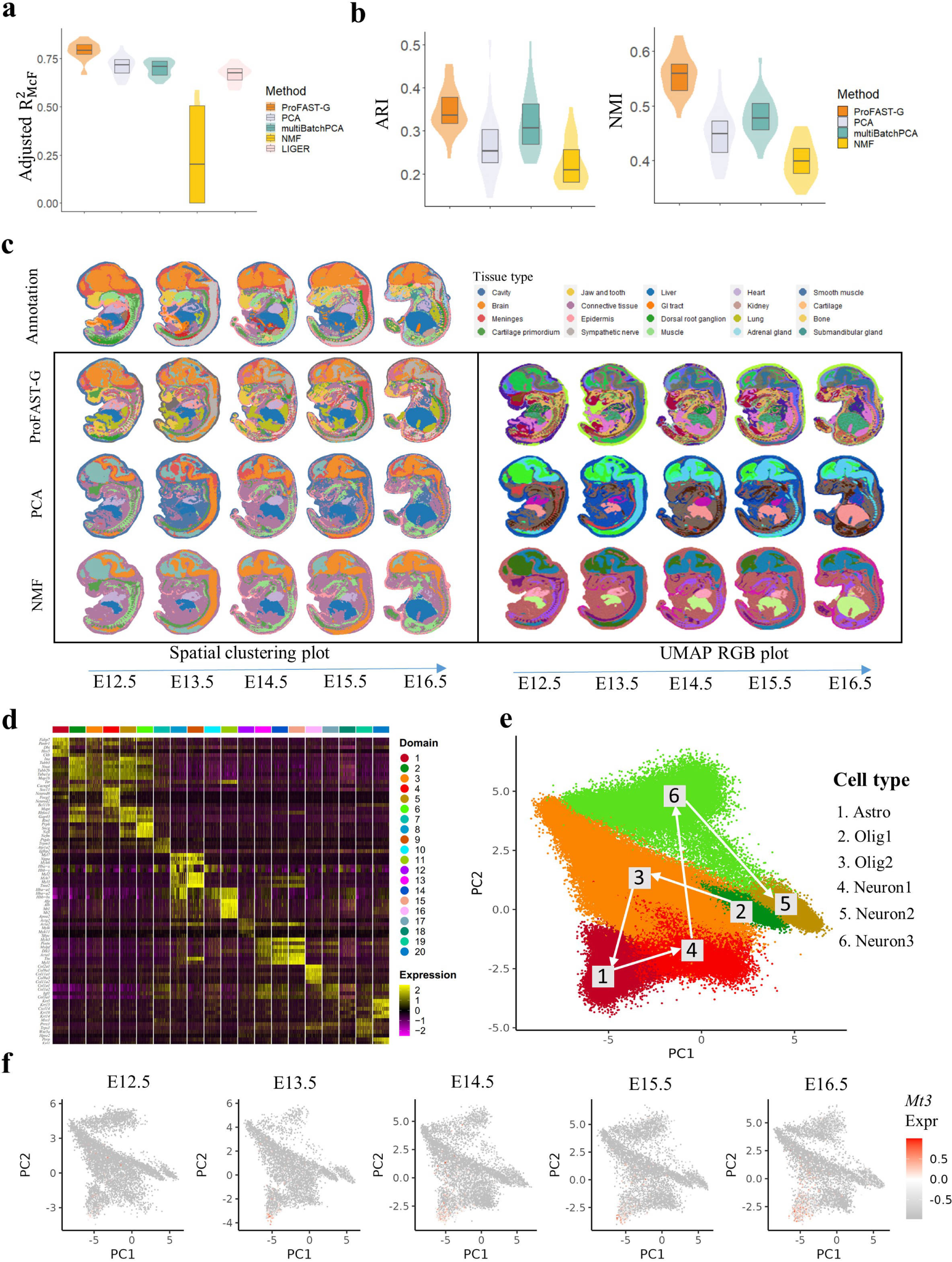
Analysis of mouse embryo Stereo-seq data (*n* = 2, 323, 044 locations over 26 tissue sections). (a): Box/violin plot of adjusted 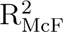 values for FAST-G and four other methods. (b): Box/violin plot of ARI/NMI values for iSC-MEB clustering based on the embeddings obtained by FAST-G and three other methods. (c): Left panel: Clustering assignment heatmaps for five sections (sample IDs: 1, 7, 11, 18, 23) from different embryo days (the first sample in E12.5, E13.5, E14.5, E15.5 and E16.5, respectively) by manual annotation, FAST-G, PCA and NMF. Right panel: UMAP RGB plots for these sections based on the aligned embeddings from FAST-G, PCA and NMF. (d): Heatmap of differentially expressed genes for the 20 domains identified by FAST-G. (e): Scatter plot of two-dimensional PCs from the embeddings of FAST-G within the brain region. The depicted path corresponds to the inferred differentiation pathway determined by Slingshot for the six distinct subregions within the brain. Astro: Astrocytes, Olig: Oligodendrocytes. (f): Scatter plot of two-dimensional PCs from the embeddings of FAST-G for the expression levels of gene *Mt3* during embryo days ranging from E12.5 to E16.5.

After removing unwanted variations in expression profiles, we detected DEGs and recovered the temporal patterns for all 26 sections across five time points (see “Supplementary Notes”). In the DE analysis, we identified a total of 3663 DEGs with adjusted *p*-values *<* 0.001 across the 20 spatial domains identified by FAST-G (Supplementary Data 4). Among these genes, 194 were specific to Domain 1, corresponding to a brain-related subregion. The heatmap illustrates the findings and demonstrates the effective separation of DEGs across various spatial domains (Fig. 5d). By further investigation of mouse embryo spatial gene expression database, EMAGE [59], we detected marker genes for different cell types in mouse embryo, including *Fabp7* [60], *Hes5* [61], *Nefl*, *Nefm* [62] (Domains 1–6, brain), *Ptgds, Slc6a13* [63] (Domain 7, meninges), *Myl2*, *Myl3*, *Myl4*, *Myl7* [64] (Domains 8–10, heart), *Alb*, *Afp* [65] (Domain 11, liver), *Sftpc* [66], *Tcf21* [67] (Domain 12, lung), *Myh3*, *Myh8*, *Myl1* [68] (Domains 13–15, muscle), *Col19a1*, *Col2a1*, *Igf1*, *Sfrp2* (Domains 16–17, trunk somite), *Krt10*, *Krt15* [69] (Domain 18, epidermis/cavity), *Trps1* [70], *Wnt5a* [71] (Domain 19, jaw and tooth), *Krt1* [72] and *Perp* [73] (Domain 20, mucosal epithelium). Furthermore, we generated heatmaps to visualize the expression patterns of marker genes specific to the brain region based on spatial coordinates. These heatmaps confirmed the prominent expression levels within the corresponding brain region (Supplementary Fig. S22).

Next, we detected genes exhibiting temporal expression trends in the brain region (see “Supplementary Notes”; Supplementary Fig. S23). We observed decreased expression of *Hbb-y*, *Hba-x*, and *Hbb-bh1*, which plays a crucial role for cell development by enhancing oxygen transport and aligns with their expression patterns during the early stages of embryonic development [74]. We found an increasing trend in the expression of *Cd44* and *Cdk8*, which encodes a protein that belongs to the cyclin-dependent protein kinase (CDK) family and is known to play a crucial role in regulating the progression of the cell cycle [75].

By the DE genes of the six subclusters within the brain region, we were able to detect three cell types (astrocytes, oligodendrocytes1/2, neurons1/2/3). By inferring the trajectory using embeddings and cluster labels estimated by FAST, we identified the trajectory from glia cells (oligodendrocytes, astrocytes) to neurons (Fig. 5e), which is consistent with the findings in the existing literature [76]. By leveraging temporal information, we successfully detected genes that exhibit specific expression trends within each local region of the brain (Fig. 5f and Supplementary Fig. S24-S28). Notably, some of these genes serve as brain markers, such as *Mt3*, *Dbi* and *Ina*, while others are associated with the development of the nervous system, including *Zic1* and *Hes6*. Among these, we observed that *Mt3* initiates expression during embryo development in brain subcluster 1 at E13.5, suggesting a significant role for *Mt3* in the differentiation process of subcluster 1 (astrocytes; Fig. 5f). These findings provide valuable information on the intricate mechanisms that govern brain development.

## Discussion

We present FAST, a probabilistic generalized factor analysis for spatially aware dimension reduction of spatial transcriptomics across multiple sections. Uniquely, FAST takes into consideration the count nature of many existing SRT technologies and models the local spatial correlation structure for “factors” from a common loading matrix during dimension reduction across multiple sections, promoting its computational efficiency and enabling its scalability up to millions of spatial locations. As a result, the “factors” estimated in FAST can be taken as low-dimensional embeddings intrinsic to biological effects with preservation of spatial correlation structure and, thus, can be paired with many existing software/tools developed in scRNA-seq studies to facilitate downstream analyses in SRT studies. Notably, only FAST was found to be applicable to a mouse embryo Stereo-seq dataset across 26 sections varying from E12.5 to E16.5 with *>*2.3 million spatial locations. Moreover, the analysis was completed in approximately 2 hours, which cannot be achieved by other spatial dimension reduction methods. After obtaining low-dimensional embeddings, we applied iSC-MEB for integrative clustering analysis to align embeddings among multiple sections, followed by a module designed to remove unwanted variations prior to downstream analyses. Using the DLPFC Visium dataset with manual annotation as our benchmark, we have illustrated the benefits of using FAST for data visualization, DE analysis, CCI, and trajectory analyses across multiple sections.

FAST provides a useful tool that can interconnect with aberrant protein activity and somatic mutation analyses to delineate the spatial genomic landscape of cancer. In our analysis of the breast cancer Xenium dataset, FAST not only identified biologically separable clusters for many cell types, such as subtypes for breast cancer cells, fibroblasts, and T cells, but also, through further analysis of aberrant protein activity, showed that these identified clusters presented differential protein activities for immune-related transcription factors. Furthermore, FAST identified a carcinogenesis factor, *CCNH*, which plays an important role in carcinogenesis [77] and was shown to be associated with many signaling pathways in breast cancer [78]. When applied to a HCC Visium dataset, FAST provided estimated embeddings similar to those from PRECAST; however, the computational process was completed in only 142 seconds. Further mutation analysis showed that somatic mutations were more prevalent in TNE than in stroma and identified four genes containing top SNPs with somatic mutations. Among these genes, *Cers2* plays a vital role in preserving hepatic chromosome polyploidization during cell division by regulating the expression of Mad2 in mouse [79], and is closely associated with the progression of liver cancer [80]. Takeda et al. [81] reported that *ETS2* functions as a key downstream transcription factor and assembles a transcription complex with *MLL* in HCC cells that directly activates *MMP1* and *MMP3*.

FAST has some caveats that may require further exploration. First, it would be interesting to perform joint dimension reduction for both single-cell multimodal omic data and SRT data with single-cell resolution, thus promoting inference for cis-regulatory interactions and/or defining gene-regulatory networks for transcription factors. Second, FAST performs spatial dimension reduction in an unsupervised manner. When manual annotations are available for some sections, it may be preferable to adopt a semi-supervised method by utilizing partial information about annotation labels across all sections. Finally, in the era of deep learning, performing a deep spatial encoding for SRT datasets would help extract a low-dimensional representation capturing nonlinear biological effects. In this study, we demonstrate the potential of FAST as a prototype for scalable extraction of cross-section embeddings among a large variety of spatial locations. Future exploration of all these issues is warranted to further confirm the important value of FAST as a method for spatially aware dimension reduction.

## Methods

### FAST model

Here, we present a basic overview of FAST; further details are available in the Supplementary Notes. For the *M* SRT sections, we observed a count expression matrix for each section. FAST has the ability to model these count matrices or log-normalized matrices of gene expressions using either a log link or identity link for multiple sections (Fig. 1a). We primarily focus on introducing the Poisson version of FAST (FAST-P) for count matrices, while details about the Gaussian version (FAST-G) can be found in the Supplementary Notes.

Specifically, we observe an *n_m_ ×p* count expression matrix **X***_m_* = (**x***_m_*_1_*, · · ·,* **x***_mi_, · · ·,* **x***_mnm_*)^T^ for section *m* (= 1*, · · ·, M*), where **x***_mi_* = (*x_mi_*_1_*, · · ·, x_mip_*) is a *p*-dimensional expression vector for each location *s_mi_ ∈* R^2^ in the section *m* on square or hexagonal lattices, among others. FAST models the count expression level of gene *j*, *x_mij_*, with its latent low-dimensional features **v***_mi_* via a log link as

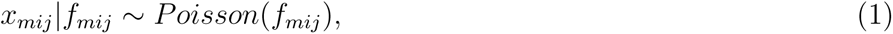

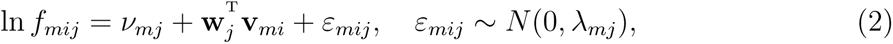

where *f_mij_* is an unknown Poisson rate that represents the underlying gene-expression level, *ν* = (*ν, · · ·, ν*) *∈* R*^p^* is a sample-specified intercept term, *W* = (**w** *, · · ·,* **w**)^T^ *∈* R*^p×q^* is a sample-shared loading matrix that captures the shared information of sections, **v***_mi_* is a *q*-dimensional factors that captures the biological information, and ***ɛ****_mi_* = (*ε_mi_*_1_*, · · ·, ε_mip_*) *∼ N* (**0**, Λ*_m_*) is an error term that captures the overdispersion, and Λ*_m_* = diag(*λ_m_*_1_*, · · ·, λ_mp_*) is the covariance matrix of the error term. To account for the spatial dependence among spatial locations within each section, we adopted a continuous multivariate Hidden Markov Random Field (HMRF) model for the factor **v***_mi_*, which captures spatial dependencies in the embedding space, referred to as a spatial factor. Specifically, we assumed an intrinsic CAR model [82] for **v***mi*

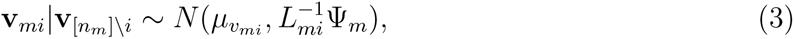

where subscript [*n_m_*] \ *i* denotes all locations but *s_mi_* in the section *m*, *L_mi_* is the number of neighbors of location *s_mi_* in section 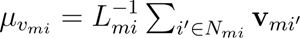 is the conditional mean relevant to the neighbors of the location *s_mi_*, and Ψ*_m_* is a *q × q* conditional covariance matrix for the elements of **v***_ri_*. The intrinsic CAR model (3) models the local dependence of the spatial factor **v***_mi_* for the location *s_mi_* in section *m* via the factors of its neighboring locations (Fig. 1a).

### Embedding alignment and spatial clustering

Once we obtained the uncorrected embeddings of FAST or other compared methods, we applied iSC-MEB [83] for joint embedding alignment and spatial clustering. However, before running iSC-MEB, we needed to determine the optimal number of clusters. Therefore, we first utilized Harmony to obtain batch corrected embeddings, followed by Louvain clustering to select the appropriate number of clusters. After determining the optimal number of clusters, we ran iSC-MEB; the implementation details can be found in the Supplementary Notes. After obtaining aligned embeddings and clusters, we compared the performance of FAST and other methods in visualization of the aligned embeddings, batch removal of the aligned embeddings, and spatial clustering.

### Unwanted variation removal for gene expressions

To remove unwanted variations in gene expression from the multiple sections, we applied iSC-MEB [83] to *V̂*, where *V̂* is the combined unaligned embeddings obtained by FAST. By doing so, we obtained the posterior probabilities of the cluster labels, *r̂_mi_*. To further address unwanted variations, we used a set of housekeeping genes, which are not influenced by other biological factors [84], as negative control genes; for more information on the selection of housekeeping genes, please refer to the Supplementary Notes.

Using the selected *L* housekeeping genes, we performed PCA and obtained the top 10 principal components, *ĥ_mi_*, which we used as covariates to adjust for unwanted variation. Let *x̃_mij_* be the log-normalized expression value for gene *j* of spot *i* in section *m*. Then, we applied a spatial linear model to the normalized expression level of gene *j*,

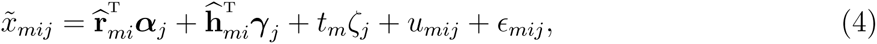

where ***α****_j_* is a *K*-dimensional vector for biological effects between cell/domain types, ***γ****_j_* is a 10-dimensional vector of regression coefficients associated with the unwanted factors, and *ζ_j_* is the regression coefficient denoting the temporal effect if the temporal information is available (e.g., in the embryo dataset), *u_mij_* follows a univariate intrinsic CAR model for retaining the spatial dependence of locations in the same section such that 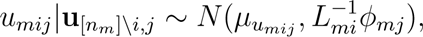, and 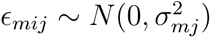 is an i.i.d (with respect to *i*) error term.

An ICM-EM algorithm was designed to solve the model parameters; details are provided in the Supplementary Notes. After obtaining the parameter estimates in Eqn. (4), users can remove batch effects from the original normalized gene expression using

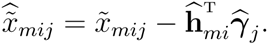

This strategy can also be applied to multiple sections from multiple biological conditions by adding additional covariates in the linear model when such information is available.

### Other analyses

After removing unwanted variation in the gene expression matrices of multiple sections, we performed DE analysis for the combined sections. To present biological discovery in different tissues, we applied various analyses including KEGG pathway analysis, CCI analysis, transcription factor analysis, cell-type deconvolution analysis, somatic mutation analysis, and analysis to detect genes with temporal trend of expressions. Details of these analyses are provided in the Supplementary Notes.

### Comparison of methods

Through simulations and with real-world data, we conducted a comprehensive comparison of FAST with existing methods for dimension reduction and spatial clustering.

To evaluate the biological embedding estimation performance of FAST, we used a range of spatially aware and non-spatially aware dimension reduction methods as benchmarks. Specifically, we compared FAST with (1) SpatialPCA [17] implemented in the R package *SpatialPCA* (version 1.2.0); (2) PRECAST [28] implemented in the R package *PRECAST* (version 1.6.1); (3) DR-SC [29] implemented in the R package *DR.SC* (version 3.2); (4) scVI [30] implemented in the Python module *scvi-tools* (version 0.20.3); (5) PCA; (6) multiBatchPCA [31] implemented in the R package *batchelor* (version 1.10.0); (7) NMF implemented in the R package *scater* (version 1.25.1); and (8) LIGER [32] implemented in the R package *rliger* (version 1.0.0). The first two methods are for spatial dimension reduction, whereas the remaining methods are for dimension reduction without consideration of spatial information. SpatialPCA has both full-rank and low-rank versions, denoted as SpatialPCA-F and SpatialPCA-L, respectively. We extracted 15-dimensional embeddings for all datasets and methods for comparison. See the Supplementary Notes for the details of each method compared.

To compare the clustering performance, after obtaining the low-dimensional embeddings for all methods except for LIGER, we applied iSC-MEB to conduct the joint embedding alignment and spatial clustering. Since LIGER had already addressed batch effects in its embeddings, we did not compare the spatial clustering performance using iSC-MEB.

### Evaluation metrics

#### McFadden’s adjusted R^2^

For DLPFC Visium and Embryo Stereo-seq datasets that have manual domain annotations serving as ground truth, we evaluated the performance of different methods in dimension reduction by measuring the association between the embeddings and ground truth for each section using a multinomial regression model with the ground truth as the response variable and the extracted embeddings as covariates for each slice. For the fitted model, we then calculated the adjusted 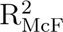 [85], which provides a measure of the amount of biological information contained in the extracted embeddings, and a higher value indicates better performance in dimension reduction.

#### ARI and NMI

For the DLPFC Visium and Embryo Steore-seq datasets, we evaluated the performance of different methods in spatial domain clustering by comparing the detected spatial domains with the ground truth. For this purpose, we employed standard clustering evaluation metrics, including ARI [86] and NMI [87].

#### Moran’s *I*

As a reflection of spatial autocorrelation, Moran’s *I* was used to measure the spatial information contained in the embeddings of each section obtained by FAST and other methods. It is defined as

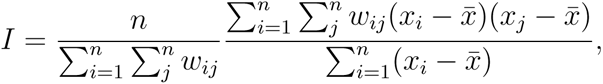

where *n* is the number of spatial units indexed by *i* and *j*, *x* is the variable of interest, *x̄* is the mean of *x*, and *w_ij_* is the (*i, j*)-th element of a matrix of spatial weights with zero on the diagonal. During the evaluation, we assigned a value of 1 to *w_ij_* if the spot *i* is a neighbor of the spot *j*, and otherwise *w_ij_* was set to 0.

### Simulations

We simulated three sections based on a single section of the human DLPFC Visium dataset (sample ID: 151672) with eight spatial domains [33], including layers 1-6, WM, and unknown cells (Unknown). Using these data, we initially established the spatial coordinates and spatial domains for the three sections. The first section was set using the same coordinates as the panel data. For the second section, the coordinates were selected such that the x coordinates were below the 90th percentile. For the last section, the coordinates were chosen such that the y coordinates were below the 90th percentile. The spatial domains for all three sections remained the same as the panel data; see Supplementary Fig. S1a.

Subsequently, we used the R package *splatter* (version 1.18.2) [88] to simulate gene expressions at predefined spatial coordinates of each section. This package generates gene expression data based on a gamma-Poisson distribution. We explored four different scenarios by adjusting the parameters of both batch effects and biological effects to either low or high values (batch effect=low or high; biological effect=low or high). The *splatter* package offers two parameters, namely de facScale and de prop, that enable the control of batch effects and biological effects. To generate the four scenarios, we set de facScale=0.2 and de facScale=0.6 for batch effect=low and high, respectively, and de prop=0.15 and de prop=0.3 for biological effect=low and high, respectively. The four scenarios tested were as follows: Scenario 1 (batch effect=low, biological effect=high), Scenario 2 (batch effect=low, biological effect=low), Scenario 3 (batch effect=high, biological effect=high), and Scenario 4 (batch effect=high, biological effect=low).

### Gene selection for multi-section analysis

During the quality control (QC) process, we applied filtering criteria to eliminate genes with zero expression in multiple locations and locations with zero expression for numerous genes (see “Data resources”). For our analyses, we used the FindVariableFeatures function of the *Seurat* (version 4.1.1) R package, with the default settings, to identify highly variable genes (HVGs) in all four datasets. Specifically, we selected the top 2000 HVGs for each section. We then prioritized genes based on the frequency of their selection as HVGs across all sections and selected the top 2000 genes based on this criterion. These 2000 genes were used as input for the comparison of FAST with other analytical methods.

### Data resources

#### Human dorsolateral prefrontal cortex Visium dataset

We obtained spatial transcriptomic data for human DLPFC from the 10*×* Visium platform; data were downloaded from https://doi.org/10.5281/zenodo.4730634. The dataset comprised 12 postmortem DLPFC tissue sections from three independent neurotypical adult donors, and the raw expression count matrix contained 33,538 genes for each section, with a total of 47,681 spatial locations. To ensure data quality, we performed QC on each section, filtering genes with non-zero expression levels for *≤* 20 locations and locations with non-zero expression levels for *≤* 20 genes. This filtering step resulted in a set of 14,535 genes on average across a total of 47,680 spatial locations. The spatial domains annotated in all 12 sections were layer 1 (*n*=5321), layer 2 (*n*=2858), layer 3 (*n*=17,587), layer 4 (*n*=3547), layer 5 (*n*=7300), layer 6 (*n*=6201), WM (*n*=4514), and undetermined locations (*n* = 352), according to the cytoarchitecture of the original study [33]. In our analysis, we considered these manual annotations as the ground truth for evaluating the dimension reduction and clustering performance of the different methods.

#### Human breast cancer Xenium dataset

We collected a dataset from two tissue sections of a patient with breast cancer measured by Xenium In Situ technology [35]; data were downloaded from https://www.dropbox.com/s/t0 5w7ccufh1v0h8/xenium_prerelease_jul12_hBreast_replicates.tar?dl=0. The dataset consisted of 313 genes of interest, with 35,868 and 36,783 spatial locations from the two tissue sections, respectively. These target genes were selected and curated primarily based on single cell atlas data for human breast tissue [35]. To ensure data quality, we performed a QC step in which we filtered locations with non-zero expression levels for *≤* 15 genes. As a result, 35,015 and 35,932 locations were retained in the two tissue sections, respectively. Aberrant protein activity was inferred by VIPER to assess protein activity from gene expression data and the regulatory network. The spatial deconvolution was then performed to examine the spatial distribution of the compositions of different cells using RCTD.

#### Human hepatocellular carcinoma Visium dataset

The HCC dataset used in this study was derived from two tissue sections, one from the tumor and the other from the tumor-adjacent regions of a patient with hepatocellular carcinoma [34]. The dataset consisted of 36,601 genes from more than 9813 spatial locations. During the QC process, genes with non-zero expression levels for *≤* 20 locations and locations with non-zero expression levels for *≤* 20 genes were removed, resulting in a set of 14,851 genes on average from a total of 9813 locations. To identify the TNE and stroma regions, manual annotations were provided by a pathologist using the Visium companion H&E images. Somatic mutation analysis was performed to investigate differences in somatic mutations between the TNE and stromal regions.

#### Mouse embryo Stereo-seq dataset

We collected data for 26 mouse embryo Stereo-seq sections from https://db.cngb.org/stom ics/mosta/. The dataset comprised 27,295 genes on average, with 2,323,044 spatial locations recorded across the 26 sections, from embryo days E12.5 to E16.5. During the QC process, we initially eliminated genes with non-zero expression levels for *≤* 20 locations, as well as locations with non-zero expression levels for *≤* 20 genes. As a result, we were left with 14,307 genes on average, totaling 2,318,423 locations.

## Data availability

The four datasets used in this study are publicly available. These include the 12 human DLPFC Visium datasets (https://doi.org/10.5281/zenodo.4730634), two human breast cancer Xenium datasets (https://www.dropbox.com/s/t05w7ccufh1v0h8/xenium_prerelease_jul12_hBreast_replicates.tar?dl=0), four human HCC Visium datasets (Raw FASTQ data are available at https://www.ncbi.nlm.nih.gov/sra?linkname=bioproject_sra_all&from_uid=858545, and H&E images are available at https://doi.org/10.6084/m9.figshare. 21280569.v1 and https://doi.org/10.6084/m9.figshare.21061990.v1), and 26 mouse embryo Stereo-seq datasets (https://db.cngb.org/stomics/mosta/).

## Code availability

The FAST methods were implemented in an open-source R package that is publicly available at https://github.com/feiyoung/FAST. The code to reproduce the analysis can be found at https://github.com/feiyoung/FAST_Analysis.

## Author contributions

J.L. initiated and designed the study, W.L. implemented the model and developed the software tool; W.L. and X.Z. performed the simulation studies and the benchmark evaluation; J.L. wrote the manuscript, and W.L., X.Z., X.C., Z.F., H.L., J.C., L.S., T.Y., J.Y. and J.L. edited and revised the manuscript.

