## supplementary materials for "FAST: a fast and scalable factor analysis for spatially aware dimension reduction of multi-section spatial transcriptomics data" (from another version of this preprint: version 1, 2023-07-12)

This PDF file includes:

- Supplementary Notes
- Supplementary Figures S1-S32

### Supplementary Notes

### 1 ProFAST method

#### 1.1 Poisson version of ProFAST and its algorithm

Consider the Poisson version of ProFAST, which can be described as follows:

$$x_{mij}|f_{mi} \sim Poisson(f_{mij}) \quad (1)$$

$$\ln f_{mij} = e_{mij} = \nu_{mj} + \mathbf{w}_j^{\mathrm{T}}\mathbf{v}_{mi} + \varepsilon_{mij}, \varepsilon_{mij} \sim N(0, \lambda_{mj}), \quad (2)$$

where $f_{mij}$ represents the underlying gene expression level as an unknown Poisson rate, $e_{mij}$ is the logarithm Poisson rate, $\nu_m = (\nu_{m1}, \cdots, \nu_{mp}) \in \mathbb{R}^p$ denotes a sample-specified intercept term, $W = (\mathbf{w}_1, \cdots, \mathbf{w}_p)^{\mathrm{T}} \in \mathbb{R}^{p\times q}$ is a sample-shared loading matrix that captures the shared information of sections, $\mathbf{v}_{mi}$ is a $q$-dimensional spatial factor that incorporates the biological information, and $\boldsymbol{\epsilon}_{mi} = (\varepsilon_{mi1}, \cdots, \varepsilon_{mip})^{\mathrm{T}} \sim N(\mathbf{0}, \Lambda_m)$ is the error term capturing overdispersion, and $\Lambda_m = \mathrm{diag}(\lambda_{m1}, \cdots, \lambda_{mp})$ is the covariance matrix of the error term. We assume an intrinsic CAR model [1] for $\mathbf{v}_{mi}$

$$\mathbf{v}_{mi}|\mathbf{v}_{[n_m]\backslash i} \sim N(\mu_{v_{mi}}, L_{mi}^{-1}\Psi_m), \quad (3)$$

where subscript ${}_{[n_m]\backslash i}$ denotes all spots except $s_{mi}$ in the section $m$, $L_{mi}$ is the number of neighbors of spot $i$ in section $m$, $\mu_{v_{mi}} = L_{mi}^{-1}\sum_{i'\in N_{mi}} \mathbf{v}_{mi'}$ is the conditional mean relevant to the neighbors of the spot $s_{mi}$, $N_{mi}$ is the index set of the neighbors of the spot $s_{mi}$, and $\Psi_m$ is a $q \times q$ conditional covariance matrix for the elements of $\mathbf{v}_{mi}$.

##### 1.1.1 Pseudo log-likelihood and evidence lower bound

Let $X_m$ and $E_m$ be the $n_m$-by-$p$ matrices with $(i, j)$-th entry $x_{mij}$ and $e_{mij}$, respectively, $V_m = (\mathbf{v}_{m1}, \cdots, \mathbf{v}_{mn_m})^{\mathrm{T}} \in \mathbb{R}^{n_m\times q}$ and $\mathbf{v}_{N_{mi}} = (\mathbf{v}_{mi'}, i' \in N_{mi})^{\mathrm{T}} \in \mathbb{R}^{L_{mi}\times q}$. Recall $\mathbf{x}_{mi} = (x_{mi1}, \cdots, x_{mip})^{\mathrm{T}}$, and let $\mathbf{e}_{mi} = (e_{mi1}, \cdots, e_{mip})^{\mathrm{T}}$, $X = (X_1^{\mathrm{T}}, \cdots, X_M^{\mathrm{T}})^{\mathrm{T}} \in \mathbb{R}^{N\times p}$ with $N = \sum_{m=1}^M n_m$, $E = (E_1^{\mathrm{T}}, \cdots, E_M^{\mathrm{T}})^{\mathrm{T}} \in \mathbb{R}^{N\times p}$, and $V = (V_1^{\mathrm{T}}, \cdots, V_M^{\mathrm{T}}) \in \mathbb{R}^{N\times q}$. By assuming the number of neighbors for each spot to be equal, we can re-parameterize $L_{mi}^{-1}\Psi_m$ as $\tilde{\Psi}_m$. The model parameters involved are denoted as $\boldsymbol{\theta} = (W, \nu_m, \tilde{\Psi}_m, \Lambda_m, m \le M)$. Assuming that there is a prediction of $V$, denoted as $\widehat{V}$, the complete pseudo-likelihood [2] is given by

$$\tilde{P}(X, E, V) = \Pi_{m,i}P(\mathbf{x}_{mi}|\mathbf{e}_{mi})P(\mathbf{e}_{mi}|\mathbf{v}_{mi})P(\mathbf{v}_{mi}|\hat{\mathbf{v}}_{N_{mi}}).$$

Moreover, the full psuedo log-likelihood is

$$\begin{aligned} \ln \tilde{P}(X, E, V) &= \sum_{m,i} \ln P(\mathbf{x}_{mi}|\mathbf{e}_{mi}) + \ln P(\mathbf{e}_{mi}|\mathbf{v}_{mi}) + \ln P(\mathbf{v}_{mi}|\hat{\mathbf{v}}_{N_{mi}}) \\ &= \sum_{m,i,j} \left\{ x_{mij}e_{mij} - \exp(e_{mij}) - \frac{1}{2}\{(e_{mij} - \nu_{mj} - \mathbf{w}_j^{\mathrm{T}}\mathbf{v}_{mi})^2/\lambda_{mj} + \ln \lambda_{mj}\} \right\} \\ &- \sum_{m,i} \frac{1}{2}\{\ln|\tilde{\Psi}_m| + (\mathbf{v}_{mi} - \widehat{\mu}_{v_{mi}})^{\mathrm{T}}\tilde{\Psi}_m^{-1}(\mathbf{v}_{mi} - \widehat{\mu}_{v_{mi}})\}, \end{aligned}$$

where $\widehat{\mu}_{v_{mi}} = \frac{1}{L_{mi}} \sum_{i' \in N_{mi}} \hat{\mathbf{v}}_{mi'}$.

To approximate the posterior of $(E, V)$, we use a mean field family given by

$$q(E, V) = \Pi_{m,i,j} N(e_{mij}; \varsigma_{mij}, \sigma^2_{mij}) N(\mathbf{v}_{mi}; \boldsymbol{\varsigma}_{v,mi}, D_{v,mi}), \tag{4}$$

where $N(e_{mij}; \varsigma_{mij}, \sigma^2_{mij})$ is a univariate Gaussian density function with mean $\varsigma_{mij}$ and variance $\sigma^2_{mij}$ for the variable $e_{mij}$ and $N(\mathbf{v}_{mi}; \boldsymbol{\varsigma}_{v,mi}, D_{v,mi})$ is a $q$-dimensional Gaussian density function with mean $\boldsymbol{\varsigma}_{v,mi}$ and covariance matrix $D_{v,mi}$ for the variable $\mathbf{v}_{mi}$. Based on the variational family (4), we can predict $V_m$ by the posterior mean:

$$\widehat{V}_m = (\boldsymbol{\varsigma}_{v,m1}, \cdots, \boldsymbol{\varsigma}_{v,mn_m})^{\mathrm{T}}. \tag{5}$$

and the conditional mean is given by

$$\widehat{\mu}_{v_{mi}} = \frac{1}{L_{mi}} \sum_{i' \in N_{mi}} \hat{\mathbf{v}}_{mi'}. \tag{6}$$

Let $\boldsymbol{\psi} = (\varsigma_{mij}, \sigma^2_{mij}, \boldsymbol{\varsigma}_{v,mi}, D_{v,mi}, m = 1, \cdots, M, i = 1, \cdots, n_m, j = 1, \cdots, p)$ be the collection of the variational parameters. The evidence lower bound (ELBO) is given by

$$\begin{aligned}
& ELBO(\boldsymbol{\theta}, \boldsymbol{\psi}) \\
= \;& \mathrm{E}_q \ln \tilde{P}(X, E, V) - \mathrm{E}_q \ln q(E, V) \\
= \;& \sum_{m,i,j} \left\{ x_{mij}\varsigma_{mij} - \exp(\varsigma_{mij} + \sigma^2_{mij}/2) \right\} \\
- \;& \frac{1}{2} \sum_{m,i,j} \left\{ \{(\varsigma_{mij} - \nu_{mj} - \mathbf{w}_j^{\mathrm{T}} \varsigma_{v,mi})^2/\lambda_{mj} + (\sigma^2_{mij} + \mathbf{w}_j^{\mathrm{T}} D_{v,mi}\mathbf{w}_j)/\lambda_{mj} + \ln \lambda_{mj} \} \right\} \\
- \;& \frac{1}{2} \sum_{m,i} \{ \ln |\tilde{\Psi}_m| + (\boldsymbol{\varsigma}_{v,mi} - \widehat{\mu}_{v_{mi}})^{\mathrm{T}} \tilde{\Psi}_m^{-1} (\boldsymbol{\varsigma}_{v,mi} - \widehat{\mu}_{v_{mi}}) + tr(\tilde{\Psi}_m^{-1} D_{v,mi}) \} \\
+ \;& \frac{1}{2} \sum_{m,i} (\sum_j \ln \sigma^2_{mij} + \ln |D_{v,mi}|) + c.
\end{aligned} \tag{7}$$

where $c$ is a general constant that is independent of $(\boldsymbol{\theta}, \boldsymbol{\psi})$.

##### 1.1.2 Variational ICM-EM algorithm

Initially, our focus lies on updating the variational parameters of $E$. However, deriving variational inference becomes challenging due to the lack of conjugacy between $q(E)$ and $P(X|E)$. To overcome this obstacle, we resort to the Laplace approximation [3] in order to obtain an approximate posterior distribution of $E$. Since $P(E|X) \propto P(X|E)P(E|Z, V)$, we employ a Taylor approximation around the Maximum A Posterior (MAP) point of $P(X|E) \cdot P(E|Z, V)$ to construct a Gaussian approximation as a substitute for the posterior distribution. Note $\ln P(X|E)P(E|Z, V) = \sum_{m,i,j} \left\{ x_{mij}e_{mij} - \exp(e_{mij}) - \frac{1}{2}\{(e_{mij} - \nu_{mj} - \mathbf{w}_j^{\mathrm{T}} \mathbf{v}_{mi})^2/\lambda_{mj}\} \right\} + c$. Using the fact that $\ln q(e_{mij}) = g_{mij}(e_{mij}) + c$, where $g_{mij}(e) = x_{mij}e - \exp(e) - \frac{1}{2}\mathrm{E}_{q(\mathbf{v}_{mi})}\{(e - \nu_{mj} - \mathbf{w}_j^{\mathrm{T}} \mathbf{v}_{mi})^2/\lambda_{mj}\}$, the posterior mean and variance of $e_{mij}$ can be estimated by

$$\hat{\varsigma}_{mij} = \arg\max_e g_{mij}(e), \qquad \hat{\sigma}^2_{mij} = -g''_{mij}(\hat{\varsigma}_{mij})^{-1}, \tag{8}$$

where the specific forms are $g_{mij}(e) = x_{mij}e - \exp(e) - \frac{1}{2}\lambda_{mj}^{-1}\{(e - \nu_{mj} - \mathbf{w}_j^{\mathrm{T}}\varsigma_{v,mi})^2 + \mathbf{w}_j^{\mathrm{T}} D_{v,mi}\mathbf{w}_j\}$ and $g''_{mij}(e) = -\exp(e) - \lambda_{mj}^{-1}$.

To update the variational parameters of $\mathbf{v}_{mi}$, we differentiate the ELBO function $ELBO(\boldsymbol{\theta}, \boldsymbol{\psi})$ with respect to $D_{v,mi}$ and $\boldsymbol{\varsigma}_{v,mi}$, and set the resulting derivatives to zero. After performing some algebraic operations, we obtain the following expressions:

$$D_{v,mi} = (W^{\mathrm{T}}\Lambda_m^{-1}W + \tilde{\Psi}_m^{-1})^{-1}, \tag{9}$$

$$\boldsymbol{\varsigma}_{v,mi} = D_{v,mi}\left\{\sum_j (\varsigma_{mij} - \nu_{mj})\lambda_{mj}^{-1}\mathbf{w}_j + \tilde{\Psi}_m^{-1}\widehat{\mu}_{v_{mi}}\right\}. \tag{10}$$

By taking derivative of $ELBO(\boldsymbol{\theta}, \boldsymbol{\psi})$ with respect to the model parameters, we obtain

$$\mathbf{w}_j = \left(\sum_{m,i}(\boldsymbol{\varsigma}_{v,mi}\boldsymbol{\varsigma}_{v,mi}^{\mathrm{T}} + D_{v,mi})/\lambda_{mj}\right)^{-1}\left(\sum_{m,i}(\varsigma_{mij} - \nu_{mj})\varsigma_{v,mi}^{\mathrm{T}}/\lambda_{mj}\right), \tag{11}$$

$$\nu_{mj} = \frac{1}{n_m}\sum_i(\varsigma_{mij} - \mathbf{w}_j^{\mathrm{T}}\boldsymbol{\varsigma}_{v,mi}), \tag{12}$$

$$\tilde{\Psi}_m = \frac{1}{n_m}\sum_i\left\{(\boldsymbol{\varsigma}_{v,mi} - \widehat{\mu}_{v_{mi}})(\boldsymbol{\varsigma}_{v,mi} - \widehat{\mu}_{v_{mi}})^{\mathrm{T}} + D_{v,mi}\right\}, \tag{13}$$

$$\lambda_{mj} = \frac{1}{n_m}\sum_i\left\{(\varsigma_{mij} - \nu_{mj} - \mathbf{w}_j^{\mathrm{T}}\boldsymbol{\varsigma}_{v,mi})^2 + (\sigma_{mij}^2 + \mathbf{w}_j^{\mathrm{T}}D_{v,mi}\mathbf{w}_j)\right\}. \tag{14}$$

The designed variational ICM-EM algorithm is implemented by iteratively predicting $V_m$ and updating the variational parameters and model parameters until convergence is reached, utilizing equations (5), (8) to (14). The algorithm is summarized in Algorithm 1.

---

**Algorithm 1** The variational ICM-EM algorithm for Poisson version of ProFAST

---

**Input:** Count matrices $\{X_m, m = 1, \cdots, M\}$, spatial coordinates $\{S_m, m = 1, \cdots, M\}$, the number of factors $q$, maximum iterations $maxIter$, relative tolerance of ELBO $epsELBO$

**Output:** $\widehat{V}_m, \widehat{W}, \widehat{\nu}_m, \widehat{\tilde{\Psi}}_m, \widehat{\Lambda}_m$

Initialize $\boldsymbol{\psi}^{(0)} = (\varsigma_{mij}^{(0)}, \sigma_{mij}^{2,(0)}, \boldsymbol{\varsigma}_{v,mi}^{(0)}, D_{v,mi}^{(0)}, m = 1, \cdots, M, i = 1, \cdots, n_m, j = 1, \cdots, p)$, $\boldsymbol{\theta}^{(0)} = (W^{(0)}, \nu_m^{(0)}, \tilde{\Psi}_m^{(0)}, \Lambda_m^{(0)}, m = 1, \cdots, M)$

**for** each $t \in 1, \cdots, maxIter$ **do**

  Predict $V_m$ using Equation (5), i.e., $\widehat{V}_m^{(t)} = (\boldsymbol{\varsigma}_{v,m1}^{(t-1)}, \cdots, \boldsymbol{\varsigma}_{v,mn_m}^{(t-1)})^{\mathrm{T}}, m = 1, \cdots, M$;

  Update variational parameters $\boldsymbol{\psi}^{(t)}$ based on Equations (8)–(10);

  Update model parameters $\boldsymbol{\theta}^{(t)}$ based on Equations (11)–(14);

  Evaluate the evidence lower bound, $ELBO_t = ELBO(\boldsymbol{\theta}^{(t)}, \boldsymbol{\psi}^{(t)})$,by (7).

  **if** $|ELBO_t - ELBO_{t-1}|/|ELBO_{t-1}| < epsELBO$ **then**

    break;

  **end if**

**end for**

**return** $\widehat{V}_m, \widehat{W}, \widehat{\nu}_m, \widehat{\tilde{\Psi}}_m, \widehat{\Lambda}_m, m = 1, \cdots, M$

---

#### 1.2 Gaussian version of ProFAST model and its algorithm

In this subsection, we discuss the Gaussian version of ProFAST model and its corresponding ICM-EM algorithm. Let $\tilde{\mathbf{x}}_{mi} = (\tilde{x}_{mi1}, \cdots, \tilde{x}_{mip})^{\mathrm{T}}$ be the $p$-dimensional log-normalized expression vector for each spot $s_{mi} \in \mathbb{R}^2$ in the section $m$. The Gaussian version of the ProFAST model is formulated by

$$\tilde{\mathbf{x}}_{mi} = \nu_m + W\mathbf{v}_{mi} + \boldsymbol{\epsilon}_{mi}, \tag{15}$$

where $\nu_m \in \mathbb{R}^p$ is a sample-specified intercept term, $W \in \mathbb{R}^{p\times q}$ is a sample-shared loading matrix that captures the shared biological information of sections, $\mathbf{v}_{mi}$ is a $q$-dimensional spatial factor that captures the biological information, and $\boldsymbol{\epsilon}_{mi}$ is the residual error and follows $N(\mathbf{0}, \Lambda_m)$ with $\Lambda_m = \mathrm{diag}(\lambda_{m1}, \cdots, \lambda_{mp})$. $\mathbf{v}_{mi}$ is assumed to follow the same intrinsic CAR model (3) to incorporate the spatial dependence among spots in each section.

##### 1.2.1 Pseudo log-likelihood

Let $\widetilde{X}_m$ be a $n_m$-by-$p$ matrix with $(i,j)$-th entry $\tilde{x}_{mij}$, and $V_m = (\mathbf{v}_{m1}, \cdots, \mathbf{v}_{mn_m})^{\mathrm{T}} \in \mathbb{R}^{n_m\times q}$. Let $\widetilde{X} = (\widetilde{X}_1^{\mathrm{T}}, \cdots, \widetilde{X}_M^{\mathrm{T}})^{\mathrm{T}} \in \mathbb{R}^{N\times p}$ with $N = \sum_{m=1}^M n_m$, and $V = (V_1^{\mathrm{T}}, \cdots, V_M^{\mathrm{T}}) \in \mathbb{R}^{N\times q}$. The involved parameters are denoted as $\boldsymbol{\theta} = (W, \nu_m, \tilde{\Psi}_m, \Lambda_m, m \le M)$. Assuming that there is a prediction of $V$, denoted as $\widehat{V}$, the complete pseudo-likelihood [2] is defined as

$$\tilde{P}(\widetilde{X}, V) = \Pi_{m,i} P(\tilde{\mathbf{x}}_{mi}|\mathbf{v}_{mi}) P(\mathbf{v}_{mi}|\hat{\mathbf{v}}_{N_{mi}}).$$

Furthermore, the pseudo full log-likelihood is given by

$$\ln \tilde{P}(\widetilde{X}, V) = \sum_{m,i} \ln P(\tilde{\mathbf{x}}_{mi}|\mathbf{v}_{mi}) + \ln P(\mathbf{v}_{mi}|\hat{\mathbf{v}}_{N_{mi}}),$$

and the observed pseudo log-likelihood is

$$\tilde{P}(\widetilde{X}) = \sum_{m,i} \ln P(\tilde{\mathbf{x}}_{mi}; \boldsymbol{\theta}),$$

where $\tilde{\mathbf{x}}_{mi} \sim N(\nu_m + W\widehat{\mu}_{v_{mi}}, W\tilde{\Psi}_m W^{\mathrm{T}} + \Lambda_m)$ with $\widehat{\mu}_{v_{mi}} = L_{mi}^{-1}\sum_{i'\in N_{mi}} \widehat{\mathbf{v}}_{mi'}$.

##### 1.2.2 ICM-EM algorithm

Let $D_m = (W\tilde{\Psi}_m W^{\mathrm{T}} + \Lambda_m)^{-1}$. Applying the Woodbury matrix identity, we derive the expression: $D_m = \Lambda_m^{-1} - \Lambda_m^{-1} W(\tilde{\Psi}_m^{-1} + W^{\mathrm{T}}\Lambda_m^{-1}W)^{-1}W^{\mathrm{T}}\Lambda_m^{-1}$. With all the model parameters determined, we can obtain predictions $\widehat{V}$ by maximizing the pseudo posterior distribution of $V$, resulting in estimated values $\hat{\mathbf{v}}_{mi} = \mathrm{E}(\mathbf{v}_{mi}|\mathbf{x}_{mi})$.

Considering that $(\tilde{\mathbf{x}}_{mi}, \mathbf{v}_{mi})$ follows a multivariate normal distribution, we can utilize the properties of conditional expectation and covariance for the joint normal distribution. This leads to the following expressions:

$$\begin{aligned}
\mathrm{E}(\mathbf{v}_{mi}|\tilde{\mathbf{x}}_{mi}; \boldsymbol{\theta}) &= \widehat{\mu}_{v_{mi}} + \tilde{\Psi}_m W^{\mathrm{T}} D_m\{\tilde{\mathbf{x}}_{mi} - \nu_m - W\widehat{\mu}_{v_{mi}})\} \\
\mathrm{var}(\mathbf{v}_{mi}|\tilde{\mathbf{x}}_{mi}; \boldsymbol{\theta}) &= \tilde{\Psi}_m - \tilde{\Psi}_m W^{\mathrm{T}} D_m W\tilde{\Psi}_m.
\end{aligned}$$

After obtaining the value $\boldsymbol{\theta}$ in the $t$-th iteration, denoted as $\boldsymbol{\theta}^{(t)}$, we define $\mathrm{E}_t(\mathbf{v}_{mi}) = \mathrm{E}(\mathbf{v}_{mi}|\tilde{\mathbf{x}}_{mi};\boldsymbol{\theta}^{(t)})$, $\mathrm{var}_t(\mathbf{v}_{mi}) = \mathrm{var}(\mathbf{v}_{mi}|\tilde{\mathbf{x}}_{mi};\boldsymbol{\theta}^{(t)})$. Thus, we predict $V_m$ by

$$\widehat{\mathbf{v}}_{mi}^{(t)} = \mathrm{E}_t(\mathbf{v}_{mi}) \tag{16}$$

and $\widehat{\mu}_{v_{mi}}^{(t)} = L_{mi}^{-1}\sum_{i'\in N_{mi}} \widehat{\mathbf{v}}_{mi'}^{(t)}$. Hence, the Q-function is derived by taking the posterior expectation of $V$ conditional on $\widetilde{X}$ with respect to the pseudo full log-likelihood given $\boldsymbol{\theta} = \boldsymbol{\theta}^{(t)}$:

$$\begin{aligned}
& Q(\boldsymbol{\theta};\boldsymbol{\theta}^{(t)}) \\
= & \sum_{m,i} \mathrm{E}_{\boldsymbol{\theta}^{(t)}}[\ln P(\tilde{\mathbf{x}}_{mi}|\mathbf{v}_{mi};\boldsymbol{\theta}) + \ln P(\mathbf{v}_{mi}|\hat{\mathbf{v}}_{N_{mi}};\boldsymbol{\theta})] \\
= & \sum_{m,i} \mathrm{E}_{\boldsymbol{\theta}^{(t)}}[-\frac{1}{2}\{\ln|\Lambda_m| + (\tilde{\mathbf{x}}_{mi} - (\nu_m + W\mathbf{v}_{mi}))^{\mathrm{T}}\Lambda_m^{-1}(\tilde{\mathbf{x}}_{mi} - (\nu_m + W\mathbf{v}_{mi}))\} \\
- & \frac{1}{2}(\ln|\tilde{\Psi}_m| + (\mathbf{v}_{mi} - \widehat{\mu}_{v_{mi}}^{(t)})^{\mathrm{T}}\tilde{\Psi}_m^{-1}(\mathbf{v}_{mi} - \widehat{\mu}_{v_{mi}}^{(t)}))] \\
= & \sum_{m,i}[-\frac{1}{2}\{\ln|\Lambda_m| + (\tilde{\mathbf{x}}_{mi} - (\nu_m + W\mathrm{E}_t(\mathbf{v}_{mi})))^{\mathrm{T}}\Lambda_m^{-1}(\tilde{\mathbf{x}}_{mi} - (\nu_m + W\mathrm{E}_t(\mathbf{v}_{mi}))) \\
& + tr(W^{\mathrm{T}}\Lambda_m^{-1}W\mathrm{var}_t(\mathbf{v}_{mi}))\} - \frac{1}{2}(\ln|\tilde{\Psi}_m| + (\mathrm{E}_t(\mathbf{v}_{mi}) - \widehat{\mu}_{v_{mi}}^{(t)})^{\mathrm{T}}\tilde{\Psi}_m^{-1}(\mathrm{E}_t(\mathbf{v}_{mi}) - \widehat{\mu}_{v_{mi}}^{(t)}) \\
& + tr(\tilde{\Psi}_m^{-1}\mathrm{var}_t(\mathbf{v}_{mi}))].
\end{aligned}$$

In addition,

$$Q_1(\boldsymbol{\theta}^{(t)}) = \sum_{m,i} \mathrm{E}_{\boldsymbol{\theta}^{(t)}} \ln P(\mathbf{v}_{mi}|\tilde{\mathbf{x}}_{mi}) = -\frac{1}{2}\sum_{m,i} \ln|\mathrm{var}_t(\mathbf{v}_{mi})|$$

and

$$ELBO(\boldsymbol{\theta}) = Q(\boldsymbol{\theta};\boldsymbol{\theta}) - Q_1(\boldsymbol{\theta}). \tag{17}$$

Taking derivatives of $Q(\boldsymbol{\theta};\boldsymbol{\theta}^{(t)})$ with respect to each parameter in $\boldsymbol{\theta}$ and setting the derivatives to zeros, we obtain the iterative solution of model parameters in the $(t+1)$-th iteration, given by

$$\begin{aligned}
\tilde{\Psi}_m &= \frac{1}{n_m}\sum_i \{(\mathrm{E}_t(\mathbf{v}_{mi}) - \widehat{\mu}_{v_{mi}}^{(t)})(\mathrm{E}_t(\mathbf{v}_{mi}) - \widehat{\mu}_{v_{mi}}^{(t)})^{\mathrm{T}} + \mathrm{var}_t(\mathbf{v}_{mi})\}, & (18) \\
\nu_m &= \frac{1}{n_m}\sum_i (\tilde{\mathbf{x}}_{mi} - W\mathrm{E}_t(\mathbf{v}_{mi})) & (19) \\
\mathrm{vec}(W^{\mathrm{T}}) &= B_w^{-1}\mathrm{vec}(A_w^{\mathrm{T}}), & (20) \\
& \text{where } A_w = \sum_m \sum_i \left\{\Lambda_m^{-1}(\tilde{\mathbf{x}}_{mi} - \nu_m)\mathrm{E}_t(\mathbf{v}_{mi})\right\}, \\
& \qquad B_w = \sum_m \sum_i \Lambda_m^{-1} \otimes \left(\mathrm{E}_t(\mathbf{v}_{mi})\mathrm{E}_t(\mathbf{v}_{mi})^{\mathrm{T}} + \mathrm{var}_t(\mathbf{v}_{mi})\right), \\
\Lambda_m &= \frac{1}{n_m}\sum_i \{(\tilde{\mathbf{x}}_{mi} - \nu_m - W\mathrm{E}_t(\mathbf{v}_{mi}))(\tilde{\mathbf{x}}_{mi} - \nu_m - W\mathrm{E}_t(\mathbf{v}_{mi}))^{\mathrm{T}} \\
&+ W\mathrm{var}_t(\mathbf{v}_{mi})W^{\mathrm{T}}\}, & (21)
\end{aligned}$$

where the operator vec vertically stacks the columns of a matrix into a column vector.

The designed ICM-EM algorithm is implemented by iteratively predicting $V_m$ and updating model parameters until convergence is reached, utilizing equations (16), (18) to (21). The algorithm is summarized in Algorithm 2.

**Algorithm 2** The ICM-EM algorithm for Gaussian version of ProFAST

**Input:** Log-normalized expression matrices $\{\widetilde{X}_m, m = 1, \cdots, M\}$, spatial coordinates $\{S_m, m = 1, \cdots, M\}$, the number of factors $q$, maximum iterations $maxIter$, relative tolerance of ELBO $epsELBO$

**Output:** $\widehat{V}_m, \widehat{W}, \widehat{\nu}_m, \widehat{\tilde{\Psi}}_m, \widehat{\Lambda}_m$

Initialize $\boldsymbol{\theta}^{(0)} = (W^{(0)}, \nu_m^{(0)}, \tilde{\Psi}_m^{(0)}, \Lambda_m^{(0)}, m = 1, \cdots, M)$
**for** each $t \in 1, \cdots, maxIter$ **do**
  Predict $V_m$ using Equation (16), denoted by $V_m^{(t)}, m = 1, \cdots, M$;
  Update model parameters $\boldsymbol{\theta}^{(t)}$ based on Equations (18)–(21);
  Evaluate the evidence lower bound, $ELBO_t = ELBO(\boldsymbol{\theta}^{(t)})$,by (17).
  **if** $|ELBO_t - ELBO_{t-1}|/|ELBO_{t-1}| < epsELBO$ **then**
    break;
  **end if**
**end for**
**return** $\widehat{V}_m, \widehat{W}, \widehat{\nu}_m, \widehat{\tilde{\Psi}}_m, \widehat{\Lambda}_m, m = 1, \cdots, M$

### 2 Details of compared methods

We provided detailed information about the dimension reduction methods we compared.

1. SpatialPCA [4] is a spatially aware dimension reduction method designed for inferring low-dimensional representations of gene expression data in a single section of spatial transcriptomics. To perform dimension reduction for multi-sections, we combined gene expression matrices and differentiated the spatial coordinates from different sections by adding a large value, $(m - 1) \times 10^6$, for section $m$. Using the R package *SpatialPCA* (version 1.2.0) and following the author's pipeline `https://lulushang.org/SpatialPCA_Tutorial/slideseq.html`, we input the combined count matrix and corresponding spatial coordinates into the function `CreateSpatialPCAObject`, with `min.locations = 0` and `min.features = 0`. Additionally, we used the `customGenelist` parameter to include our selected HVGs since we had already performed data filtering and selected the HVGs. Next, we employed the function `SpatialPCA_EstimateLoading` along with either `fast=TRUE` (SpatialPCA-L) or `fast=FALSE` (SpatialPCA-F), along with `SpatialPCnum=15`, to estimate the 15-dimensional loading matrix. Finally, we utilized the function `SpatialPCA_SpatialPCs` with either `fast=TRUE` (SpatialPCA-L) or `fast=FALSE` (SpatialPCA-F) to estimate the spatial factor matrix.

2. PRECAST [5] is a newly developed probabilistic embedding and clustering method with alignment for integrating spatial transcriptomics data. Using the R package *PRECAST* (version 1.6.1), we applied the R function `ICM.EM` with `q=15` to estimate the 15-dimensional

embeddings and used the function `SelectModel` to determine the optimal number of clusters.

3. DR-SC [6] is a joint dimension reduction and spatial clustering method specifically designed for single-section spatial transcriptomics. We performed dimension reduction for multi-sections using the same strategy as that in SpatialPCA. With the R package *DR.SC* (version 3.2), we applied the function `DR.SC_fit` with `q=15` to extract 15-dimensional factors and used the function `selectModel` to determine the optimal number of clusters.

4. scVI [7] is a deep learning-based dimension reduction method. We followed the scVI pipeline provided at `https://docs.scvi-tools.org/en/stable/tutorials/notebooks/api_overview.html`, setting the parameter `n_latent=15` in the function `scvi.model.SCVI` to extract 15-dimensional embeddings. This analysis was based on the Python module *scvi-tools* (version 0.20.3).

5. PCA was implemented using the function `DR.SC:::wpca` with `q=15` and `weighted=FALSE` in the R package *DR.SC* (version 3.2).

6. multiBatchPCA [8] was implemented using the function `multiBatchPCA` with `d=15` in the R package *batchelor* (version 1.10.0).

7. NMF is a non-negative matrix factorization method that factorizes a non-negative matrix into two lower-rank non-negative matrices. Using the R package *scater* (version 1.25.1), we applied the function `calculateNMF` with `ncomponents=15` to extract 15-dimensional embeddings.

8. LIGER [9] is a commonly used dimension reduction and integration method in single-cell RNA sequencing data analysis. Following the author's pipeline described at `http://htmlpreview.github.io/?https://github.com/welch-lab/liger/blob/master/vignettes/Integrating_multi_scRNA_data.html`, we used the R package *rliger* (version 1.0.0) and set the parameter `k=15` in the function `optimizeALS` to extract 15-dimensional embeddings.

Please note that SpatialPCA-F is not capable of handling large-scale datasets such as the human DLPFC Visium dataset and the human breast cancer Xenium dataset. Therefore, we excluded it from the comparison of these two datasets. Similarly, SpatialPCA, PRECAST, DR-SC, and scVI were not compared for the mouse embryo Stereo-seq dataset as these methods are not suitable for handling such large-scale data.

### 3 Downstream analysis

#### 3.1 Embedding alignment and spatial clustering

For the four datasets, we initially applied Harmony to uncorrected embeddings from ProFAST and the other compared methods. Subsequently, we utilized Louvain clustering to select the optimal number of clusters. When running Harmony, we utilized the `HarmonyMatrix` function in the R package *harmony* (version 0.1.0) with `do_pca = FALSE` and default parameters for the other settings. For Louvain clustering, we employed the `FindClusters` function in the R package *Seurat* (version 4.1.1) with a resolution of 0.2 for the human DLPFC Visium dataset and the mouse embryo Stereo-seq dataset, 0.3 for the human HCC Visium dataset, and 0.5 for the human breast cancer Xenium dataset. After determining the optimal number of clusters, we utilized the `fit.iscmeb` function in the R package *iSC.MEB* (version 1.0.1) to jointly perform embedding alignment and spatial clustering with default parameters.

#### 3.2 Removing unwanted variation in multi-section SRT data

As described in the "Methods" section of the main text, we utilized the spatial linear model to model the normalized expression level of gene $j$,

$$\tilde{x}_{mij} = \widehat{\mathbf{r}}_{mi}^{\mathrm{T}} \boldsymbol{\alpha}_j + \widehat{\mathbf{h}}_{mi}^{\mathrm{T}} \boldsymbol{\gamma}_j + t_m \zeta_j + u_{mij} + \epsilon_{mij}, \tag{22}$$

where $\boldsymbol{\alpha}_j$ is a $K$-dimensional vector of biological effects between cell/domain types, $\boldsymbol{\gamma}_j$ is a 10-dimensional vector of regression coefficients associated with the unwanted factors, $\zeta_j$ is the regression coefficient denoting the temporal effect if time-point information is available such as in the embryo dataset, $u_{mij}$ is an intrinsic CAR model capturing the spatial dependence of locations in the same section such that $u_{mij} | \mathbf{u}_{[n_m] \backslash i, j} \sim N(\mu_{u_{mij}}, L_{mi}^{-1} \phi_{mj})$, and $\epsilon_{mij} \sim N(0, \sigma_{mj}^2)$ is an i.i.d (with respect to $i$) error term. For each gene $j$, we designed an ICM-EM algorithm to solve the model parameters.

##### 3.2.1 ICM-EM algorithm

For the gene $j$, let $\mathbf{u}_{mj} = (u_{m1j}, \cdots, u_{mn_m j})^{\mathrm{T}}$, $\mathbf{u}_{mN_i j} = (u_{mi'j}, i' \in N_{mi})^{\mathrm{T}}$, $\tilde{\mathbf{x}}_{mj} = (\tilde{x}_{m1j}, \cdots, \tilde{x}_{mn_m j})^{\mathrm{T}}$, and $\boldsymbol{\vartheta}_j = (\boldsymbol{\alpha}_j, \boldsymbol{\gamma}_j, \zeta_j, \phi_{mj}, \sigma_{mj}^2)$ be the collection of model parameters. Given a prediction of $\mathbf{u}_{mj}$, $\widehat{\mathbf{u}}_{mj}$, the full pseudo log-likelihood is given by

$$\begin{aligned} L_j(\boldsymbol{\vartheta}_j; \tilde{\mathbf{x}}_{mj}) &= \sum_m \sum_i \{\ln P(x_{mij} | u_{mij}) + \ln P(u_{mij} | \widehat{\mathbf{u}}_{mN_i j})\} \\ &= \sum_m \sum_i -\frac{1}{2} \Big\{ (\ln \sigma_{mj}^2 + (x_{mij} - (\widehat{\mathbf{r}}_{mi}^{\mathrm{T}} \boldsymbol{\alpha}_j + \widehat{\mathbf{h}}_{mi}^{\mathrm{T}} \boldsymbol{\gamma}_j + t_m \zeta_j + u_{mij}))^2 / \sigma_{mj}^2) \\ &\quad + \ln \psi_{mj} + (u_{mij} - \hat{\mu}_{u,mij})^2 / \psi_{mj} \Big\}, \end{aligned}$$

where $\hat{\mu}_{u,mij} = \frac{1}{|N_{mi}|} \sum_{i' \in N_{mi}} \hat{u}_{mi'j}$.

We can infer from the joint normal distribution of $(\tilde{x}_{mij}, u_{mij})$ that

$$\begin{aligned} \mathrm{E}(u_{mij} | \tilde{x}_{mij}; \boldsymbol{\vartheta}_j) &= \mu_{u,mij} + (\psi_{mj} + \sigma_{mj}^2)^{-1} \psi_{mj} (x_{mij} - \mu_{x,mij}), \\ \mathrm{var}(u_{mij} | \tilde{x}_{mij}; \boldsymbol{\vartheta}_j) &= \psi_{mj} - \psi_{mj} (\psi_{mj} + \sigma_{mj}^2)^{-1} \psi_{mj}, \end{aligned}$$

where $\mu_{x,mij} = \widehat{\mathbf{r}}_{mi}^{\mathrm{T}} \boldsymbol{\alpha}_j + \widehat{\mathbf{h}}_{mi}^{\mathrm{T}} \boldsymbol{\gamma}_j + t_m \zeta_j + \hat{\mu}_{u,mij}$. After obtaining the value $\boldsymbol{\vartheta}_j$ in the $t$-th iteration, denoted as $\boldsymbol{\vartheta}_j^{(t)}$, we define $\mathrm{E}_t(u_{mij}) = \mathrm{E}(u_{mij} | \tilde{x}_{mij}; \boldsymbol{\vartheta}_j^{(t)})$, $\mathrm{var}_t(u_{mij}) = \mathrm{var}(u_{mij} | \tilde{x}_{mij}; \boldsymbol{\vartheta}_j^{(t)})$. Thus, we predict $u_{mij}$ by

$$\hat{u}_{mij}^{(t)} = \mathrm{E}_t(u_{mij}) \tag{23}$$

and $\hat{\mu}^{(t)}_{u,mij} = \frac{1}{|N_{mi}|}\sum_{i' \in N_{mi}} \hat{u}^{(t)}_{mi'j}$.

Taking the posterior expectation of $\mathbf{u}_{mj}$ conditional on $\tilde{\mathbf{x}}_{mj}$ with respect to $L_j(\boldsymbol{\vartheta}_j; \tilde{\mathbf{x}}_{mj})$, we obtain the $Q$ function,

$$\begin{aligned} & Q(\boldsymbol{\vartheta}_j; \boldsymbol{\vartheta}_j^{(t)}) \\ = & \sum_m \sum_i -\frac{1}{2}\Big\{ (\ln \sigma^2_{mj} + (\tilde{x}_{mij} - (\widehat{\mathbf{r}}^{\mathrm{T}}_{mi}\boldsymbol{\alpha}_j + \widehat{\mathbf{h}}^{\mathrm{T}}_{mi}\boldsymbol{\gamma}_j + t_m\zeta_j + \mathrm{E}_t(u_{mij})))^2/\sigma^2_{mj} + \mathrm{var}_t(u_{mij})/\sigma^2_{mj}) \\ & + \ln \psi_{mj} + (\mathrm{E}_t(u_{mij}) - \hat{\mu}^{(t)}_{u,mij})^2/\psi_{mj} + \mathrm{var}_t(u_{mij})/\psi_{mj} \Big\}. \end{aligned}$$

Then taking derivative with respect to model parameters and setting the derivatives to zeros, we obtain the iterative equation:

$$\boldsymbol{\alpha}_j = (\sum_{m,i} \widehat{\mathbf{r}}_{mi}\widehat{\mathbf{r}}^{\mathrm{T}}_{mi}/\sigma^2_{mj})^{-1}\left[\sum_{m,i}\widehat{\mathbf{r}}_{mi}/\sigma^2_{mj}\{(\tilde{x}_{mij} - (\widehat{\mathbf{h}}^{\mathrm{T}}_{mi}\boldsymbol{\gamma}_j + t_m\zeta_j + \mathrm{E}_t(u_{mij})))\}\right], \quad (24)$$

$$\boldsymbol{\gamma}_j = (\sum_{m,i} \widehat{\mathbf{h}}_{mi}\widehat{\mathbf{h}}^{\mathrm{T}}_{mi}/\sigma^2_{mj})^{-1}\left[\sum_{m,i}\widehat{\mathbf{h}}_{mi}/\sigma^2_{mj}\{(\tilde{x}_{mij} - (\widehat{\mathbf{r}}^{\mathrm{T}}_{mi}\boldsymbol{\alpha}_j + t_m\zeta_j + \mathrm{E}_t(u_{mij})))\}\right], \quad (25)$$

$$\zeta_j = (\sum_m (n_m t^2_m/\sigma^2_{mj}))^{-1}(\sum_m t_m/\sigma^2_{mj}\sum_i(\tilde{x}_{mij} - (\widehat{\mathbf{r}}^{\mathrm{T}}_{mi}\boldsymbol{\alpha}_j + \widehat{\mathbf{h}}^{\mathrm{T}}_{mi}\boldsymbol{\gamma}_j + \mathrm{E}_t(u_{mij}))), \quad (26)$$

$$\sigma^2_{mj} = \frac{1}{n_m}\sum_i[(\tilde{x}_{mij} - (\widehat{\mathbf{r}}^{\mathrm{T}}_{mi}\boldsymbol{\alpha}_j + \widehat{\mathbf{h}}^{\mathrm{T}}_{mi}\boldsymbol{\gamma}_j + t_m\zeta_j + \mathrm{E}_t(u_{mij})))^2 + \mathrm{var}_t(u_{mij})], \quad (27)$$

$$\psi_{mj} = \frac{1}{n_m}\sum_i[(\mathrm{E}_t(u_{mij}) - \hat{\mu}^{(t)}_{u,mij})^2 + \mathrm{var}_t(u_{mij})]. \quad (28)$$

The designed ICM-EM algorithm is implemented by iteratively predicting $\mathbf{u}_{mj}$ and updating model parameters until convergence, utilizing equations (23), (24) to (28).

After obtaining the parameter estimates in Eqn. (22), users can remove unwanted variations from the original normalized gene expression using

$$\widehat{\tilde{x}}_{mij} = \tilde{x}_{mij} - \widehat{\mathbf{h}}^{\mathrm{T}}_{mi}\widehat{\boldsymbol{\gamma}}_j.$$

##### 3.2.2 Selection of housekeeping genes

In the model (22), we utilized a set of housekeeping genes as negative control genes, which are not influenced by other biological factors [10]. The set of housekeeping genes were selected as follows. To obtain the expression vectors of the housekeeping genes, we matched the set of housekeeping genes with genes passing QC for the dataset of each section. For the matched housekeeping genes, we identified the spatial pattern of each gene using SPARK-X [11]. We select the top 200 genes with lowest spatial patterns from the matched list of housekeeping genes in the LIBD human DLPFC dataset, the human hepatocellular carcinoma Visium dataset and the mouse embryo Stereo-seq dataset. Specifically, we selected the top 200 genes with the largest $p$-values in SPARK-X for each section. We then prioritized genes based on the frequency of their selection across all sections and selected the top 200 genes based on this criterion. By the properties of negative control genes, the expression vector of these genes satisfies

$$\tilde{x}_{L,mil} = \mathbf{h}^{\mathrm{T}}_{mi}\boldsymbol{\gamma}_{L,l} + \epsilon_{L,mil}, l = 1, \cdots, L.$$

By performing the PCA, we estimated $\mathbf{h}_{mi}$, $\widehat{\mathbf{h}}_{mi}$, as the factors capturing the unwanted variations. Among the target genes measured in the human breast cancer Xenium dataset, only six housekeeping genes matched. Therefore, we used the log-normalized expressions of these six genes to capture the unwanted variations.

#### 3.3 Differential expression analysis

After removing the unwanted variation in the gene expression matrices of multiple sections, we conducted DE analyses on all four datasets. Specifically, we utilized the `FindAllMarkers` function within the R package *Seurat* (version 4.1.1), employing default settings to identify the DEGs for each domain detected by ProFAST. We defined domain-specific DEGs as those with adjusted $p$-values $< 0.001$ and a log-fold change $> 0.25$. In the human DLPFC Visium dataset, based on the marker genes, we discovered that Domains 2 and 3 belong to the subclass of Layers 2/3. Consequently, we merged these domains and assigned them the label Layer 2/3. Similarly, we combined Domains 7 and 8 and designated them as WM. We then constructed a dot plot of normalized expression levels of DEGs for different layers identified by ProFAST-P.

#### 3.4 KEGG pathway analysis

To investigate the pathways associated with diseases and drugs in the breast cancer Xenium dataset, we conducted KEGG pathway analysis on the marker genes identified in each cluster obtained by ProFAST-P. To perform this analysis, we utilized g:Profiler with the g:SCS multiple testing correction method and applied a significance threshold of 0.05.

#### 3.5 Cell-cell interaction analysis

In the LIBD human DLPFC dataset, we conducted cell-cell interaction analysis for various spatial locations across multiple sections utilizing CellChat [12]. We utilized the expression matrix with unwanted variations removed and layer labels estimated by ProFAST-P as inputs for the functions in the *CellChat* (version 1.1.3) R package. Specifically, we determined the communication probability using the R function `computeCommunProb`, and inferred the cellular communication network and signaling pathways using the R function `computeCommunProbPathway`.

#### 3.6 PAGA trajectory inference

To create a spatial map of the DLPFC Visium dataset, we utilized the PAGA algorithm implemented in the Python module *scanpy* (version 1.9.1). This approach enabled us to maintain the overall topology in the embeddings obtained through ProFAST-P for each DLPFC section. In detail, we utilized the embeddings and layers obtained from ProFAST-P as inputs for the functions `scanpy.pp.neighbors` and `scanpy.tl.paga` within the Python module *scanpy*. These functions were employed to generate a graphical representation of the data, taking into account the neighborhood relationships between spots. Subsequently, we utilized the scanpy.pl.paga function to visualize the structure graph, which captures significant cellular transitions and relationships within the data.

#### 3.7 Somatic mutation analysis

For the HCC Visium dataset, we performed somatic mutation analysis from the SRT data. First, we applied the *GATK* tool [13] to call genotypes using the combined HCC mapped bam files. Specifically, we used MergeBamAlignment and MarkDuplicates for data cleanup. We then used SplitNCigarReads to split reads with $N$ in the cigar into multiple supplementary alignments and hard clips mismatching overhangs. The base quality scores were recalibrated by BaseRecalibrator to deal with the problems of over or under-estimated scores caused by errors of sequencing machines. Finally, We used HaplotypeCaller to conduct variant calling to obtain the genotypes.

Next, we obtained the RS number of the called SNPs from the 1000 Genomes Project databases and added the 21,315 SNPs of LD regions of the called SNPs by using the LDproxy function in the R package *LDlinkR* (version 1.2.3). We matched these SNPs to the HCC-related database (https://www.cbioportal.org/) and obtained 106 overlapped SNPs. For each of the 106 SNPs, we determined the reference count, alternate count and total count of somatic mutations in every location of each HCC section. We then focused our analysis on the somatic mutations of the 106 SNPs and compared their frequency in the tumor and non-tumor regions. We calculated the sum of the alternate counts of the 106 SNPs for all sections across the identified nine clusters and divided the number of locations to obtain the mutation count per location.

#### 3.8 Transcription factor analysis

In the breast cancer Xenium dataset, we assessed aberrant protein activity by Virtual Inference of Protein-activity by Enriched Regulon analysis (VIPER) [14], which enabled us to accurately assess protein activity from gene expression data and regulatory networks. Specifically, we obtained the human breast carcinoma context-specific regulatory network from the R package aracne.networks [15], and inferred protein activity for each section by VIPER. We then obtained the activity of five regulatory proteins for visualization in violin plots.

#### 3.9 Deconvolution analysis

We utilized Robust Cell Type Decomposition (RCTD) [16], a supervised learning technique, for deconvolution analysis. RCTD enabled us to break down each spatial transcriptomics pixel into a blend of individual cell types while adjusting for platform effects. For the breast cancer Xenium dataset, we utilized a 10× chromium-based scRNA dataset [17] collected from patients with primary untreated breast cancers. The reference data included 100,064 cells belonging to 49 cell types: two B cell types (B cells memory: $n = 2,581$; B cells naive: $n = 625$), five CAFs types (CAFs MSC iCAF-like s1: $n = 2,309$, CAFs MSC iCAF-like s2: $n = 844$, CAFs myCAF like s4: $n = 602$, CAFs myCAF like s5: $n = 2,265$, CAFs transitioning s3:

$n = 553$), five cancer types (cancer basal SC: $n = 4,312$, cancer cycling: $n = 5,359$, cancer Her2 SC: $n = 3,708$, cancer LumA SC: $n = 7,742$, cancer LumB SC: $n = 3,368$), two cycling cell types (cycling PVL: $n = 50$, cycling myeloid: $n = 463$), four endothelial cell types (endothelial ACKR1: $n = 4,611$, endothelial CXCL12: $n = 1,644$, endothelial lymphatic LYVE1: $n = 203$, endothelial RGS5: $n = 1,147$), luminal progenitors ($n = 1,992$), mature luminal ($n = 1,265$), 10 myeloid types (myeloid_c0_DC_LAMP3: $n = 131$, myeloid_c1_LAM1_FABP5: $n = 2,122$, myeloid_c10_macrophage_1_EGR1: $n = 2,150$, myeloid_c11_cDC2_CD1C: $n = 337$, myeloid_c12_monocyte_1_IL1B: $n = 1,339$, myeloid_c2_LAM2_APOE: $n = 1,126$, myeloid_c3_cDC1_CLEC9A: $n = 156$, myeloid_c4_DCs_pDC_IRF7: $n = 331$, myeloid_c5_macrophage_3_SIGLEC1: $n = 26$, myeloid_c7_monocyte_3_FCGR3A:$n = 54$, myeloid_c8_monocyte_2_S100A9: $n = 935$, myeloid_c9_macrophage_2_CXCL10: $n = 505$), myoepithelial ($n = 1,098$), plasmablasts ($n = 3,524$), three PVL types (PVL differentiated s3: $n = 3,487$, PVL immature s1: $n = 1,169$, PVL immature s2: $n = 717$), 12 T cell types (T cells_c0_CD4$^+$_CCR7: $n = 4,952$, T cells_c1_CD4$^+$_IL7R: $n = 7,786$, T cells_c10_NKT_cells_FCGR3A: $n = 1,122$, T cells_c11_MKI67: $n = 1,528$,T cells_c2_CD4$^+$_T-regs_FOXP3: $n = 4,246$, T cells_c3_CD4$^+$_Tfh_CXCL13: $n = 2,247$, T cells_c4_CD8$^+$_ZFP36: $n = 5,063$, T cells_c5_CD8$^+$_GZMK: $n = 284$, T cells_c6_IFIT1: $n = 1,017$, T cells_c7_CD8$^+$_IFNG: $n = 3,168$, T cells_c8_CD8$^+$_LAG3: $n = 1,955$, T cells_c9_NK_cells_AREG: $n = 1,846$). Subsequently, we merged the proportions of subtypes of B cells, CAFs, cancer cells, cycling cells, endothelial cells, myeloid, PVL and T cells into a single group for visualization, after obtaining the deconvolution outcomes using the aforementioned reference panel.

##### 3.10 Detection of the genes with temporal trend of expressions

For mouse embryo Stereo-seq dataset, we employed linear regression to identify genes exhibiting temporal trends in expression within the same tissue, with the expression level serving as the response variable and the time point as the covariate for each gene. Subsequently, we performed gene ranking based on adjusted $R^2$ values, considering only genes with adjusted $p$-values $<$ 0.001.

[3] Chong Wang and David M Blei. Variational inference in nonconjugate models. *Journal of Machine Learning Research*, 2013.

[13] Geraldine A Van der Auwera and Brian D O'Connor. *Genomics in the cloud: using Docker, GATK, and WDL in Terra*. O'Reilly Media, 2020.

[14] Mariano J Alvarez, Yao Shen, Federico M Giorgi, Alexander Lachmann, B Belinda Ding, B Hilda Ye, and Andrea Califano. Functional characterization of somatic mutations in cancer using network-based inference of protein activity. *Nature Genetics*, 48(8):838–847, June 2016.

[15] Alexander Lachmann, Federico M Giorgi, Gonzalo Lopez, and Andrea Califano. Aracne-ap: gene network reverse engineering through adaptive partitioning inference of mutual information. *Bioinformatics*, 32(14):2233–2235, 2016.

[16] Dylan M Cable, Evan Murray, Luli S Zou, Aleksandrina Goeva, Evan Z Macosko, Fei Chen, and Rafael A Irizarry. Robust decomposition of cell type mixtures in spatial transcriptomics. *Nature Biotechnology*, 40(4):517–526, 2022.

[17] Sunny Z Wu, Ghamdan Al-Eryani, Daniel Lee Roden, Simon Junankar, Kate Harvey, Alma Andersson, Aatish Thennavan, Chenfei Wang, James R Torpy, Nenad Bartonicek, et al. A single-cell and spatially resolved atlas of human breast cancers. *Nature genetics*, 53(9):1334–1347, 2021.

### 4 Supplementary Figures

#### 4.1 Validation using simulated data

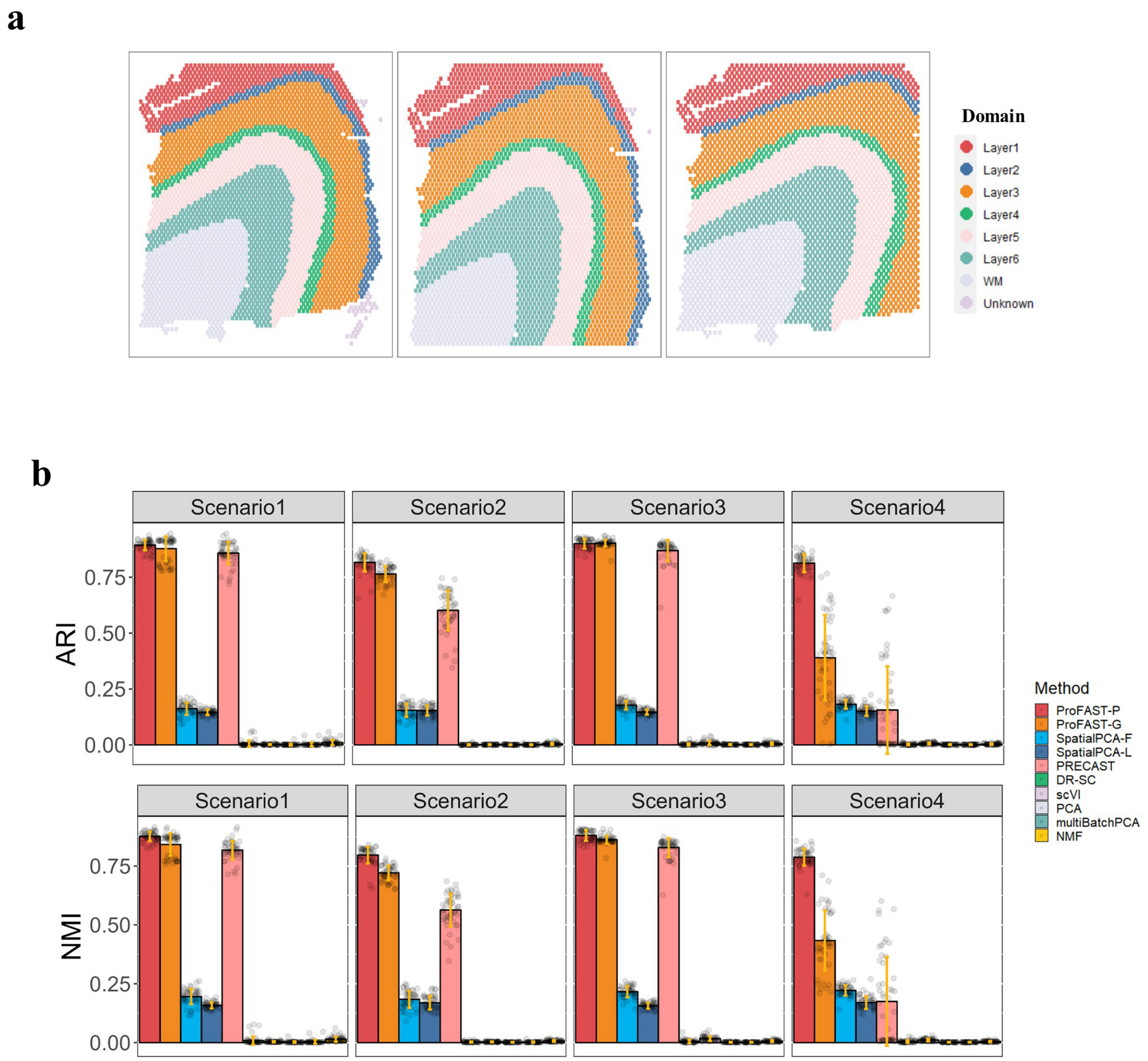


Figure S1: Comparison of clustering performance in simulations by generating data based on human dorsolateral prefrontal cortex (DLPFC) Visium data. (a): Visual representation of the spatial distribution of eight domains for each of the three simulated sections. (b): Comparison of clustering performance for ProFAST and other compared methods. iSC-MEB applied to perform spatial clustering based on the embeddings from ProFAST and other methods.

#### 4.2 Application to human dorsolateral prefrontal cortex Visium dataset

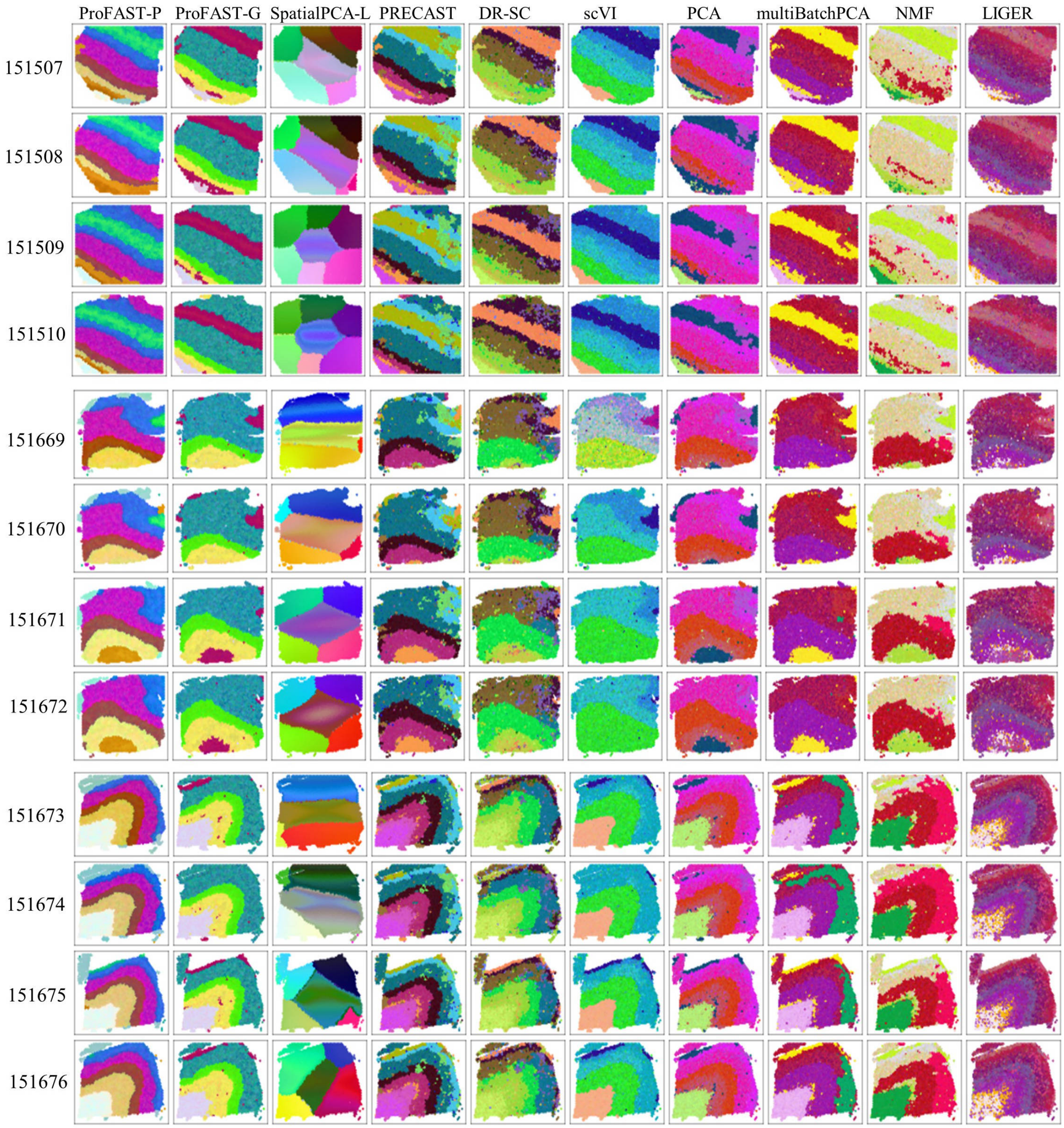


Figure S2: Dimension reduction with embedding alignment for the 12 dorsolateral prefrontal cortex Visium sections. UMAP RGB plots for the 12 sections based on the aligned embeddings via iSC-MEB using the low-dimensional embeddings from ProFAST and other methods.

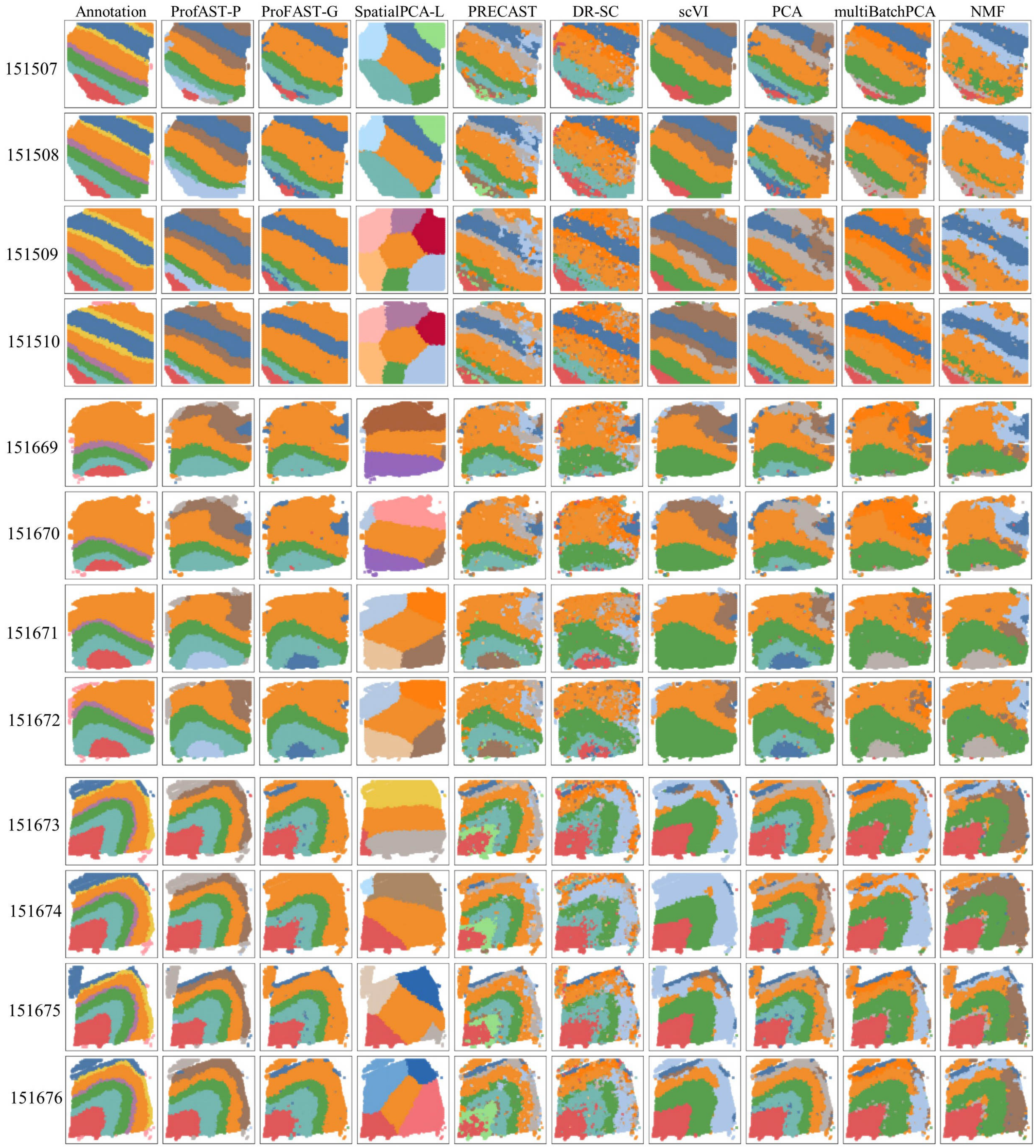


Figure S3: Clustering analysis for the 12 dorsolateral prefrontal cortex Visium sections. Clustering assignment heatmaps of the 12 sections for ProFAST, other methods, and manual annotations, with iSC-MEB used for clustering based on the low-dimensional embeddings from ProFAST and other compared methods.

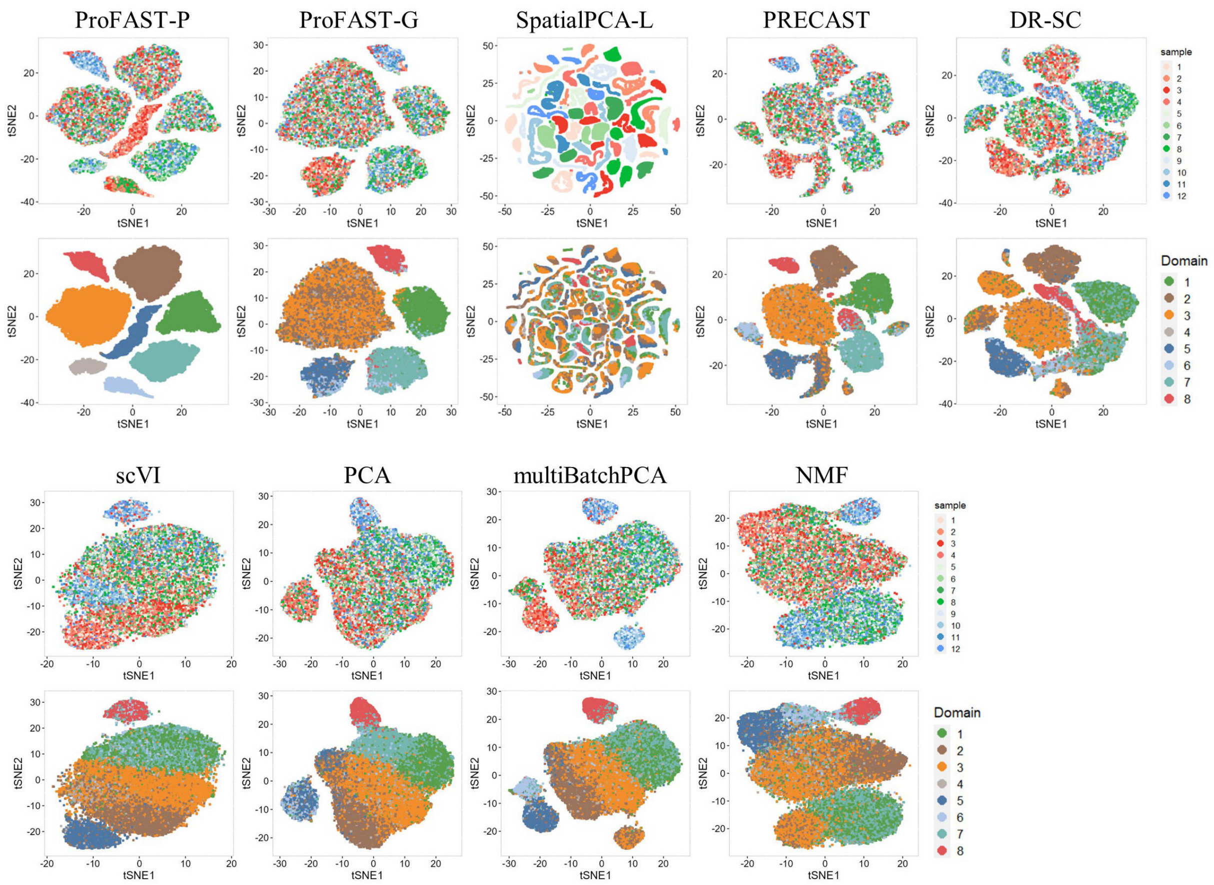


Figure S4: Comparison of integration performance for the 12 dorsolateral prefrontal cortex Visium sections. tSNE plots for the aligned embeddings via iSC-MEB based on the low-dimensional embeddings from ProFAST and other compared methods.

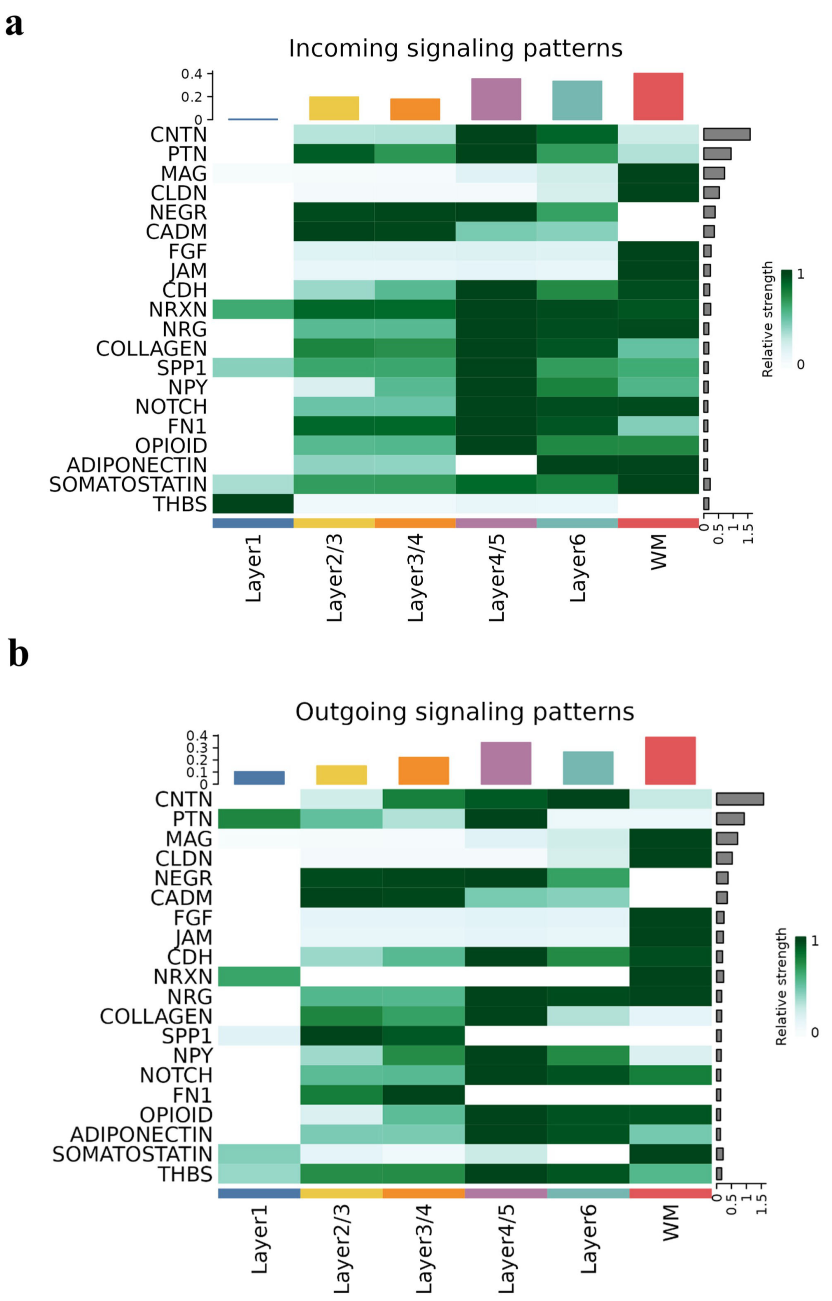


Figure S5: Cell-cell interaction analysis for the 12 dorsolateral prefrontal cortex Visium sections. Heatmaps of the incoming and outgoing signaling pathway patterns among the layers detected by ProFAST-P.

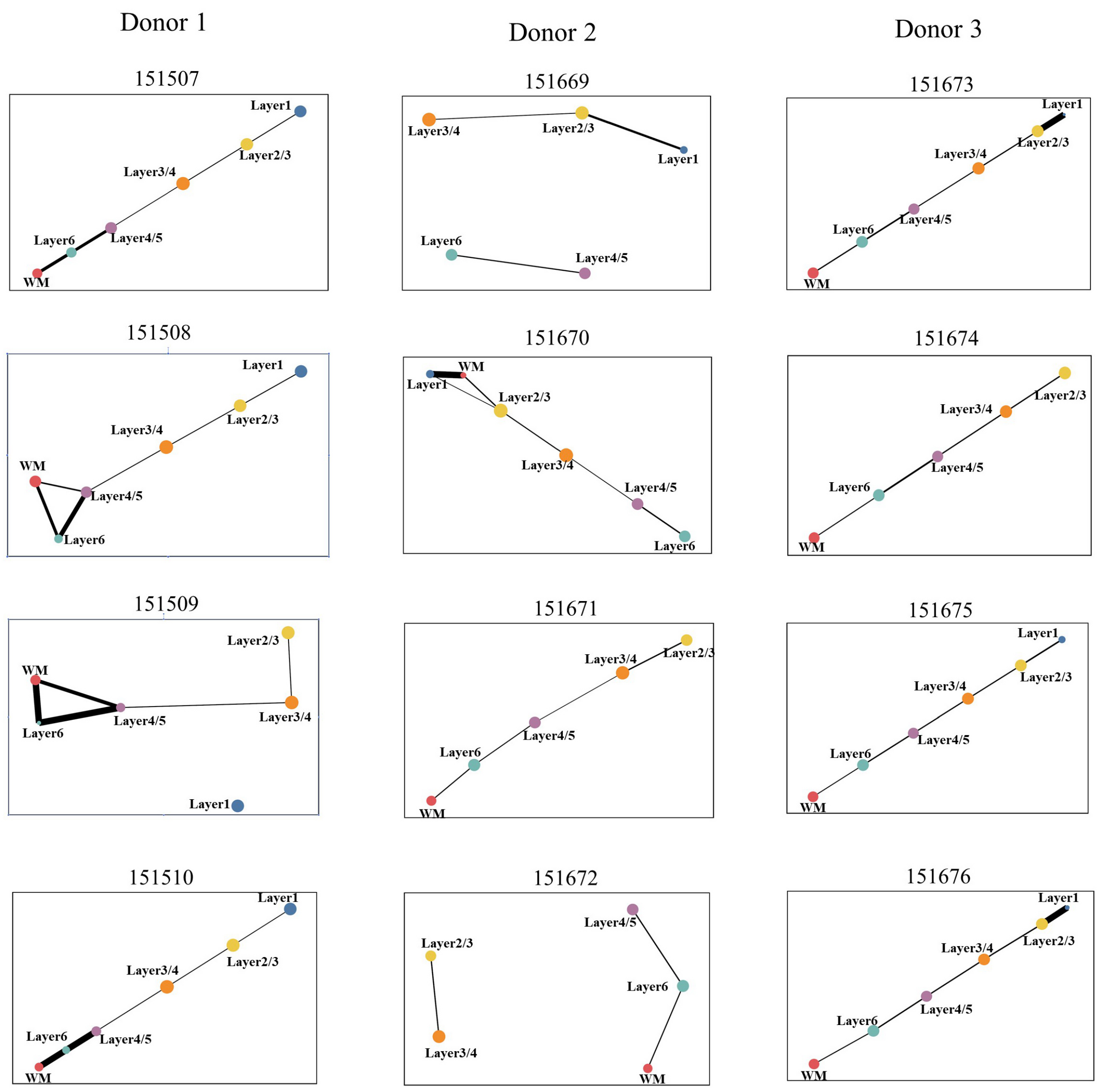


Figure S6: PAGA trajectory for the 12 dorsolateral prefrontal cortex Visium sections. PAGA trajectory for the 12 sections using the embeddings and layers label obtained by ProFAST-P.

##### 4.3 Application to breast cancer Xenium dataset

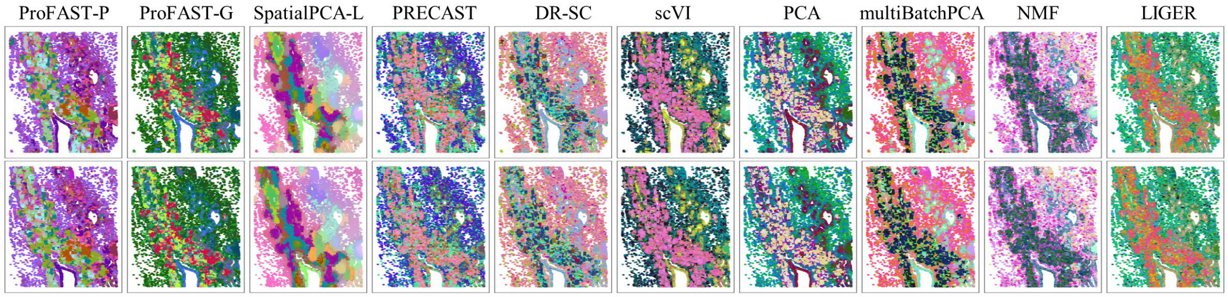


Figure S7: Dimension reduction with batch correction for the two human breast cancer Xenium sections. UMAP RGB plots for the two sections based on the aligned embeddings via iSC-MEB using the low-dimensional embeddings from ProFAST and other methods.

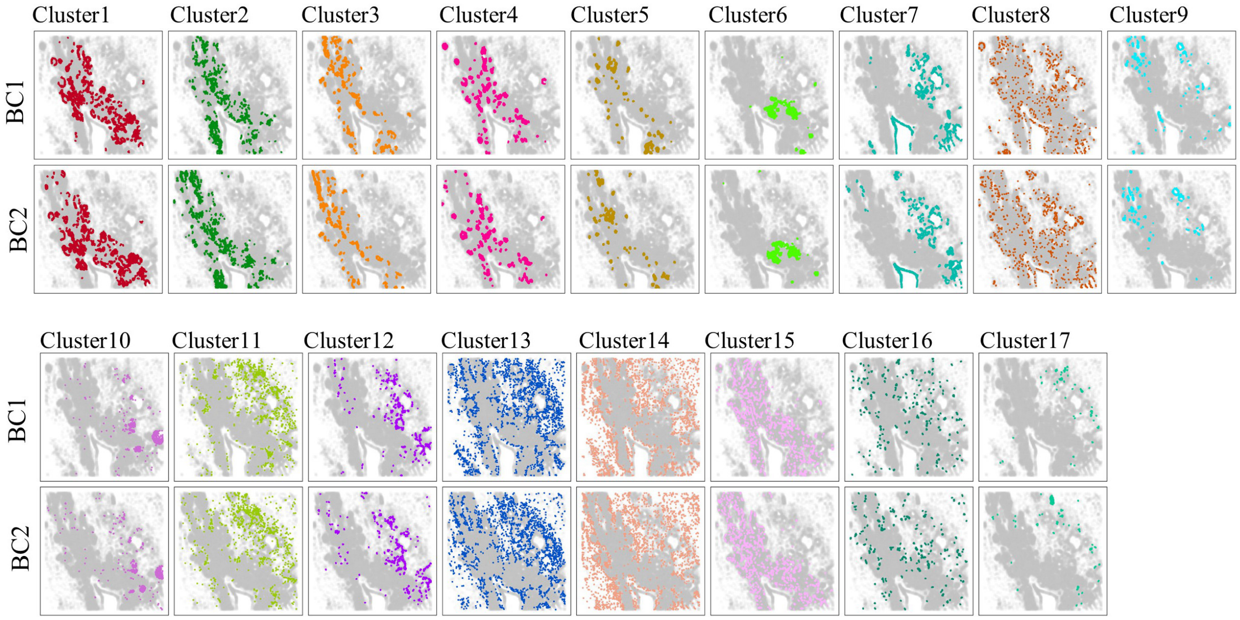


Figure S8: Spatial clustering analysis of data for the two human breast cancer Xenium sections. Clustering assignment heatmaps of the two sections for the 17 clusters via iSC-MEB based on the low-dimensional embeddings obtained by ProFAST-P.

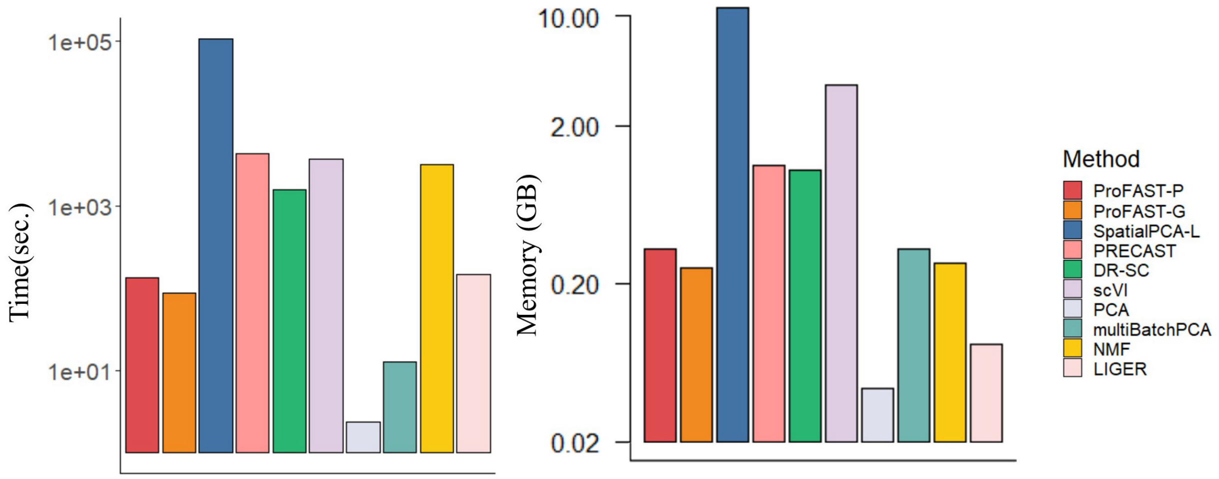


Figure S9: Comparison of computational cost for the two human breast cancer Xenium sections. Bar plots of running time (seconds) and memory usage (GB) for ProFAST and other methods. Note that a logarithmic scale, specifically $\log_{10}$, is employed for the y-axis.

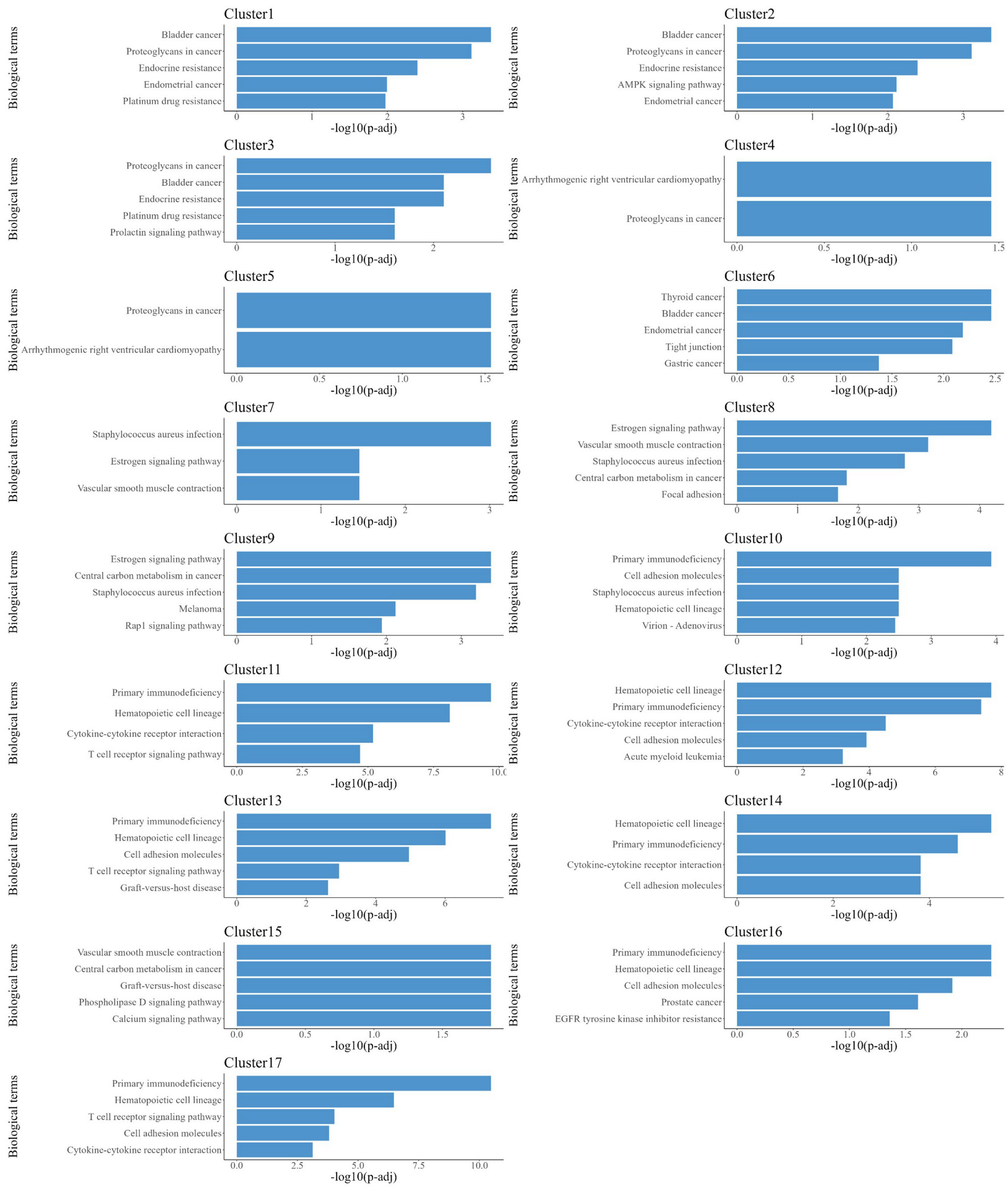


Figure S10: KEGG enrichment analysis of the marker genes of the 17 clusters detected by ProFAST-P from the two human breast cancer Xenium sections. Bar plot of -log10(adjusted $p$-value) for top five KEGG pathways in enrichment analysis. The $p$-values are based on one-sided Fisher's exact tests with multiple testing corrections using the Benjamini-Hochberg FDR method.

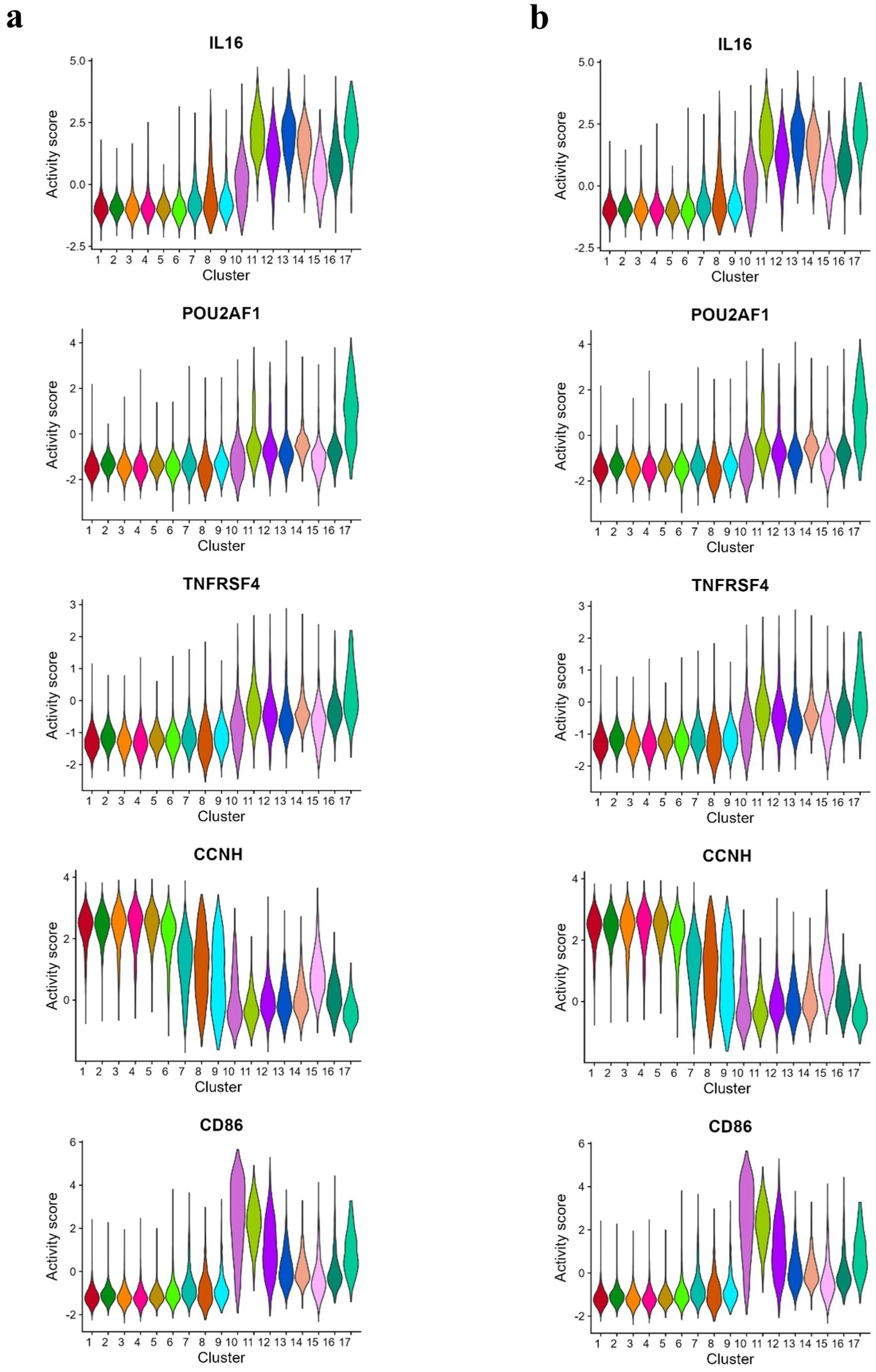


Figure S11: Transcription factor analysis for the two human breast cancer Xenium sections. Violin plots of the activity scores for the detected five regulatory proteins in the 17 clusters identified by ProFAST-P for each section. (a) for section 1; (b) for section 2.

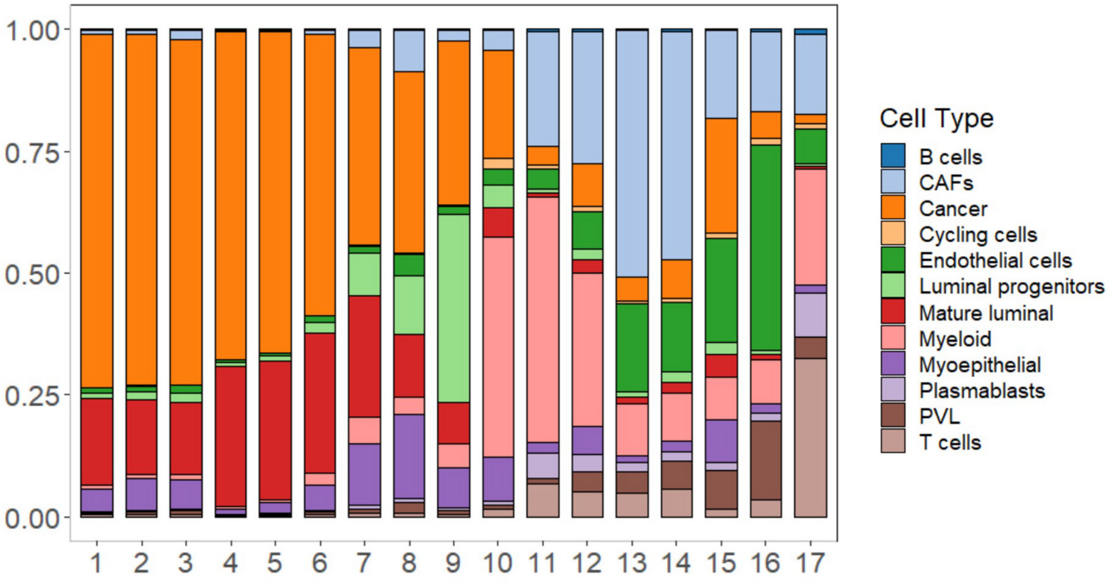


Figure S12: Cell-type deconvolution analysis for the two human breast cancer Xenium sections. Percentage of different cell types in each domain detected by ProFAST-P with scaling to the summation of all cell types in each domain equal to 100%.

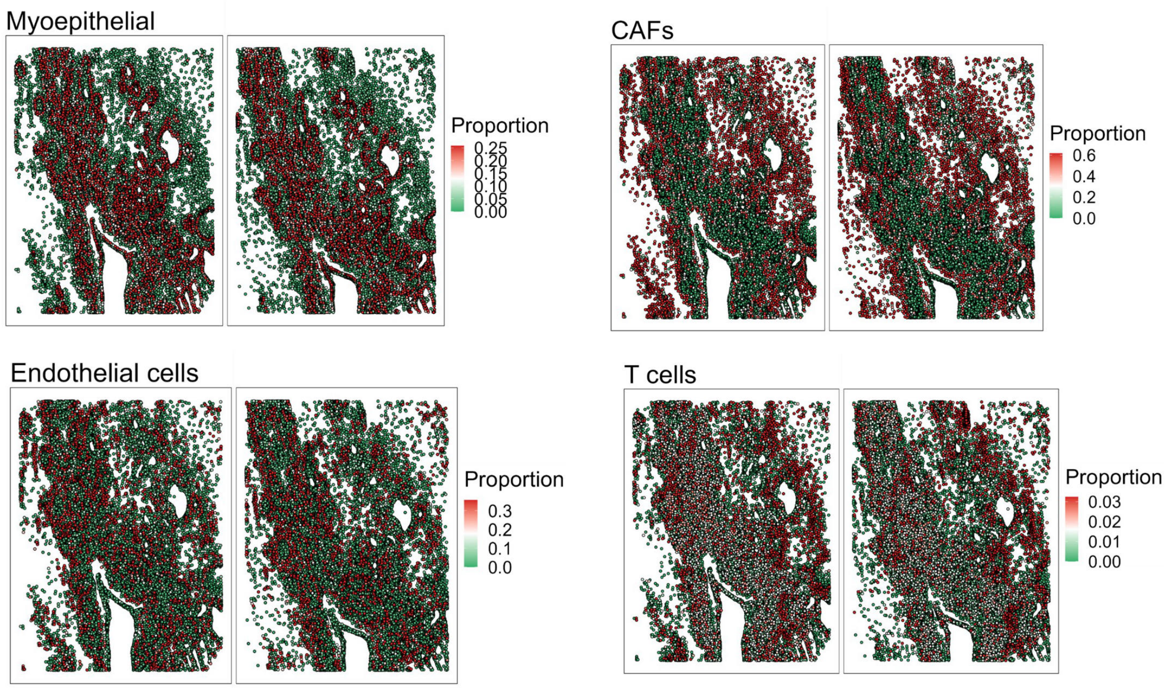


Figure S13: Cell-type deconvolution analysis for the two human breast cancer Xenium sections. Spatial heatmaps of proportions of four cell types (myoepithelial, CAFs: cancer-associated fibroblasts, endothelial cells, and T cells) annotated in the reference panel for the two sections.

#### 4.4 Application to hepatocellular carcinoma Visium dataset

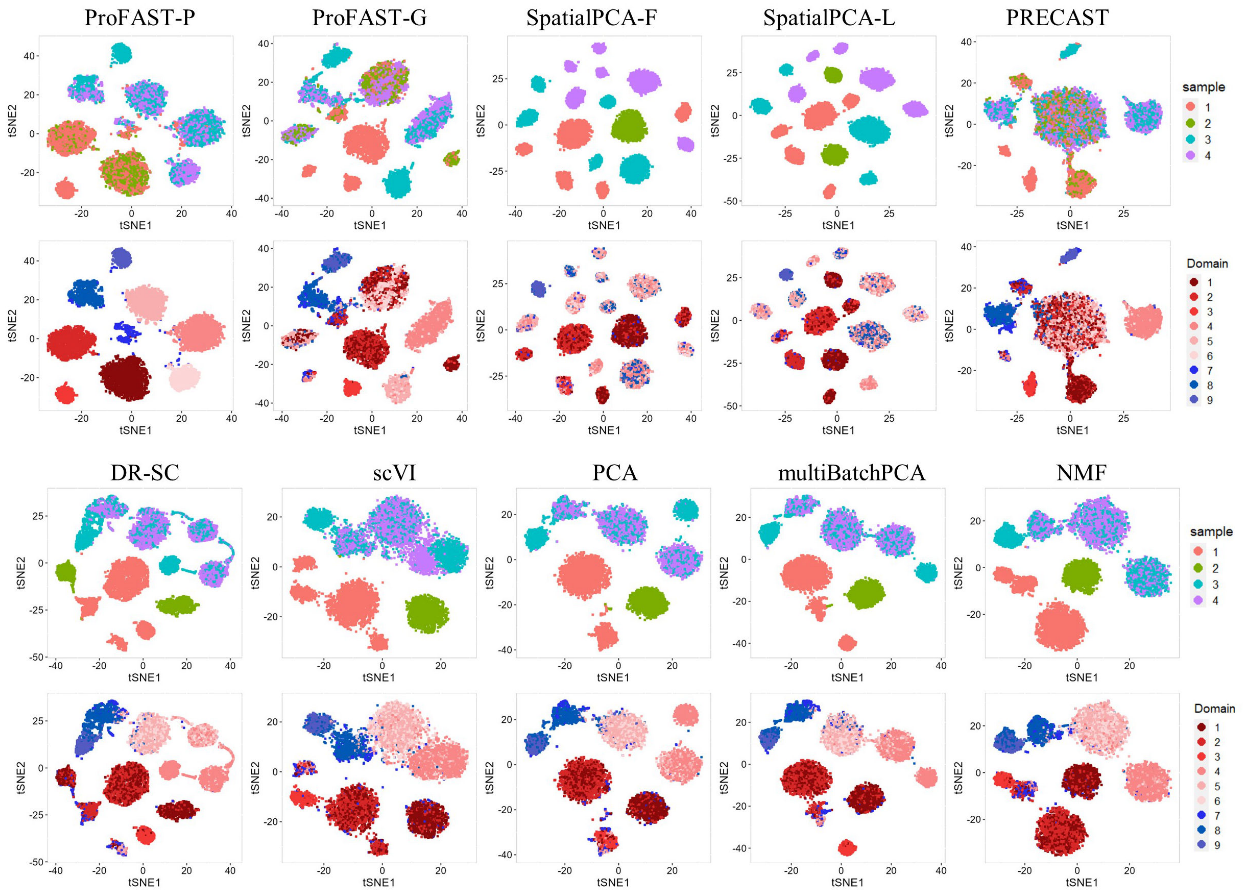


Figure S14: Comparison of integration performance for the four HCC Visium sections. tSNE plots for the aligned embeddings via iSC-MEB based on the low-dimensional embeddings from ProFAST and other compared methods.

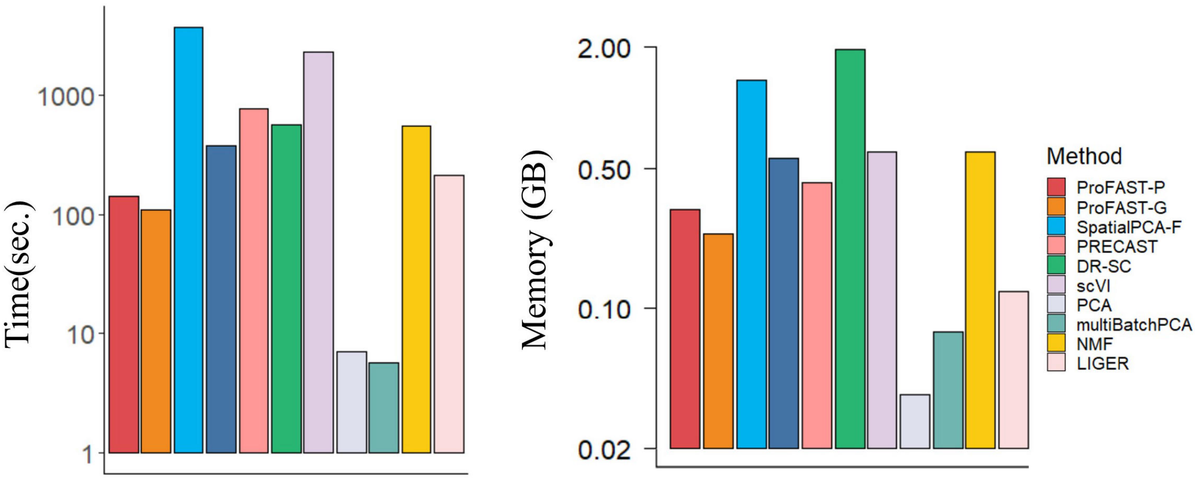


Figure S15: Comparison of computational cost for the four HCC Visium sections. Bar plots of running time (seconds) and memory usage (GB) for ProFAST and other methods. Note that a logarithmic scale, specifically $\log_{10}$, was employed for the y-axis.

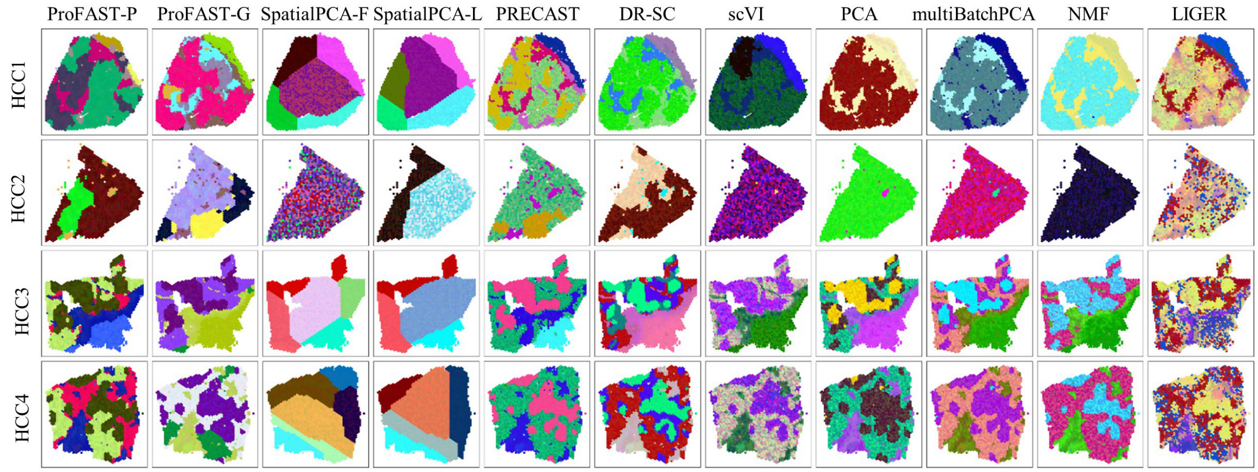


Figure S16: Dimension reduction with embedding alignment for the four HCC Visium sections. UMAP RGB plots for the four sections based on the embeddings via iSC-MEB using the low-dimensional embeddings from ProFAST and other methods.

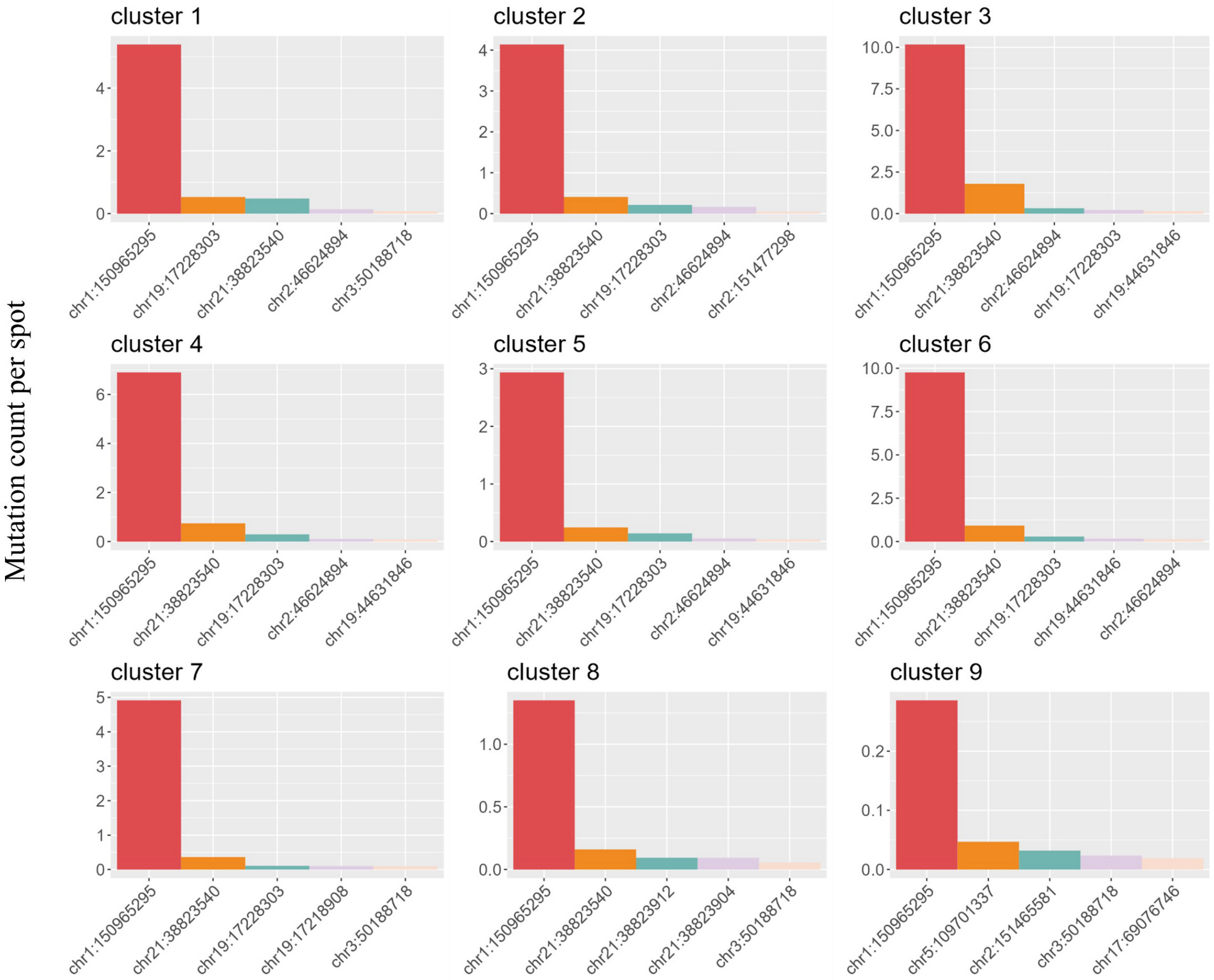


Figure S17: Somatic mutation analysis for the four human HCC sections. Bar plot of the mutation count of the top five SNPs in each cluster detected by ProFAST-P.

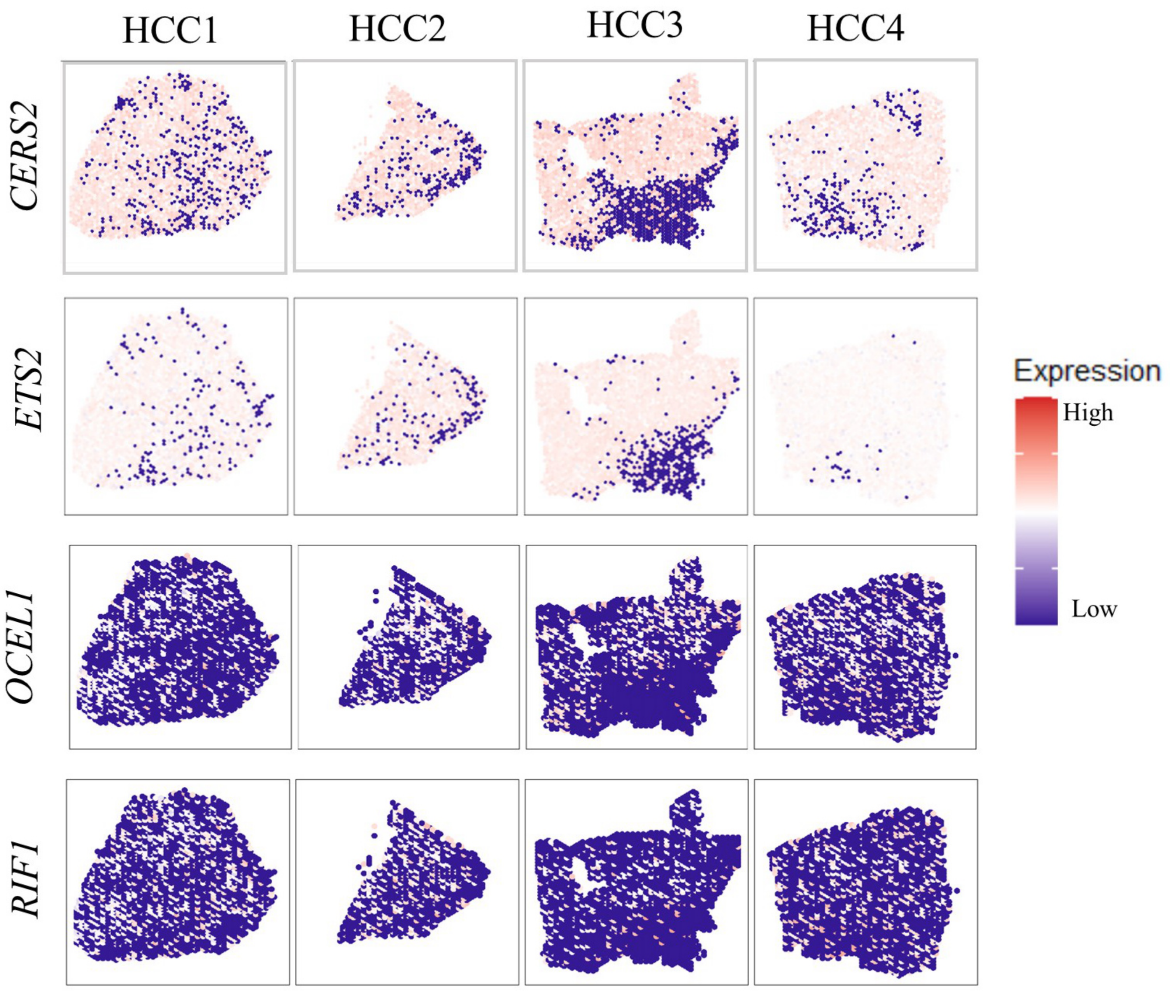


Figure S18: Somatic mutation analysis for the four human HCC sections. Spatial heatmap of the expressions of the four genes corresponding to the top four SNPs with somatic mutations for the four sections.

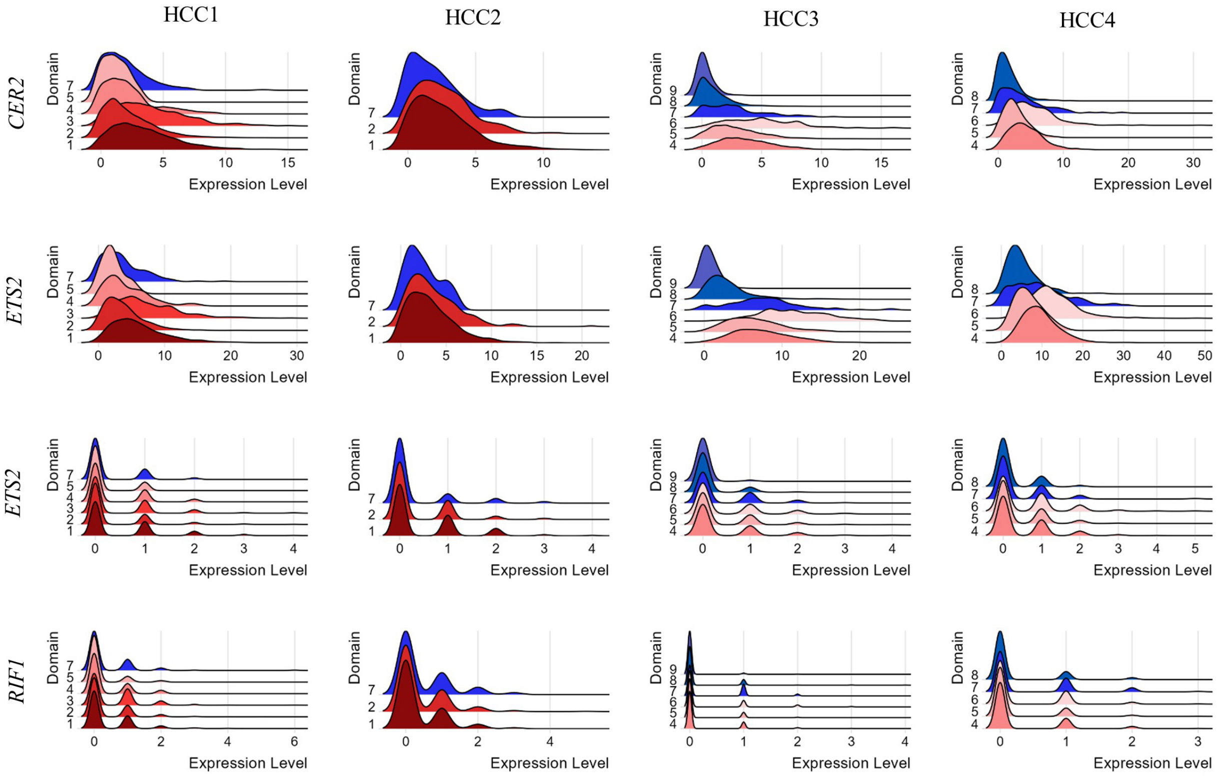


Figure S19: Somatic mutation analysis for the four human HCC sections. Ridge plots of the expressions of the four genes, corresponding to the top four SNPs with somatic mutations, for the domains identified by ProFAST-P in the four sections.

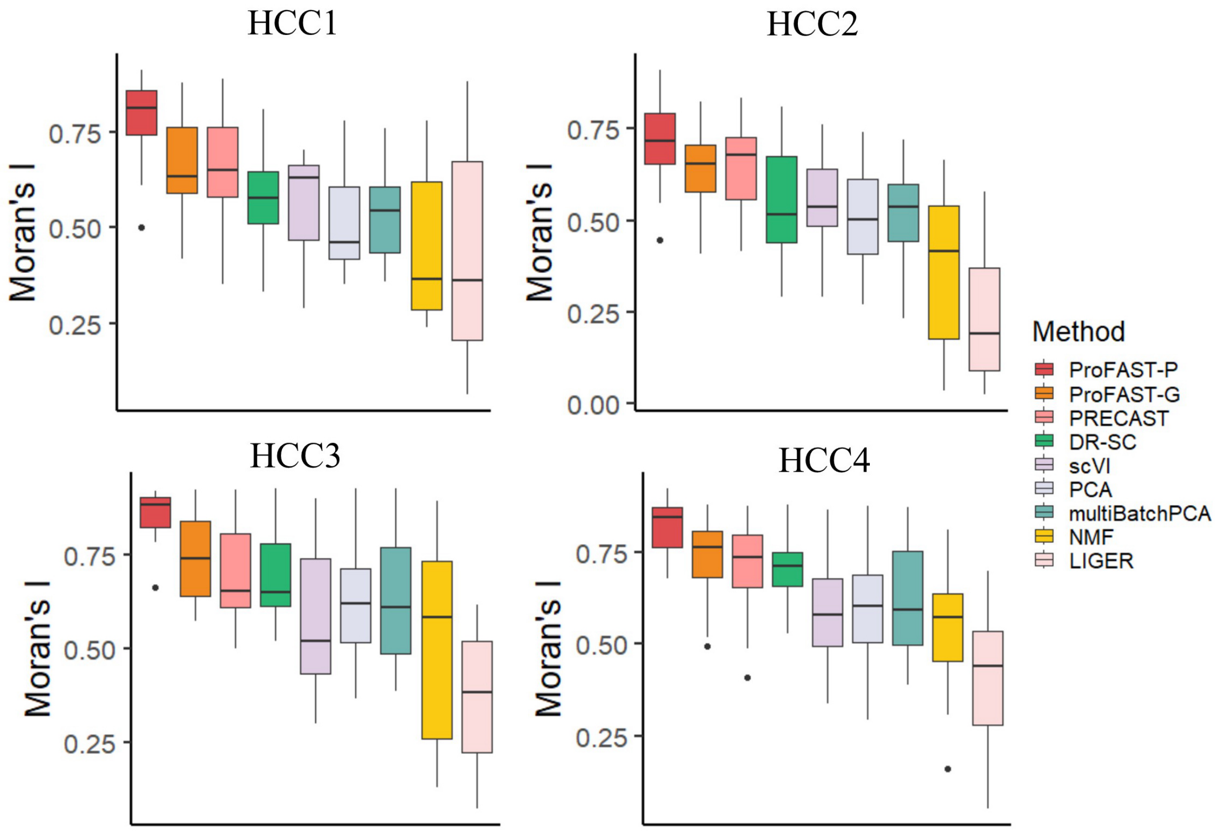


Figure S20: Spatial strength analysis for embeddings from different methods in the four human HCC sections. Boxplots of the Moran's I values for the 15-dimensional embeddings obtained by ProFAST and other methods. We did not plot the Moran's I values for SpatialPCA because SpatialPCA produced almost identical embedding for each spot and had a Moran's I value of 1. In the boxplots, the center line, box lines and whiskers represent the median, upper, and lower quartiles, and 1.5-times interquartile range, respectively.

##### 4.5 Application to mouse embryo Stereo-seq dataset

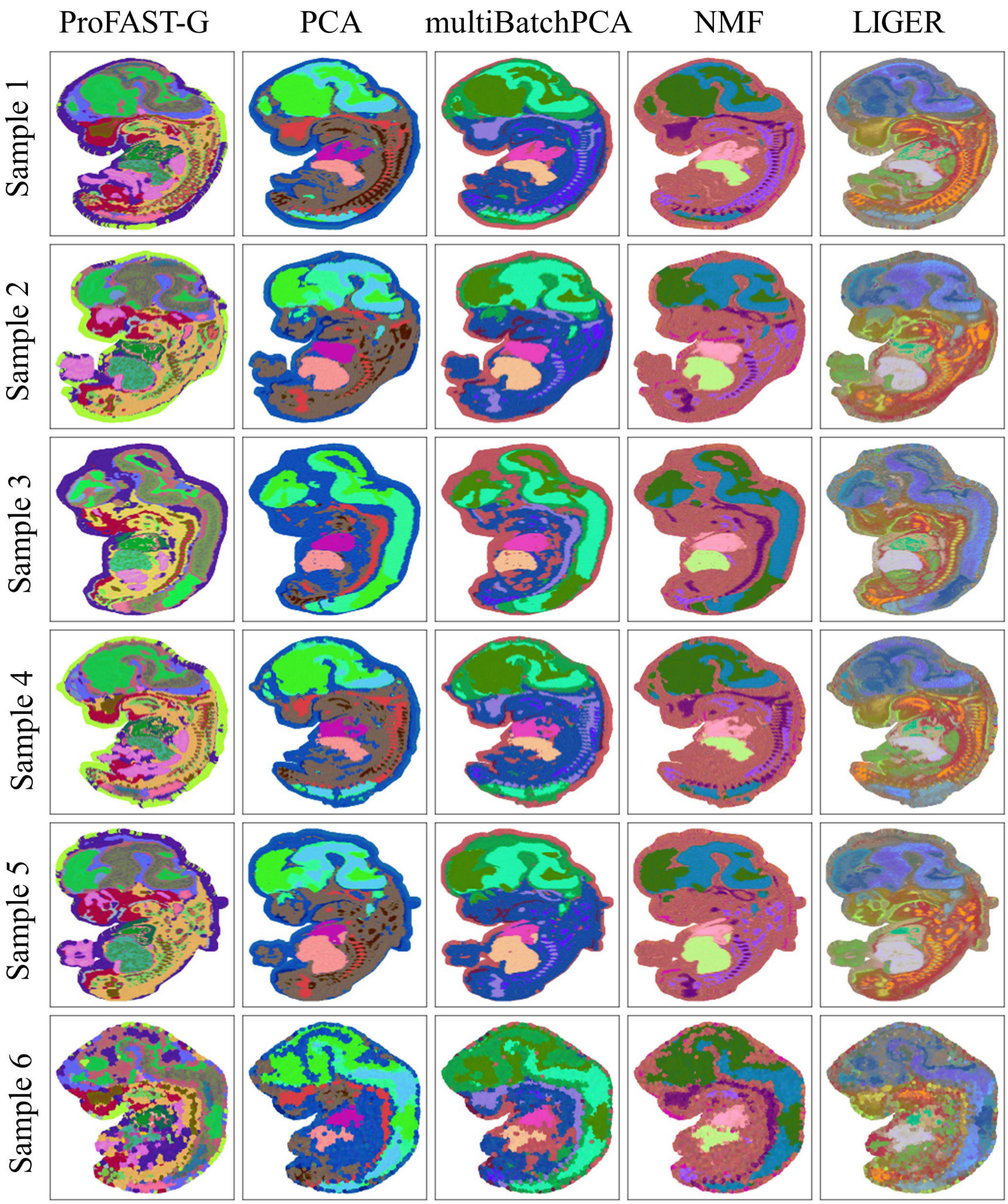


Figure S21: Dimension reduction with embedding alignment for the 26 mouse embryo Stereo-seq sections. UMAP RGB plots for the six sections in the embryo day E12.5 based on the embeddings obtained by iSC-MEB using the low-dimensional embeddings from ProFAST and other methods.

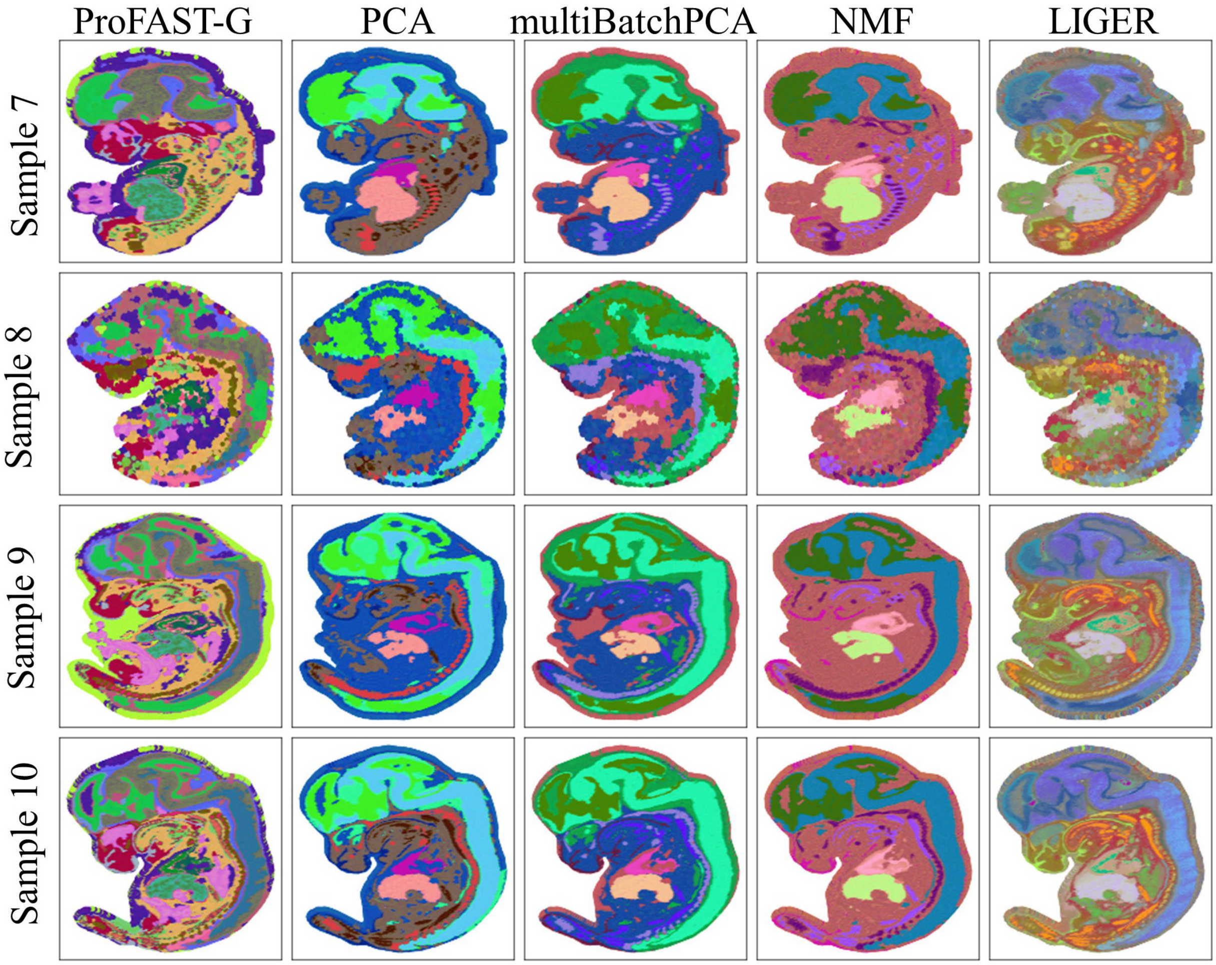


Figure S22: Dimension reduction with embedding alignment for the 26 mouse embryo Stereo-seq sections. UMAP RGB plots for the four sections in the embryo day E13.5 based on the embeddings obtained by iSC-MEB using the low-dimensional embeddings from ProFAST and other methods.

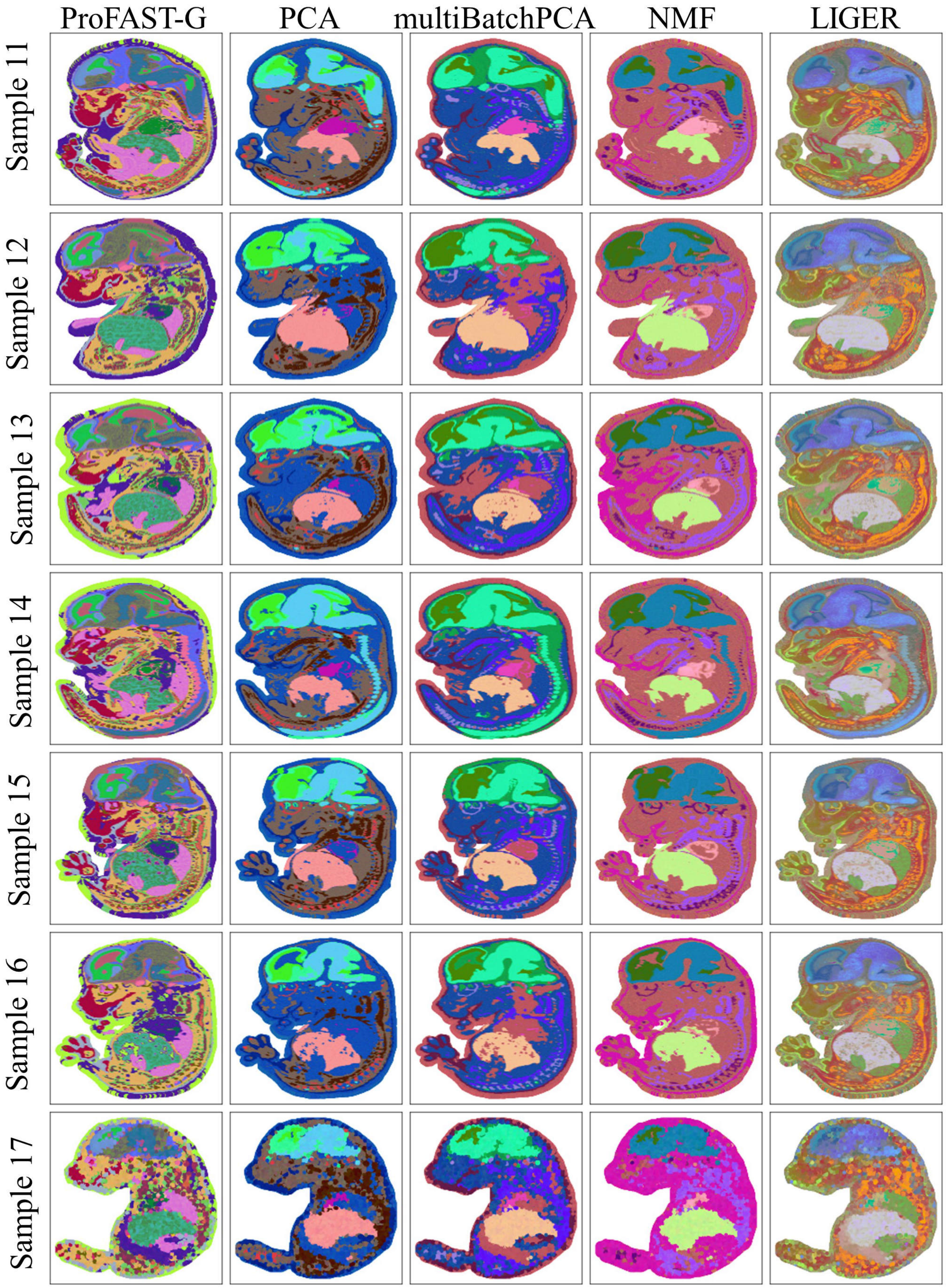


Figure S23: Dimension reduction with embedding alignment for the 26 mouse embryo Stereo-seq sections. UMAP RGB plots for the seven sections in the embryo day E14.5 based on the embeddings obtained by iSC-MEB using the low-dimensional embeddings from ProFAST and other methods.

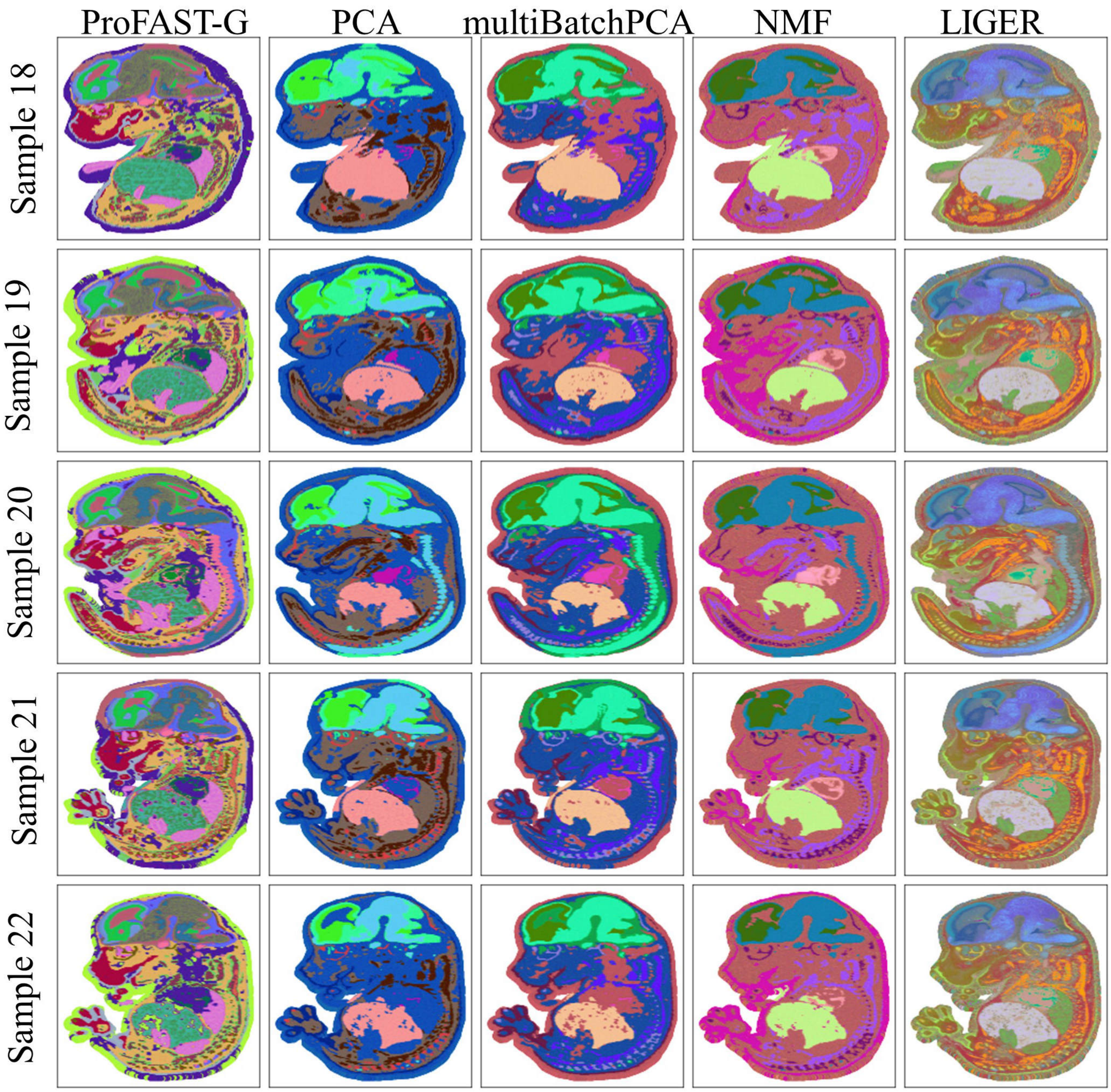


Figure S24: Dimension reduction with embedding alignment for the 26 mouse embryo Stereo-seq sections. UMAP RGB plots for the five sections in the embryo day E15.5 based on the embeddings obtained by iSC-MEB using the low-dimensional embeddings from ProFAST and other methods.

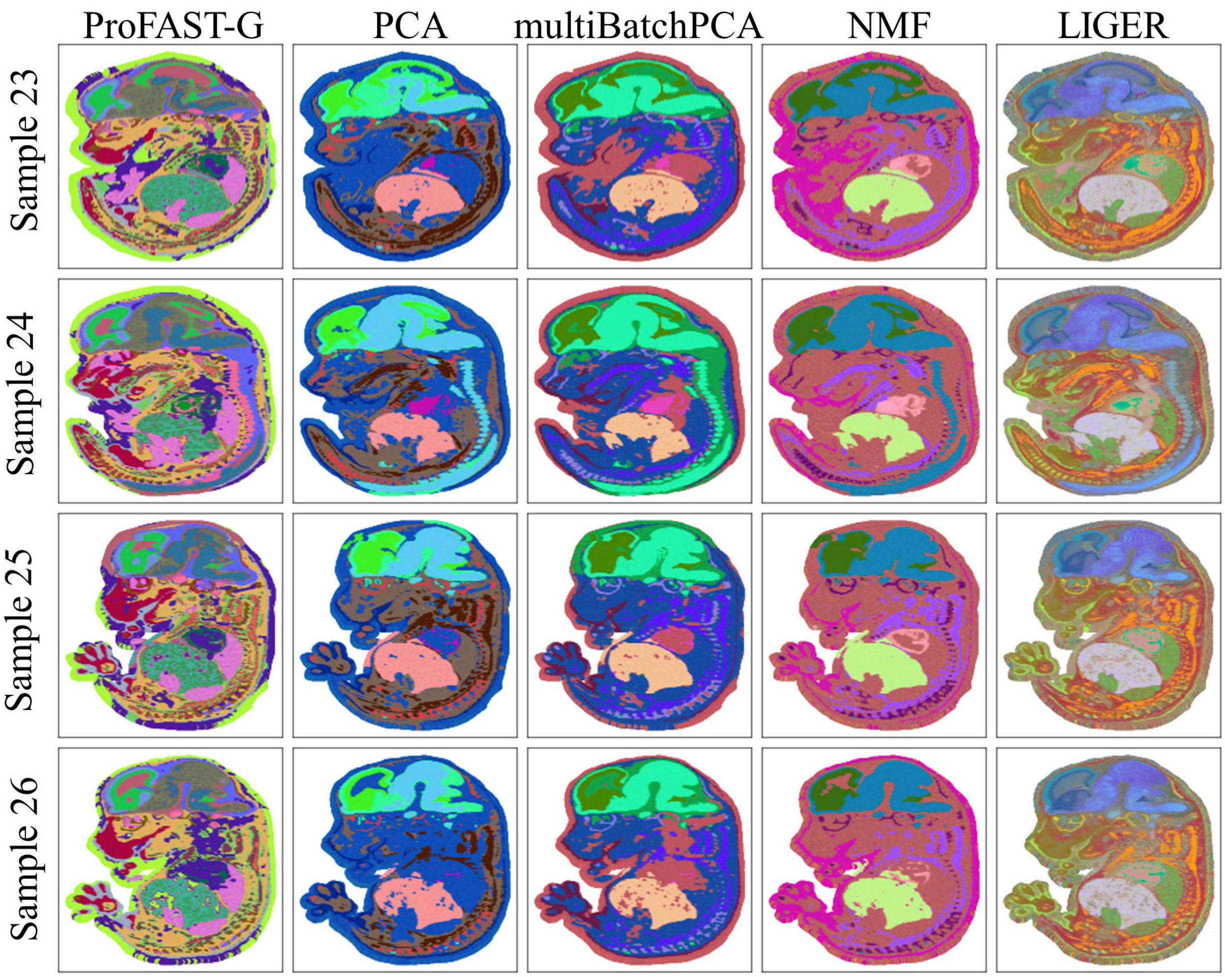


Figure S25: Dimension reduction with embedding alignment for the 26 mouse embryo Stereo-seq sections. UMAP RGB plots for the four sections in the embryo day E16.5 based on the embeddings obtained by iSC-MEB using the low-dimensional embeddings from ProFAST and other methods.

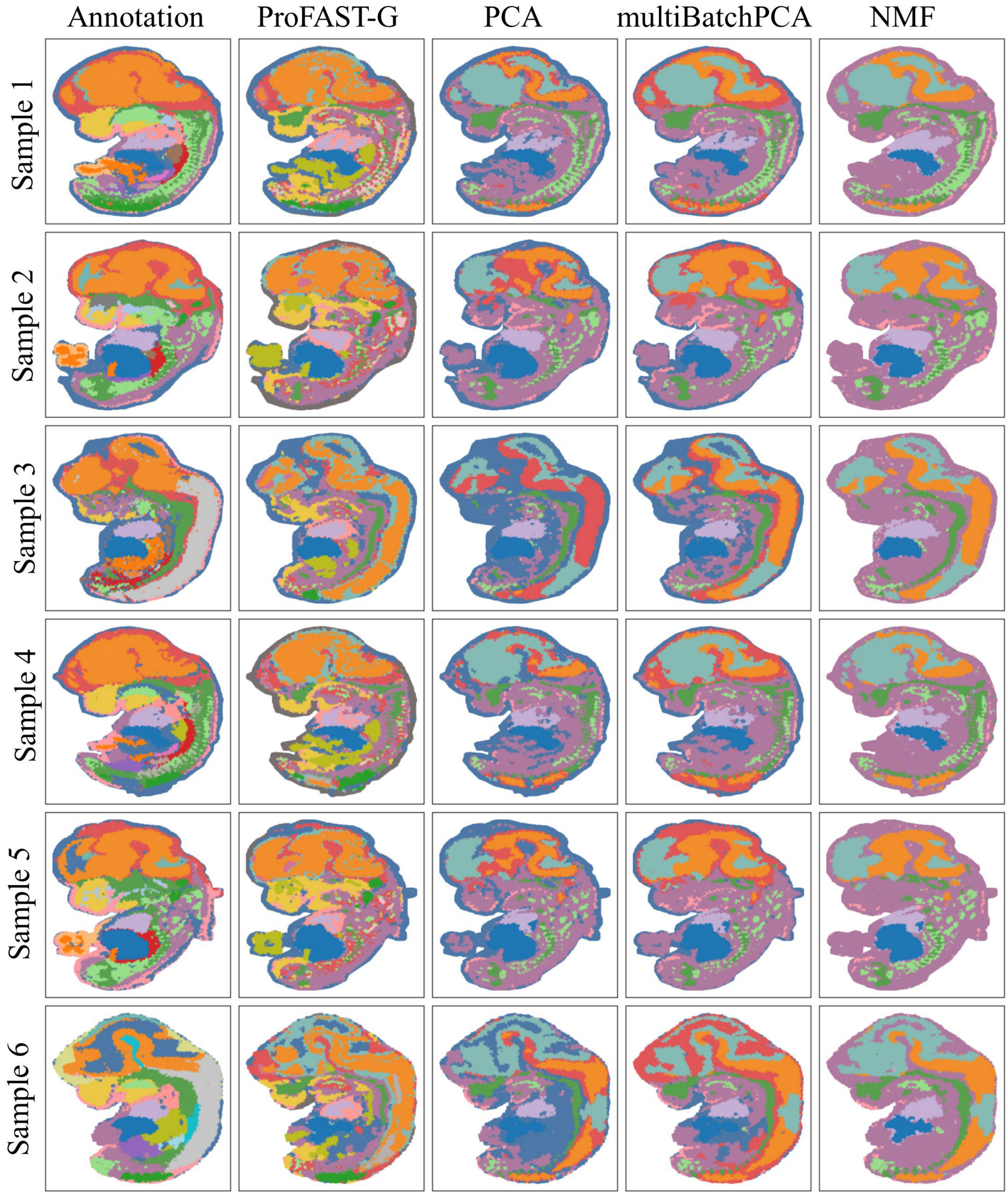


Figure S26: Clustering analysis for the 26 mouse embryo Stereo-seq sections. Clustering assignment heatmaps of the six sections in the embryo day E12.5 for ProFAST, other methods, and manual annotations, with iSC-MEB used for clustering based on the low-dimensional embeddings from ProFAST and other compared methods.

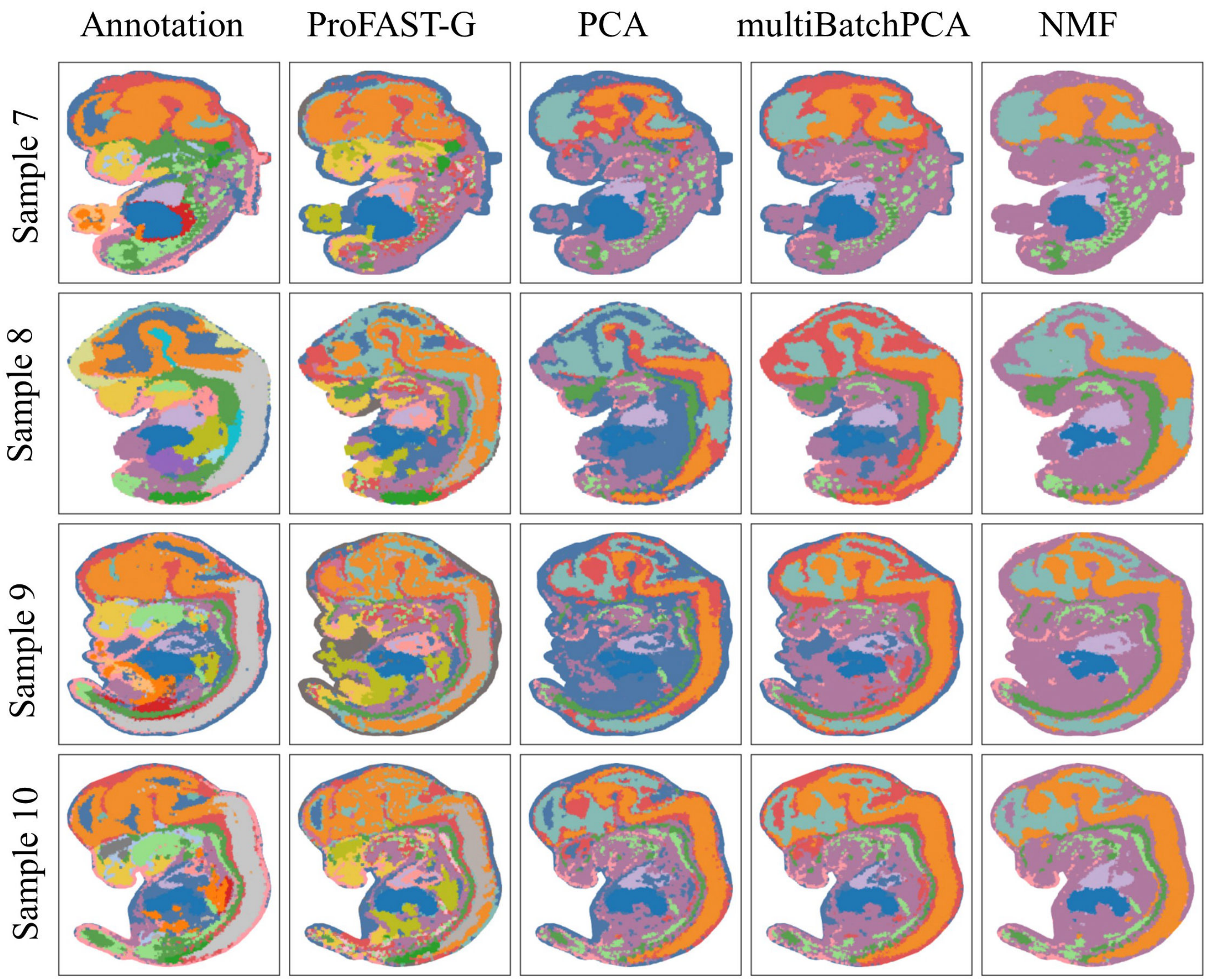


Figure S27: Clustering analysis for the 26 mouse embryo Stereo-seq sections. Clustering assignment heatmaps of the four sections in the embryo day E13.5 for ProFAST, other methods, and manual annotations, with iSC-MEB used for clustering based on the low-dimensional embeddings from ProFAST and other compared methods.

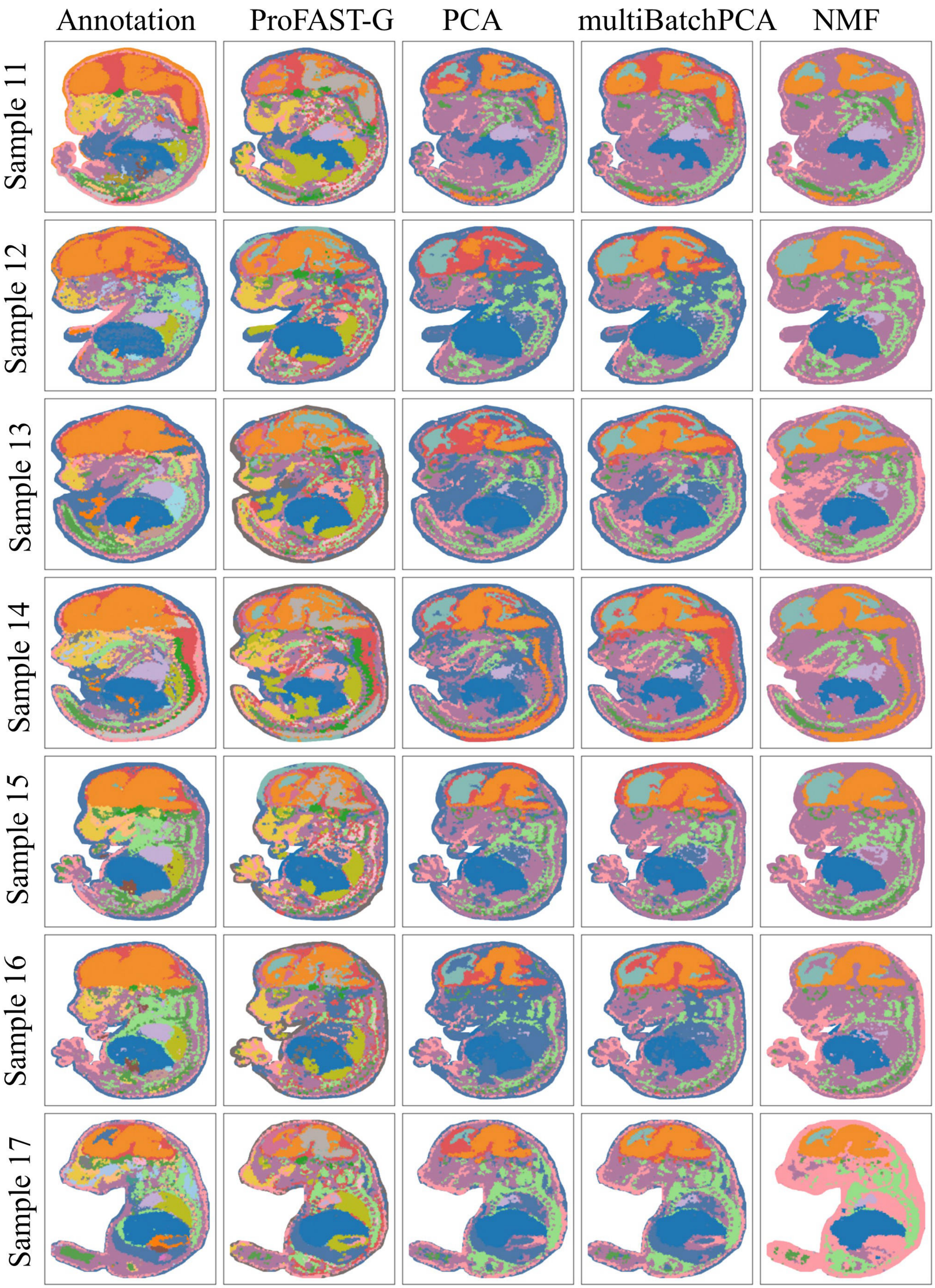


Figure S28: Clustering analysis for the 26 mouse embryo Stereo-seq sections. Clustering assignment heatmaps of the seven sections in the embryo day E14.5 for ProFAST, other methods, and manual annotations, with iSC-MEB used for clustering based on the low-dimensional embeddings from ProFAST and other compared methods.

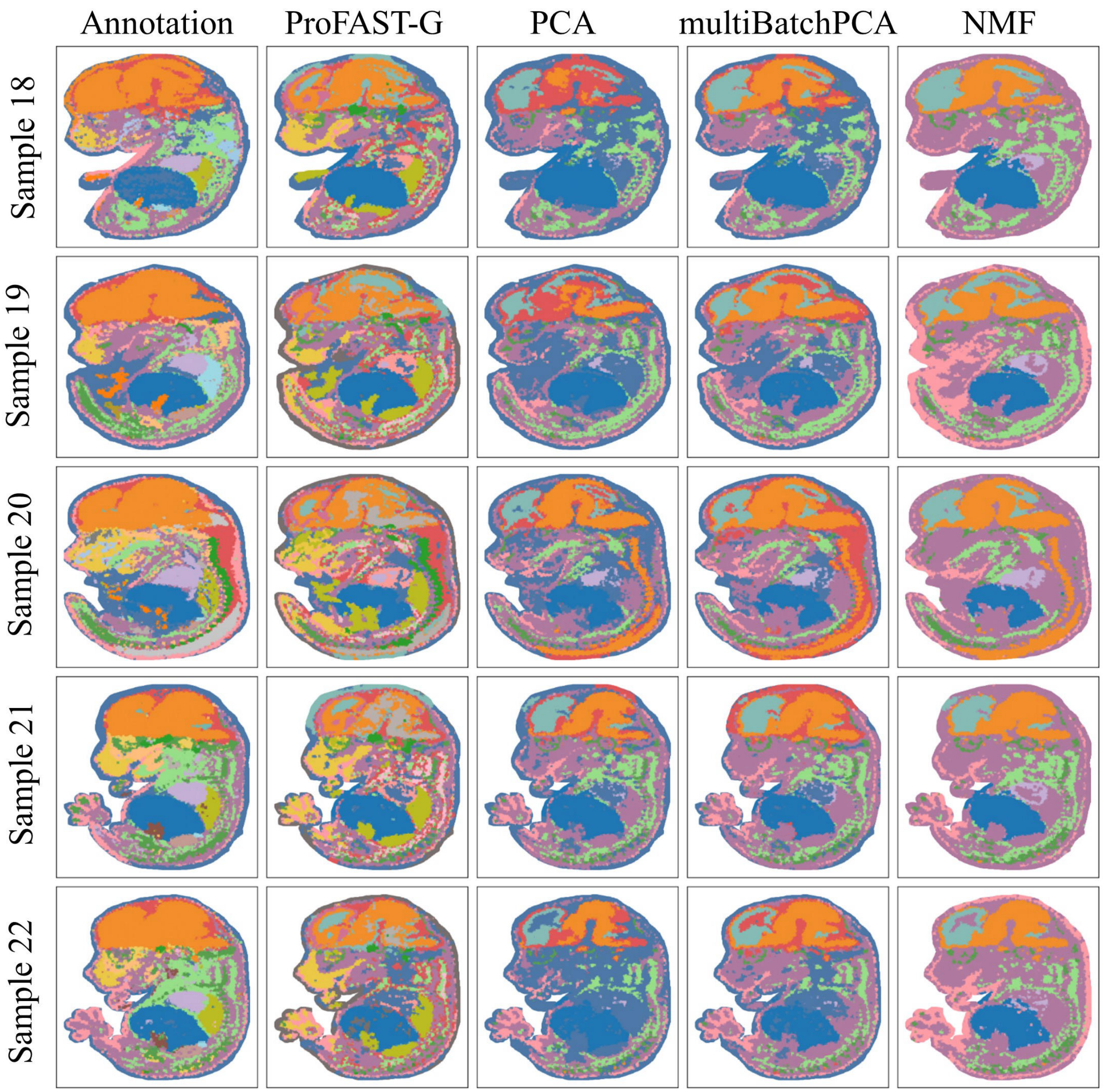


Figure S29: Clustering analysis for the 26 mouse embryo Stereo-seq sections. Clustering assignment heatmaps of the five sections in the embryo day E15.5 for ProFAST, other methods, and manual annotations, with iSC-MEB used for clustering based on the low-dimensional embeddings from ProFAST and other compared methods.

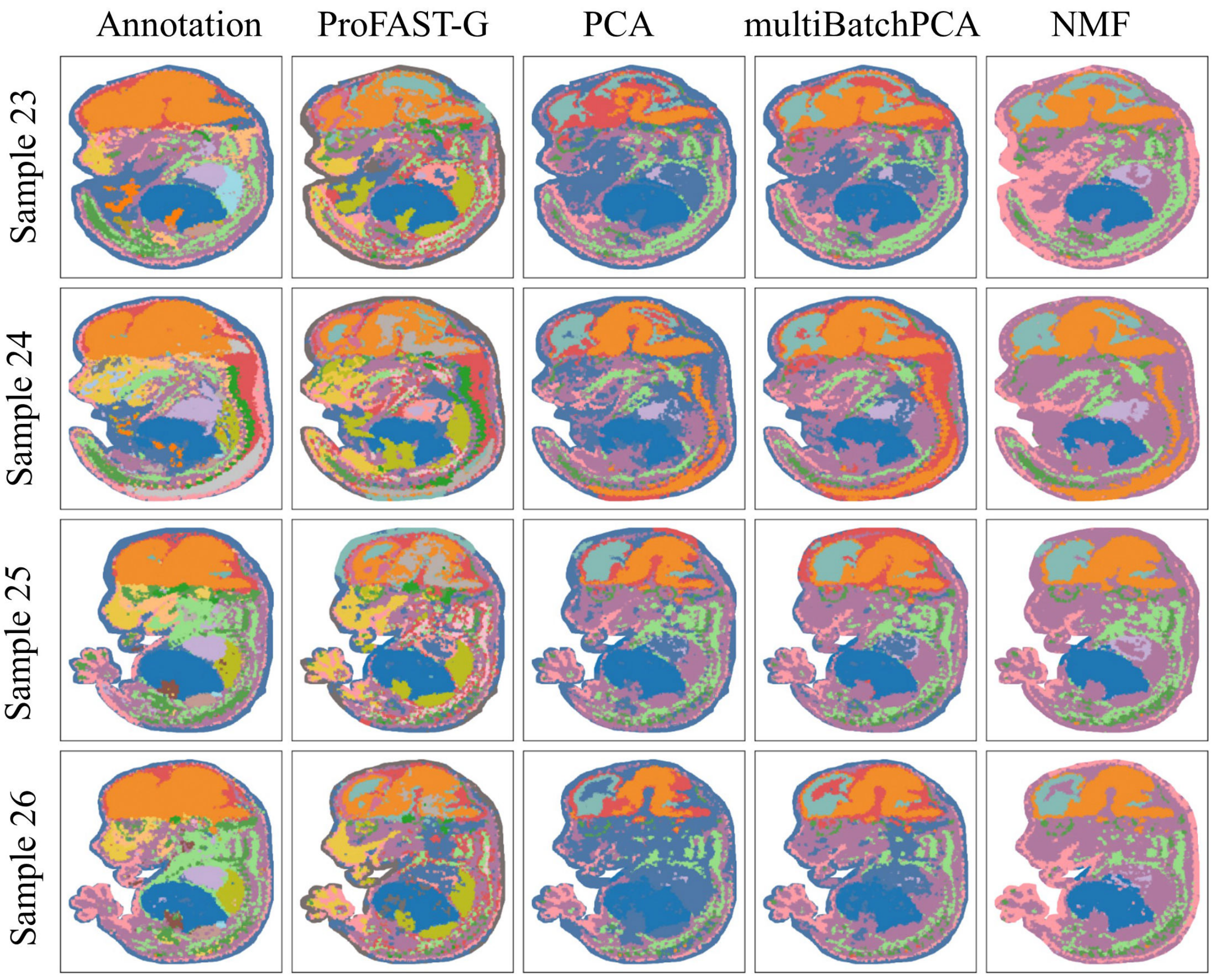


Figure S30: Clustering analysis for the 26 mouse embryo Stereo-seq sections. Clustering assignment heatmaps of the four sections in the embryo day E16.5 for ProFAST, other methods, and manual annotations, with iSC-MEB used for clustering based on the low-dimensional embeddings from ProFAST and other compared methods.

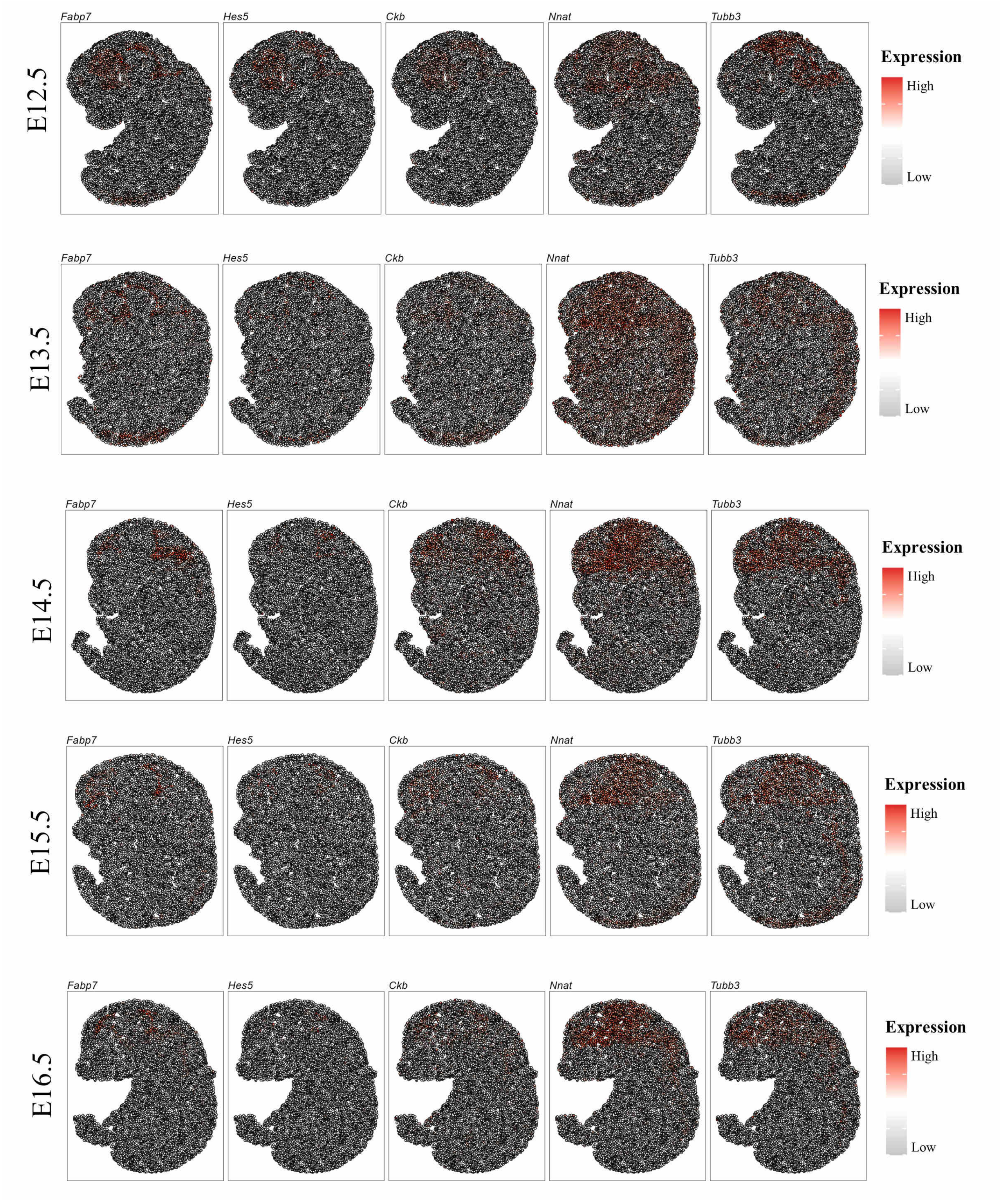


Figure S31: Combined DE analysis of data for the 26 mouse embryo Stereo-seq sections. Spatial heatmaps of top marker genes specific to the brain region for sections from different embryo days (the first sample in E12.5, E13.5, E14.5, E15.5 and E16.5, respectively).

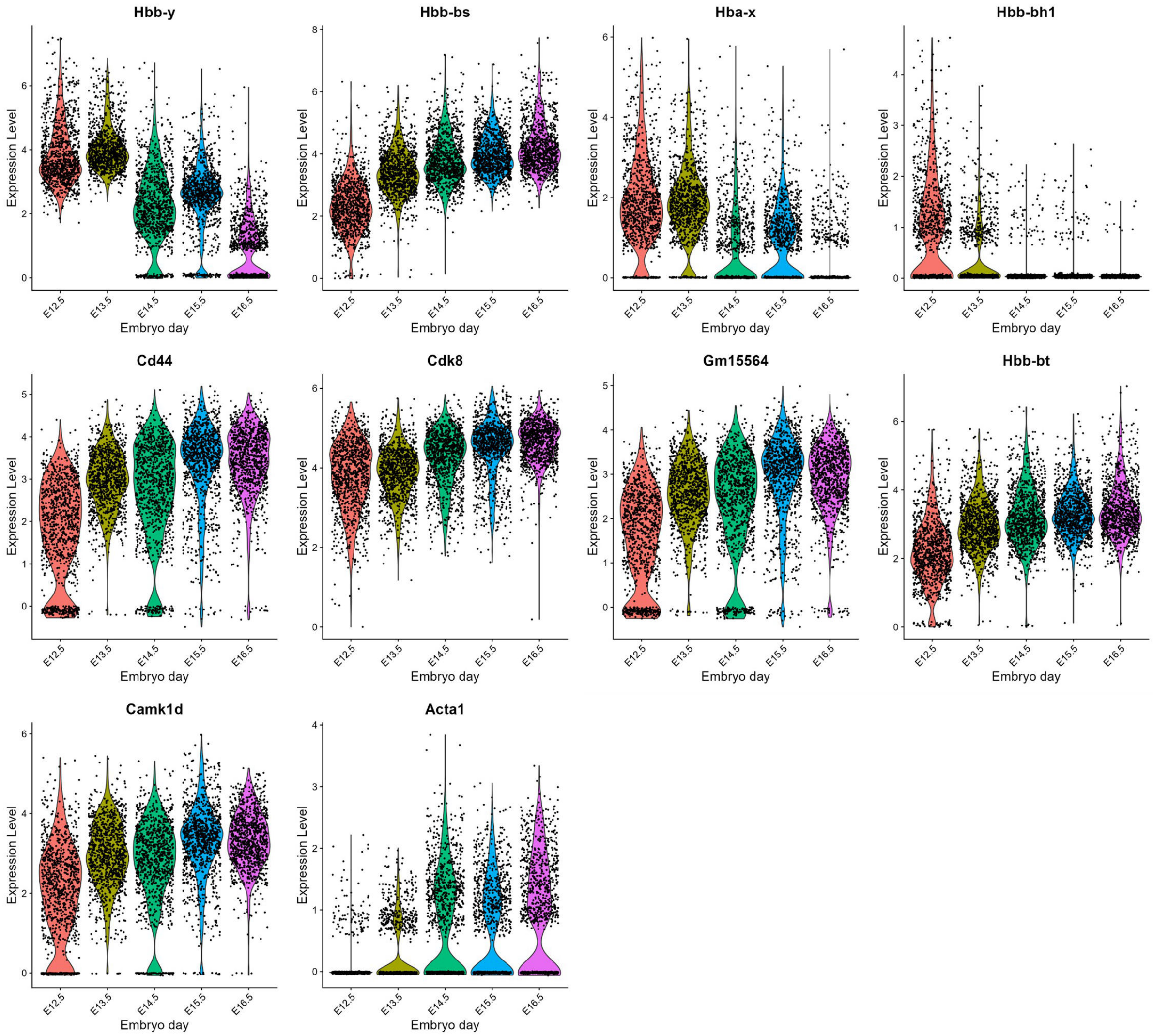


Figure S32: Detection of temporal trend of gene expression levels in the brain region for the 26 mouse embryo Stereo-seq sections. Violin plots for the expression levels of the top 10 genes.
